# *HOTAIR* primes the Ewing sarcoma family of tumors for tumorigenesis via epigenetic dysregulation involving LSD1

**DOI:** 10.1101/244558

**Authors:** Hasan Siddiqui, Julia Selich-Anderson, Joshua Felgenhauer, James Fitch, Vijay Nadella, Cenny Taslim, Laura Tomino, Emily Theisen, Satoru Otsuru, Edwin Horwitz, Stephen Lessnick, Peter White, Nilay Shah

**Affiliations:** Center for Childhood Cancer and Blood Diseases, The Research Institute at Nationwide Children’s Hospital, 700 Children’s Drive, Columbus, OH, 43205; Institute for Genomic Medicine, The Research Institute at Nationwide Children’s Hospital, 700 Children’s Drive, Columbus, OH, 43205; Biomedical Genomics Core, The Research Institute at Nationwide Children’s Hospital, 700 Children’s Drive, Columbus, OH, 43205; The Ohio State University College of Medicine, 370 W 9^th^ Ave, Columbus, OH, 43210

**Keywords:** Ewing sarcoma, Epigenetics, HOTAIR, long noncoding RNA, pediatrics

## Abstract

The EWS-FLI1 fusion protein drives oncogenesis in the Ewing sarcoma family of tumors (ESFT) in humans, but its toxicity in normal cells requires additional cellular events for oncogenesis. We show that the lncRNA *HOTAIR* maintains cell viability in the presence of EWS-FLI1 and redirects epigenetic regulation in ESFT. *HOTAIR* is consistently overexpressed in ESFTs and is not driven by EWS-FLI1. Repression of *HOTAIR* in ESFT cell lines significantly reduces anchorage-independent colony formation in vitro and impairs tumor xenograft growth in vivo. Overexpression of *HOTAIR* in human mesenchymal stem cells (hMSCs), a putative cell of origin of ESFT, and IMR90 cells induces colony formation. Critically, HOTAIR-expressing hMSCs and IMR90 cells remain viable with subsequent *EWS-FLI1* expression. *HOTAIR* induces histone modifications and gene repression through interaction with the epigenetic modifier LSD1 in ESFT cell lines and hTERT-hMSCs. Our findings suggest that *HOTAIR* maintains ESFT viability through epigenetic dysregulation.

**Significance:** While the *EWS-FLI1* fusion gene was determined to be the oncogenic driver in the overwhelming majority of ESFT, it is toxic to cell physiology and requires one or more additional molecular events to maintain cell viability. As these tumors have surprisingly few genetic mutations at diagnosis, epigenetic changes have been considered to be such an event, but the mechanism by which these changes are driven remains unclear. Our work shows that *HOTAIR* is consistently expressed among ESFT and induces epigenetic and gene expression changes that cooperate in tumorigenesis. Furthermore, expression of *HOTAIR* allows for cell viability in the setting of subsequent *EWS-FLI1* expression. Our findings elucidate new steps of malignant transformation in this cancer and identify novel therapeutic targets.

## INTRODUCTION

The Ewing sarcoma family of tumors (ESFT) consists of primitive cancers of the bone and soft tissues that arise in children and young adults. These tumors harbor chromosomal translocations that result in the fusion of the 5’ portion of the *EWSR1* gene to the 3’ end of an ETS family member, with >85% resulting in the *EWS-FLI1* fusion gene(1). The resultant oncoprotein alters gene expression and alternative splicing(2, 3) and is necessary for tumor viability in laboratory models of ESFT(4). However, exogenous expression of *EWS-FLI1* is toxic to normal cells and other cancer cell types, inducing rapid apoptosis or senescence(5). This toxicity suggests that additional cellular events must allow tolerance of the oncoprotein. Three large genomic sequencing studies identified some recurrent genetic mutations in ESFT, including loss-of-function mutations in *TP53, CDKN2A*, and *STAG2(6–8)*, but these mutations were found in only 25–30% of all samples. Thus, additional molecular changes, such as transcriptional or epigenetic events, must occur to allow tumor cell survival.

Long noncoding RNAs (lncRNAs) have significant roles in the regulation of gene expression, either directly or epigenetically. The lncRNA *HOTAIR* was specifically shown to direct epigenetic repression *in trans* across the genome, in part through recruitment of the LSD1/REST/CoREST complex at its 3’ end(9). This RNA-protein complex alters histone methylation at histone 3 lysine 4 (H3K4), demethylated by LSD1 from a dimethylated state (Me2) to mono- (Me) or unmethylated. This histone modification represses gene expression and maintains an embryonic state in tissues where *HOTAIR* is expressed. *HOTAIR* is abnormally overexpressed in numerous cancers (reviewed in (10)), and *HOTAIR* has been shown to epigenetically modify gene expression in these cancers (11–13).

In this study, we evaluated the function of *HOTAIR* in ESFT. We confirmed that *HOTAIR* is overexpressed in ESFT cell lines and primary tumors, as compared to normal tissues and to human mesenchymal stem cells (hMSCs), a putative cell of origin of these tumors(14, 15). We demonstrated that *HOTAIR* is necessary for the formation and viability of ESFT cell line-derived anchorage-independent colonies. We also showed that repression of *HOTAIR* by shRNA reduced tumor xenograft formation from ESFT cell lines in immunodeficient mice. In contrast, overexpression of *HOTAIR* in primary and hTERT-immortalized hMSCs induces anchorage-independent colony formation. *HOTAIR* expression in hMSCs and IMR90 fibroblasts also allows for subsequent viable expression of *EWS-FLI1*. We verified that *HOTAIR* associates with LSD1 in ESFT cell lines, and interaction with LSD1 is necessary for colony formation in hMSCs. We modulated *HOTAIR* expression in ESFT cell lines and hMSCs, alone and with concomitant *EWS-FLI1* expression. This change in *HOTAIR* expression was associated with significant gene expression changes across the transcriptome, including a set of genes modulated across all models, as determined by next-generation RNA-Sequencing (RNA-Seq). We further demonstrated by Chromatin Immunoprecipitation and Sequencing (ChIP-Seq) that *HOTAIR* expression induced H3K4 demethylation across the genome including in a significant number of genes with corresponding repression of expression.

## RESULTS

### *HOTAIR* is overexpressed, independently of *EWS-FLI1*, in ESFT cell lines and primary tumors as compared to normal tissues and mesenchymal stem cells

We first assessed the expression of *HOTAIR* in ESFT as compared to normal tissues, utilizing the OncoGenomics DB of the National Cancer Institute (https://pob.abcc.ncifcrf.gov/cgi-bin/JK). We examined next-generation RNA-sequencing (RNA-Seq) data annotated there for ESFT, including 50 cell lines and 72 primary tumor RNA samples(6), with expression normalized to that of a set of samples of normal adult tissues (Figure 1A). *HOTAIR* was expressed in all samples tested, with over a log-fold higher expression in >90% of cell lines and tumors as compared to normal tissues.

**Figure 1:**
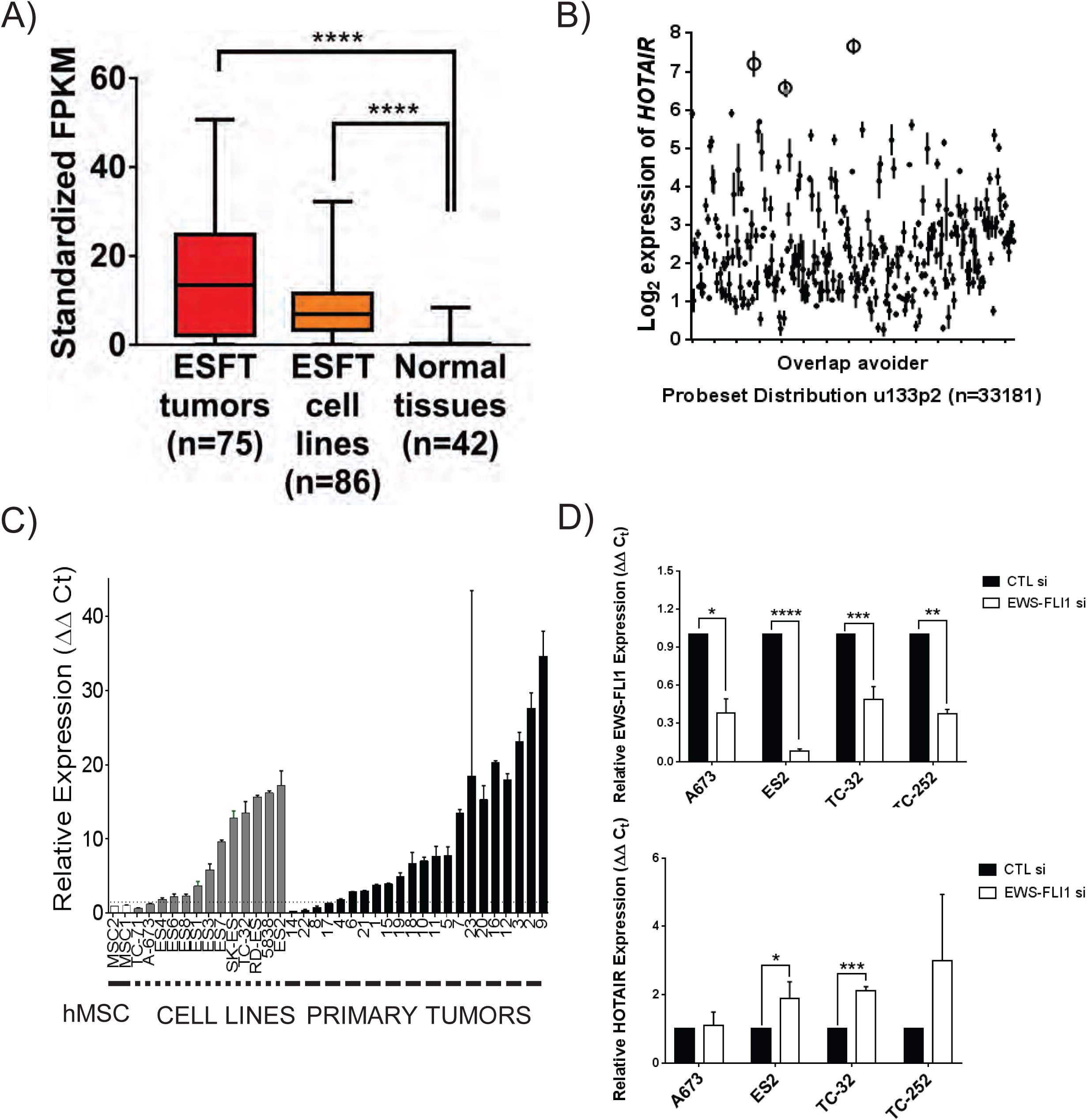
*The lncRNA* HOTAIR *is highly expressed in primary Ewing sarcoma tumors and cell lines*. **A**) *HOTAIR* expression levels in RNA-Seq data from Oncogenomics Database at the National Cancer Institute (NCI). 50 cell lines and 72 primary ESFT tumors were evaluated and normalized to a set of normal human tissues. Expression is expressed as standardized fragments per kilobase of transcript per million reads (FPKM); All ESFT samples have *HOTAIR* expression above average expression in normal tissues. **B)** *HOTAIR* expression levels as assessed by Affymetrix u133p2 microarray in 311 tumor datasets maintained at R2 Genomics Analysis and Visualization Platform. Open circles indicate ESFT datasets, which have the highest expression of all datasets analyzed. **C)** *HOTAIR* expression as measured by RT-qPCR in 13 ESFT cell lines (gray bars) and 22 primary ESFT tumors (black bars), as compared to primary mesenchymal stem cells (MSC, white bars) from two healthy donors, used as a normal control. 11/13 cell lines and 17/22 primary samples had over two-fold expression of *HOTAIR* as compared to MSC. **D)** *EWS-FLI1* (top) and *HOTAIR* (bottom) mRNA expression in ESFT cell lines treated with nonsilencing control siRNA or siRNA targeting *EWS-FLI1*. In all cell lines, knockdown of *EWS-FLI1* does not result in repression of *HOTAIR* expression and leads to a modest but significant increase in expression in ES2 and TC32 cell lines.

We next examined expression in ESFT as compared to other cancer types. Using the R2 Genomics Analysis and Visualization platform, (http://R2.amc.nl), we compared *HOTAIR* expression among cancer gene expression datasets that were analyzed using Affymetrix u133 microarrays (Figure 1B). The three datasets with the highest average *HOTAIR* expression were comprised of ESFT samples (16–18), as compared to 311 other datasets including other cancer cell lines, non-ESFT primary tumor sets, and sets of mixed normal and cancerous samples.

We validated these findings directly by analysis of 13 ESFT cell lines, 22 primary tumor RNA samples, and 2 primary hMSC samples by RT-qPCR. Using a threshold of two-fold expression as compared to hMSC, 11/13 cell lines and 17/22 tumor samples had high *HOTAIR* expression (Figure 1C). This consistent overexpression of *HOTAIR* in ESFT supports a functional role for the lncRNA in this cancer.

We hypothesized that *HOTAIR* expression in ESFT is an *EWS-FLI1*-independent event. We confirmed this by knocking down *EWS-FLI1* expression in 4 ESFT cell lines by siRNA (Figure 1D). In all four cell lines, knockdown of *EWS-FLI1* did not result in loss of expression of *HOTAIR*, and in two lines this knockdown actually led to a modest upregulation of *HOTAIR* expression. Thus, *HOTAIR* expression in ESFT cell lines is not driven by EWS-FLI1 and represents an independent biological pathway.

### *HOTAIR* expression correlates with anchorage independent colony formation in ESFT cell lines and hMSCs while maintaining MSC characteristics

We examined the phenotypic effects of *HOTAIR* in ESFT cell lines by first knocking down expression using shmiRNA in the ES2, TC32, and SK-ES cell lines. We repressed expression to 20–50% of baseline *HOTAIR* expression (Figure 2A), with some variation among the cell lines. We were unable to maintain viable cells with expression below this level for each cell line, supporting a role for *HOTAIR* in maintaining cell viability. We also confirmed that loss of *HOTAIR* expression had no significant effect on EWS-FLI1 protein expression (SI 1A).

**Figure 2:**
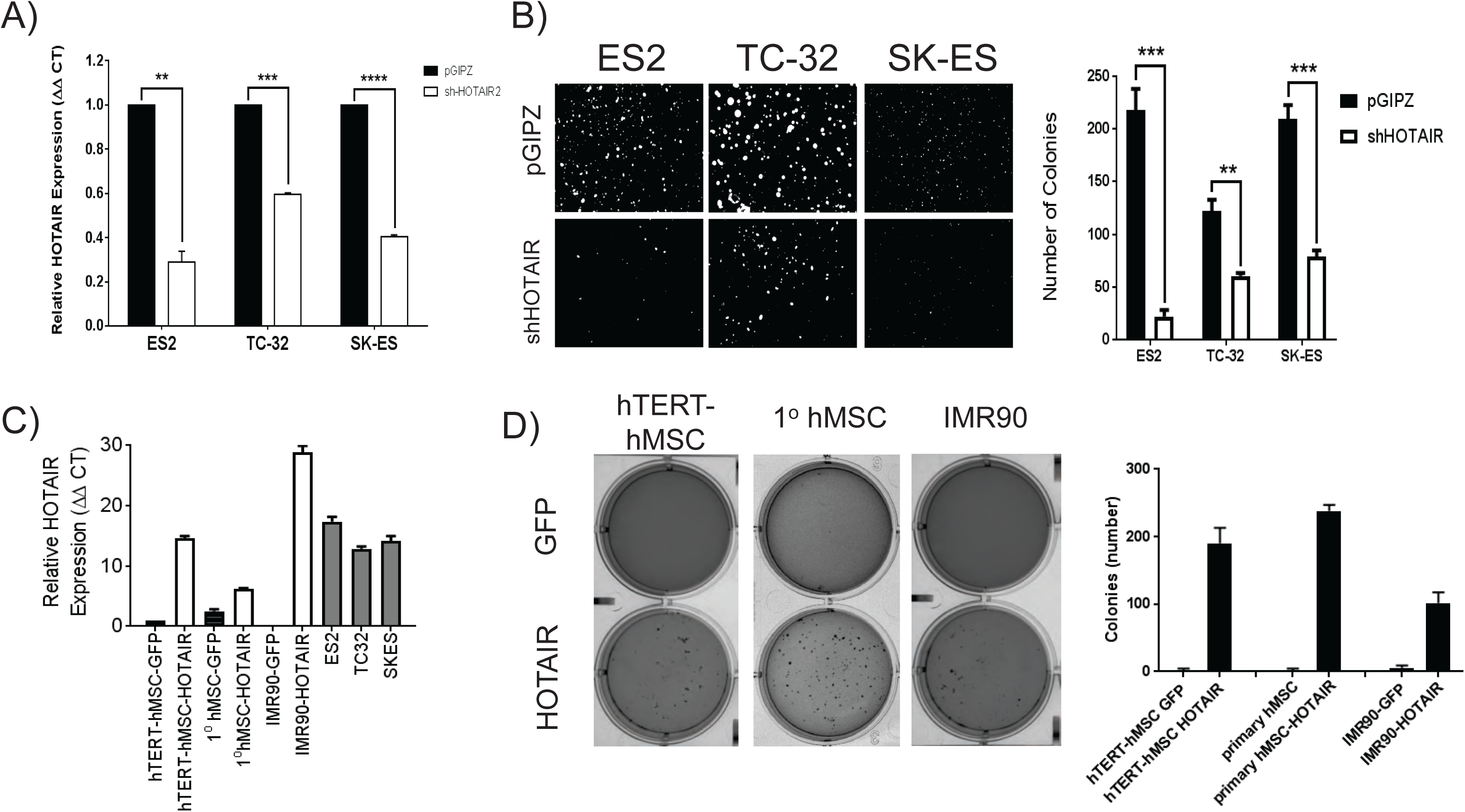
*HOTAIR is required for anchorage-independent growth of ESFT cells*. **A)** *HOTAIR* expression in three ESFT cell lines, infected with lentivirus-delivered GFP-coexpressing shmiRNA against *HOTAIR* or nonsilencing control (pGIPZ). *HOTAIR* was repressed to 25–50% of baseline expression. **B)** Soft agar colony formation assay of ESFT cell lines from figure 2A. Representative fluorescence photomicrographs of the colonies are shown. Colony counts from three independent experiments are shown in the bar graph to the right (error bars standard deviation (SD)). For all cell lines, fewer colonies were formed in cells with repression of *HOTAIR*. **C)** *HOTAIR* expression as measured by RT-qPCR in stable selected hTERT-hMSCs, 1^o^ hMSCs, and IMR90 lung fibroblasts transfected with vector expressing GFP alone or GFP and *HOTAIR*. Expression is compared to basal *HOTAIR* expression in three ESFT cell lines. Expression is normalized to expression in the hTERT-hMSC-GFP control. **D)** Soft agar colony formation assay of hTERT-hMSC, 1^o^ hMSC, and IMR90 cells (from Figure 2C) after two weeks of growth. Representative photomicrographs are shown in the left panels. Colony counts from three independent experiments are shown in the bar graph to the right (error bars SD). More colonies were formed from hTERT-hMSCs, 1^o^ hMSCs, and IMR90 cells expressing *HOTAIR* than GFP alone. p<0.001 for all three lineages.

We next examined the effects of *HOTAIR* repression on the growth of ESFT cells. Proliferation of these cells in two-dimensional tissue culture did not show any significant difference at this level of repression (SI 1B). However, prior work in Ewing sarcoma biology demonstrated that inhibition of LSD1, shown in other cancers to interact with *HOTAIR*, had no significant effect on proliferation but did alter 3D tumorsphere growth(19). Accordingly, we evaluated growth of tumorspheres in soft agar. We found in all three cell line models that repression of *HOTAIR* resulted in a significant decrease in anchorage-independent colony formation in soft agar (Figure 2B).

hMSCs are the presumptive cell of origin for Ewing sarcoma but they poorly form colonies in 3D culture in vitro and lack tumorigenicity in vivo(20, 21). We evaluated how exogenous *HOTAIR* expression affected the growth of these cells. We initially used hTERT-immortalized hMSCs for this assay for ease of use in *in vitro* culture. These cells phenocopy primary hMSCs except for avoiding senescence, and they poorly form colonies in 3D culture. We overexpressed *HOTAIR* or a control vector in these cells, at levels comparable to ESFT cell lines (Figure 2C). *HOTAIR* induced colony formation in soft agar, whereas expression of the empty vector did not (Figure 2D). This ability to enable anchorage-independent growth supports a role for *HOTAIR* not only in cell proliferation and viability but also in tumorigenesis. We repeated this experiment with early passage primary mesenchymal stem cells and found that overexpression of *HOTAIR* induced colony formation in soft agar (Figure 2C and D). We also used the wildtype IMR90 lung fibroblast cell line, which has been shown to be useful as a model of EWS-FLI1 expression(22) and does not form colonies in 3D culture. Again, overexpression of *HOTAIR* induced anchorage-independent colony formation in these cells (Figure 2C and D).

MSCs are defined by characteristics that include the ability to grow on plastic, expression of the surface markers CD73, CD90, and CD105, and maintenance of differentiation capacity along mesenchymal lineages(23). ESFT also have most of these properties, distinct from other sarcomas(24, 25). We confirmed that HOTAIR-expressing hTERT-hMSCs maintained these properties, as did the ESFT cell lines with knockdown of *HOTAIR* expression. Analysis by flow cytometry showed that control cells and cells with modulated *HOTAIR* expression had strong surface expression of CD73, CD90, and CD105, and had no significant alteration of other lineage markers (SI 2). The cells with modulated *HOTAIR* expression were also able to be differentiated into osteoblasts or adipocytes in patterns similar to their GFP-expressing controls (SI 3).

### *HOTAIR* interacts with LSD1 in ESFT and requires its 3’ interaction domain for tumorsphere formation capacity in hMSCs

We performed formaldehyde-crosslinked RNA immunoprecipitation and confirmed that, in ESFT cell lines, *HOTAIR* interacts with LSD1 (SI 4A). We wanted to evaluate if this interaction is necessary for the anchorage-independent colony formation seen in the hMSC models. The interaction domain of *HOTAIR* with LSD1 has been previously defined(9). We generated a *HOTAIR* deletion construct that lacked the 3' LSD1 interaction domain (Figure 3A). We then expressed this construct or wild-type *HOTAIR* in hTERT-hMSCs. We confirmed that expression of each of the constructs was not significantly different amongst the pools of cells (SI 4B), then repeated the anchorage-dependent colony formation assay. As compared to the cells with overexpression of wild-type *HOTAIR*, those cells with expression of the mutant *HOTAIR* had a markedly diminished or absent colony formation capacity in soft agar (Figure 3B). This loss of function suggests that *HOTAIR* must interact with LSD1 to support tumor formation in hMSCs and, analogously, in ESFT.

**Figure 3:**
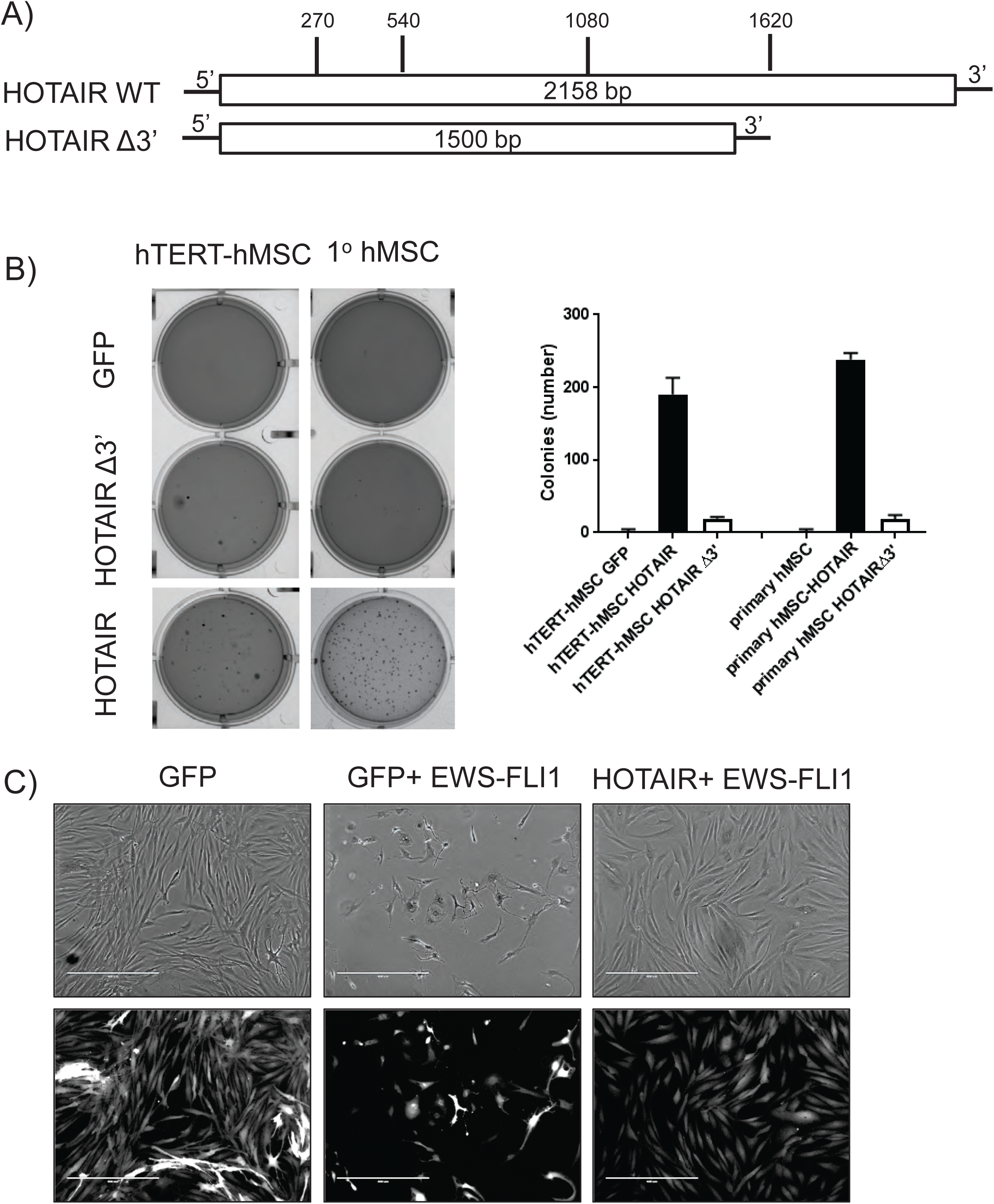
Anchorage independent growth mediated by HOTAIR requires both 5' and 3' ends. **A)** Schematic of wild-type and deletion mutant of *HOTAIR* generated with, the LSD1-interacting 3’ domain deleted. Variant was cloned into GFP-coexpressing plasmid vector for transfection into hTERT-hMSCs, which were then selected for stable integrants. **B)** Soft agar colony formation assay of hTERT-hMSC (left) and 1^o^ hMSC cell lines with GFP control, 3'-deleted *HOTAIR* mutant or wild-type *HOTAIR* expression, after three weeks of growth. Representative photomicrographs are shown of each variant, with colony counts of three independent experiments shown in the bar graph (error bars SD). Only cells expressing wild-type *HOTAIR* form significantly more colonies than GFP-expressing control. **C)** Phase contrast and fluorescent photomicrographs of hTERT-hMSC cells transfected with GFP alone (left), GFP and *EWS-FLI1* (middle) or GFP and *HOTAIR* first followed by *EWS-FLI1* (right). Cells expressing only *EWS-FLI1* undergo apoptosis rapidly, but cells expressing *HOTAIR* stably first then *EWS-FLI1* are fully viable.

### *HOTAIR* primes hMSCs for tolerance of EWS-FLI1 and maintains cell viability

As previously noted, exogenous expression of *EWS-FLI1* in most normal and malignant cell lines rapidly causes cell death. We confirmed this phenotype by expression of either GFP alone or *EWS-FLI1* and GFP in hTERT-hMSCs. Within 48 hours, the *EWS-FLI1*-overexpressing hTERT-hMSCs morphologically changed with cell shrinkage and nuclear collapse, and no viable cells could be seen at 72 hours (Figure 3C), whereas the GFP-expressing cells remained unchanged. We created a vector for simultaneous co-expression of *HOTAIR* and *EWS-FLI1* and transfected hTERT-hMSCs with this vector. The hTERT-hMSCs again underwent rapid apoptosis. We next used the hTERT-hMSCs that stably expressed *HOTAIR* and overexpressed *EWS-FLI1* in those cells. In marked contrast, these cells stably expressed both *EWS-FLI1* and *HOTAIR*, as confirmed by western blot and RT-qPCR respectively (SI 5A and B), and remained viable without morphologic evidence of differentiation (Figure 3C). These cells also formed tumorspheres in soft agar, at an increased rate as compared to hTERT-hMSC-*HOTAIR* cells. We were similarly able to stably express *EWS-FLI1* in primary hMSCs and IMR90 cells with stable *HOTAIR* expression (SI 5A-C). These data support the hypothesis that *HOTAIR* expression in the ESFT precursor cell primes the cell to tolerate the subsequent chromosomal translocation that results in *EWS-FLI1*, driving malignant transformation to ESFT.

### *HOTAIR* is necessary but not sufficient for tumor xenograft growth in immunodeficient mice

We next evaluated how *HOTAIR* affects the tumor-initiating capacity of ESFT cells *in vivo*. We implanted the ESFT cell lines above, with shRNA-mediated knockdown of *HOTAIR* or nonsilencing control, into the flanks of SCID mice. The tumor xenografts of cells with repressed *HOTAIR* expression had significantly slower tumor growth across all three cell lines as compared to control (p<0.001 for all three cell lines, Figure 4). We also implanted into the SCID mice the primary hMSC, hTERT-hMSC, and IMR90-GFP, *-HOTAIR*, and *-HOTAIR-EWS-FLI1* cells, at cell numbers up to 1×10^7^cells/implant, but no tumors grew in any of the mice after 12 weeks. These data suggest that *HOTAIR* is necessary but not sufficient for tumor growth in cells in the context of concomitant EWS-FLI1 expression.

**Figure 4:**
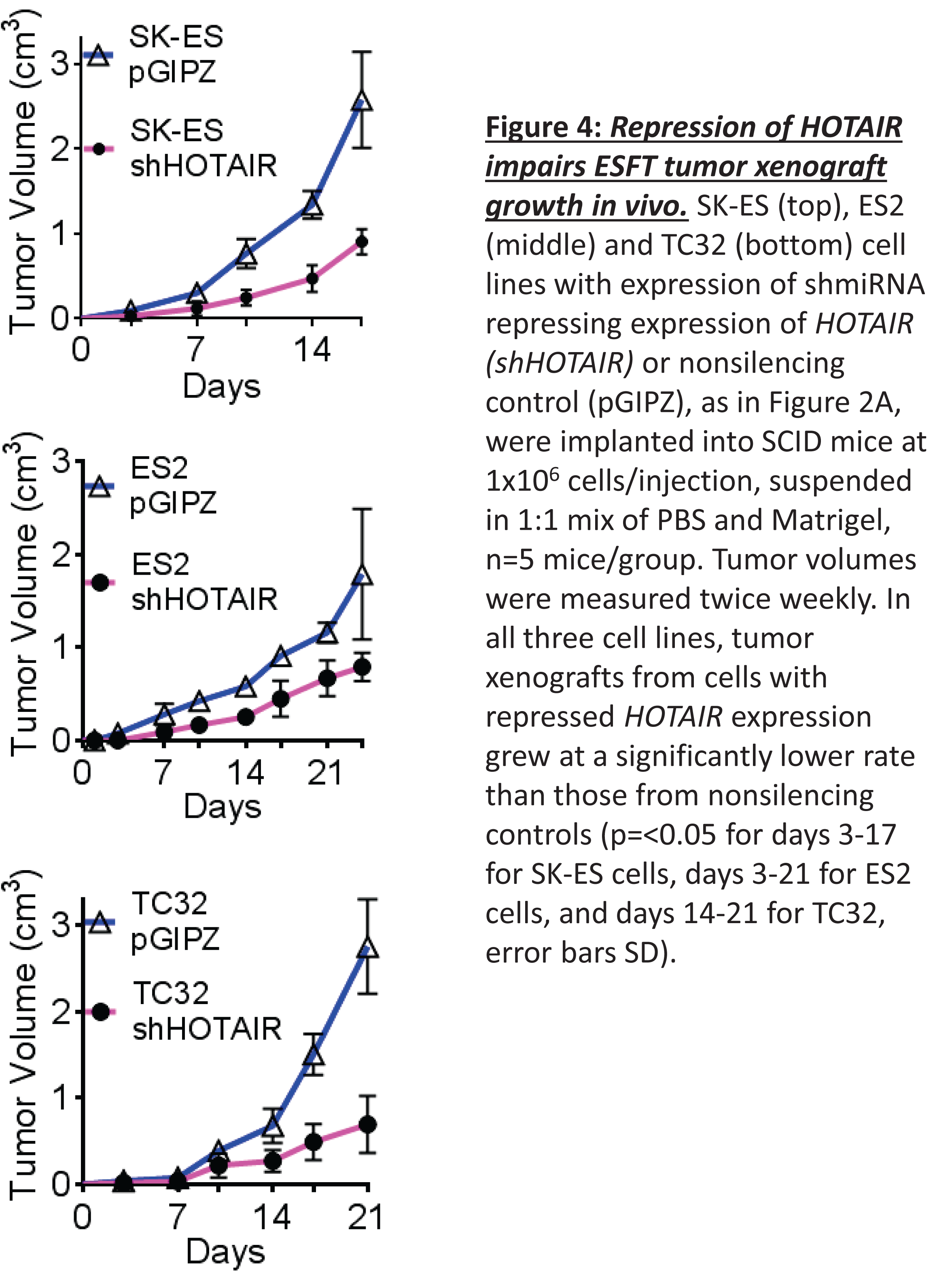
Repression of HOTAIR impairs ESFT tumor xenograft growth in vivo. SK-ES (top), ES2 (middle) and TC32 (bottom) cell lines with expression of shmiRNA repressing expression of *HOTAIR (shHOTAIR)* or nonsilencing control (pGIPZ), as in Figure 2A, were implanted into SCID mice at 1×10^6^ cells/injection, suspended in 1:1 mix of PBS and Matrigel, n=5 mice/group. Tumor volumes were measured twice weekly. In all three cell lines, tumor xenografts from cells with repressed *HOTAIR* expression grew at a significantly lower rate than those from nonsilencing controls (p=<0.05 for days 3–17 for SK-ES cells, days 3–21 for ES2 cells, and days 14–21 for TC32, error bars SD).

### *HOTAIR* expression in hMSCs and ESFT cells is associated with gene expression changes affecting cell adhesion, extracellular matrix, and embryonic stemness

Given the ubiquitous expression of *HOTAIR* in ESFT cell lines and tumors and its tumorigenic effect in hMSCs, we aimed to characterize the effects of *HOTAIR* on gene expression in these models. We used the hTERT-hMSC models as described above, with expression of *HOTAIR* alone or with subsequent expression of *EWS-FLI1*; these models expressed *HOTAIR* 5–8 fold over baseline (SI 5A), in a range observed in ESFT tumors. In ESFT cell lines, we knocked down *HOTAIR* expression by use of GapmeR antisense oligonucleotides. Treatment of cells with GapmeRs for 48 hours repressed expression to <30% of baseline in 3 ESFT cell lines, ES2, A673, and SK-ES (SI 6A).

We examined mRNA expression in these models by RNA-Seq, identifying transcripts with significantly different expression as compared to the control (adjusted p-value<0.05). In the hTERT-hMSC cells, we identified 2781 transcripts upregulated and 2169 transcripts downregulated in the HOTAIR-overexpressing cells, and 3888 transcripts upregulated and 2435 transcripts downregulated in the *HOTAIR-EWS-FLI1* overexpressing cells, as compared to the GFP control (SI 6B). We identified the top 100 differentially-expressed transcripts between the *HOTAIR-EWS-FLI1* expressing cells and GFP control and examined their expression in all three cell types (Figure 5A, SI data). Among these transcripts, we found a set of genes that are downregulated in cells with increased *HOTAIR* expression independently of EWS-FLI1. We also defined a set of transcripts whose expression is upregulated in cells with increased *HOTAIR* expression independently of EWS-FLI1, as well as a larger set of transcripts specifically upregulated by EWS-FLI1. These patterns are similarly seen in the complete dataset of all differentially expressed genes (SI Data).

In the ESFT cells, a different pattern emerged. Treatment of ES2, A673, and SK-ES cells with the GapmeR repressing *HOTAIR* caused more genes to be upregulated than downregulated as compared to control (972 upregulated vs. 472 downregulated for ES2, 717 upregulated vs. 506 downregulated for A673, 1080 upregulated vs 815 downregulated for SK-ES, SI 3C). Across all three cell lines there was an intersection set of 81 differentially expressed genes; of these, 49 were upregulated when cells were treated with GapmeRs as compared to control (Figure 5B, SI data). This suggests that *HOTAIR* expression plays a greater role in directing gene repression in ESFT cells than in driving gene expression.

**Figure 5:**
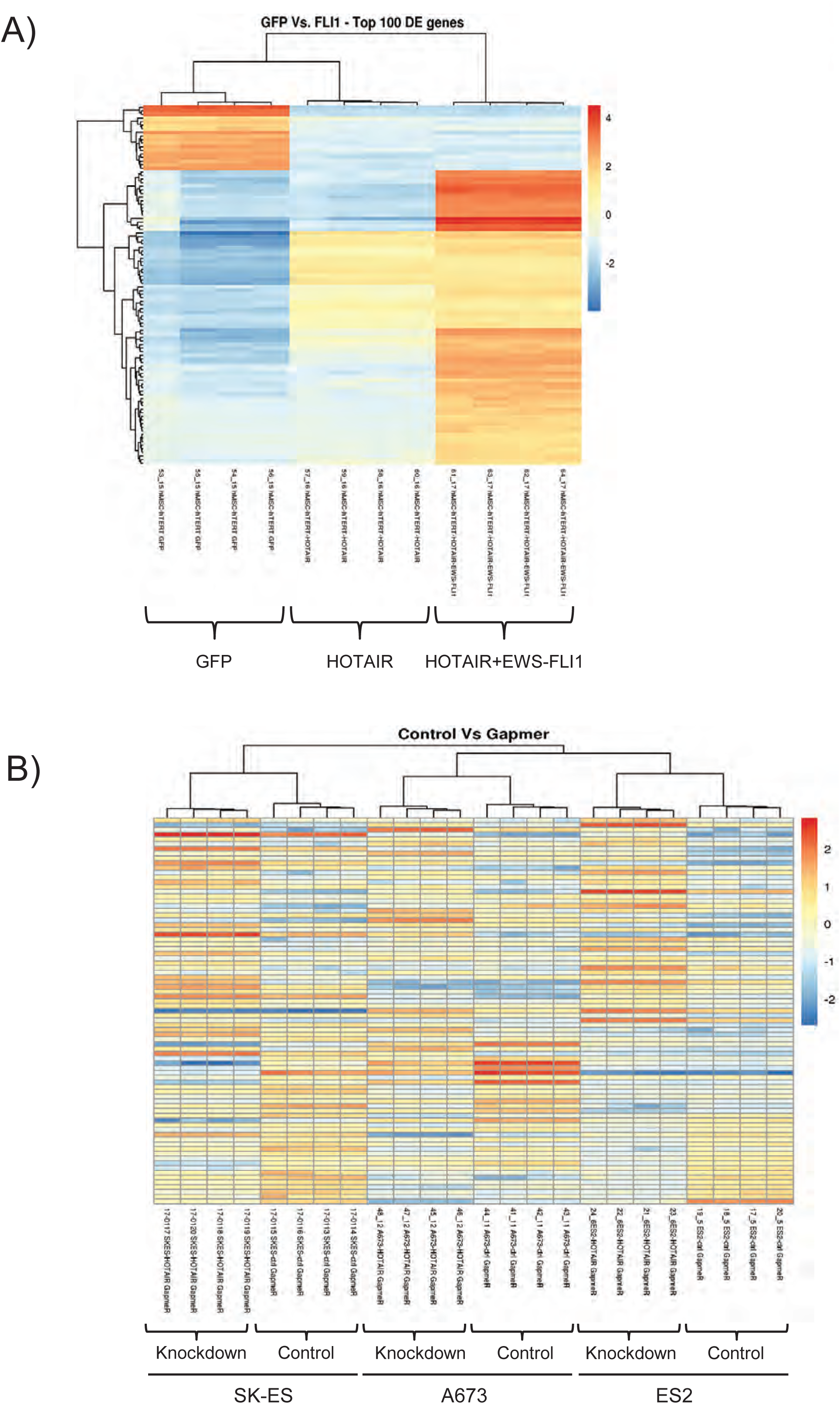
HOTAIR regulates gene expression independent of EWS-FLI1. **A)** Heatmap of top 100 differentially expressed genes in hTERT-hMSC cells expressing GFP, *HOTAIR*, or *HOTAIR* and *EWS-FLI1*. There is a set of genes repressed in correlation with *HOTAIR* specifically (top section) and another set specifically upregulated in correlation with *HOTAIR* (middle section), with additional genes whose expression correlate best with *EWS-FLI* co-expression. **B)** Heatmaps of 81 differentially regulated genes in ES2, SK-ES, and A673 cells treated with nonsilencing GapmeR or GapmeR targeting *HOTAIR*. The majority of genes identified demonstrate increased expression in cells treated with GapmeR repressing *HOTAIR* expression (i.e., gene expression is inversely correlated with *HOTAIR* expression).

We evaluated which biological pathways were affected by *HOTAIR* expression in these cell line model sets. We noted significant heterogeneity of the specific genes affected within each cell line, both in terms of basal expression and change in expression relative to *HOTAIR*. As such, we identified those differentially-expressed genes in the hTERT-hMSC cells and any of the ESFT cell lines and defined common biological pathways affected by those genes (Table 1, SI Data). *HOTAIR* expression correlated with repression of genes involved in cell differentiation, extracellular matrix organization, and cell-cell adhesion, and with upregulation of genes affecting angiogenesis, cell motility, and biological adhesion. These functions correlate with tumorigenesis, metastasis, and inhibition of differentiation.

**Table 1:** Enrichment gene ontologies for differentially expressed genes - biologically relevant processes.

| Biological Process Set | Percentage of Genes Represented in Combined Gene Set | Correlation with HOTAIR expression | Enrichment adjusted p-value |
| --- | --- | --- | --- |
| Angiogenesis (GO:0001525) | 2.75 | Upregulated | 3.51E-2 |
| Chemotaxis (GO:0006935) | 4.45 | Upregulated | 6.4E-5 |
| Cell migration (GO:0016477) | 6.45 | Upregulated | 5.86E-5 |
| Locomotion (GO:0040011) | 9.26 | Upregulated | 1.39E-9 |
| Biological adhesion (GO:0022610) | 9.61 | Upregulated | 2.46E-5 |
| Connective tissue development (GO:0061448) | 2.09 | Downregulated | 2.79E-2 |
| Mesenchyme development (GO:0060485) | 2.14 | Downregulated | 8.38E-3 |
| Homophilic cell adhesion via plasma membrane adhesion molecules (GO:0007156) | 2.09 | Downregulated | 1.15E-5 |
| Extracellular matrix organization (GO:0030198) | 3.32 | Downregulated | 7.41E-6 |
| Skeletal system development (GO:0001501) | 4.65 | Downregulated | 2.44E-7 |
| Cell-cell adhesion (GO:0098609) | 6.36 | Downregulated | 2.99E-2 |
| Cytoskeleton organization (GO:0007010) | 6.98 | Downregulated | 2.26E-3 |
| Nervous system development (GO:0007399) | 15.67 | Downregulated | 2.15E-9 |
| Cell differentiation (GO:0030154) | 22.42 | Downregulated | 3.36E-12 |

### *HOTAIR* expression induces H3K4 demethylation at the promoters of *HOTAIR-repressed* genes

As previously mentioned, RNA immunoprecipitation experiments in ESFT cell lines confirmed that *HOTAIR* associates with LSD1 (SI 4A). LSD1 has been shown in other cancers to repress gene expression through this association with *HOTAIR* by demethylation of histone 3 at lysine 4 (H3K4), particularly from a dimethylated (Me2) to a monomethylated (Me) state(9). We hypothesized that differential *HOTAIR* expression in our model systems would alter LSD1-directed histone methylation at gene promoters, repressing gene expression. As such, we performed Chromatin Immunoprecipitation-Sequencing (ChIP-Seq) for H3K4Me1 and H3K4Me2, using ES2 and SK-ES cells with shRNA-mediated knockdown of *HOTAIR* or nonsilencing control as described above, and the hTERT-hMSC-GFP and hTERT-hMSC-*HOTAIR* cells. In the ES2 cell lines, we examined those genes that had increased expression when *HOTAIR* expression was repressed (Figure 5A and B). We found that the loss of *HOTAIR* also resulted in decreased H3K4Me1 and increased H3K4Me2 around the transcriptional start sites (TSS) of these genes (Figure 6, SI data). In the hTERT-hMSC cells, *HOTAIR* expression similarly induces increased H3K4Me1 around the TSS of HOTAIR-repressed genes, with a less-pronounced but still significant decrease in H3K4Me2 (SI 7, SI data). In the SK-ES cells, loss of *HOTAIR* is again associated with significantly decreased H3K4Me broadly adjacent to the TSS, though not as markedly as in the ES2 or hTERT-hMSC cells, and with a particularly increased amount of H3K4Me2 immediately adjacent to the TSS (SI 8, SI data). These data demonstrate that genes repressed in the context of *HOTAIR* expression consistently have histone methylation changes at their promoters associated with LSD1-mediated epigenetic repression.

**Figure 6:**
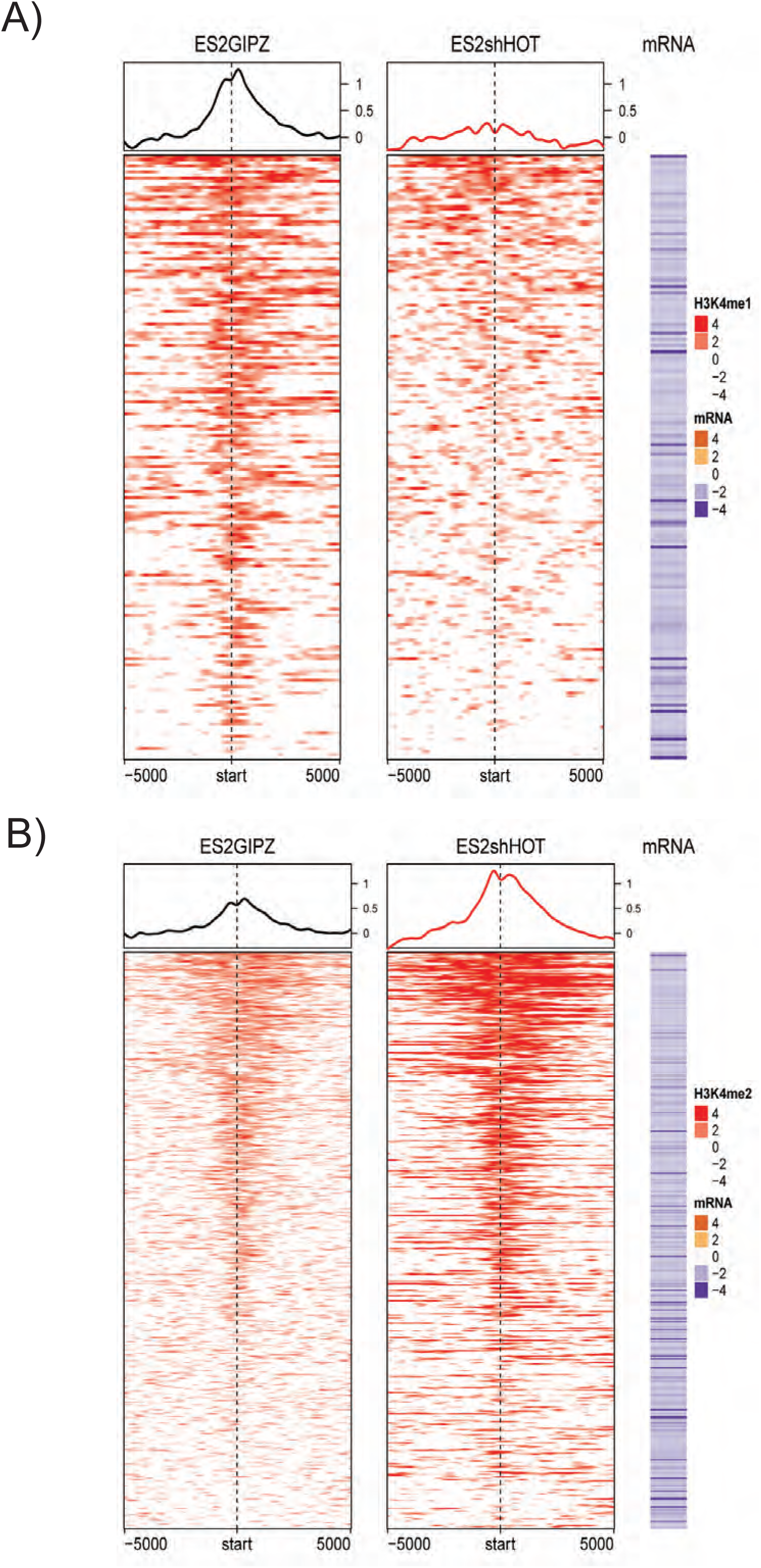
Genes repressed by HOTAIR expression have increased H3K4 monomethylation and decreased H3K4 dimethylation around their transcriptional start sites (TSS) in ES2 cells. Chromatin Immunoprecipitation and Sequencing (ChIP-Seq) identified genomic regions with differential H3K4Me1 and H3K4Me2 binding in ES2-pGIPZ cells as compared to ES2-shHOTAIR, with FDR<0.05. A) Profiles of HOTAIR-induced H3K4Me1 modification around TSS of HOTAIR-repressed genes. Each row corresponds to HOTAIR-repressed genes (195 genes). Color for H3K4Me1 profile represents log_2_ ratio of ChIP to input control. Positive (red) represents enrichment of H3K4Me1. Mean H3K4Me1 profiles in each cell line are plotted on top. The corresponding mRNA expression (log2 ratio) is shown on the right panel. Negative (purple) represents decreased expression in ES2-pGIPZ compared to ES2-shHOTAIR (i.e. genes that are repressed by HOTAIR). H3K4Me1 was differentially bound adjacent to the TSS of these *HOTAIR-repressed* genes in the context of *HOTAIR* expression in ES2-pGIPZ cells as compared to ES2-shHOTAIR2 cells. B) Profiles of HOTAIR-repressed H3K4Me2 bindings in ES2-pGIPZ and ES2-shHOTAIR around TSS of HOTAIR-repressed genes. Each row corresponds to HOTAIR-repressed genes (495 genes). Color for H3K4Me2 profile represents log_2_ ratio of ChIP to input control. Positive (red) represents enrichment of H3K4Me2. Mean H3K4Me2 profiles in each cell line are plotted on top. The corresponding mRNA expression (log2 ratio) is shown on the right panel. Negative (purple) represents decreased expression in ES2-pGIPZ compared to ES2-shHOTAIR (i.e. genes that are repressed by HOTAIR). H3K4Me2 was differentially bound adjacent to the TSS of HOTAIR-repressed genes in the context of loss of *HOTAIR* in ES2-shHOTAIR cells as compared to the ES2-pGIPZ control cells (i.e., loss of *HOTAIR* correlates with increased H3K4Me2 binding and increased expression of these correlated genes).

## DISCUSSION

The Ewing sarcoma family of tumors is characterized by the *EWS-FLI1* fusion gene or similar fusion genes. The resultant fusion proteins are the drivers of oncogenesis in these tumors, but these proteins are toxic to most studied normal cells and even other types of cancer cells. The paucity of additional mutations in these tumors suggests that specific epigenetic profiles may allow viability and tumorigenesis of the *EWS-FLI1*-expressing cells. Our studies suggest that the expression of the lncRNA *HOTAIR* redirects epigenetic regulation to induce just such a permissive state and allow formation of Ewing sarcoma tumors. *HOTAIR* is consistently overexpressed in virtually all ESFT primary tumors and cell lines examined by either microarray(18), RNA-Seq(26), or RT-qPCR. This is in stark contrast to the minority of ESFT tumors with genomic mutations, as assayed in three large whole genome sequencing studies(6-8). This consistency of expression supports a function for *HOTAIR* within this cancer. We additionally confirmed that *HOTAIR* expression is not driven by EWS-FLI1, further supporting an independent function in ESFT tumorigenesis and viability.

Phenotypically, *HOTAIR* directly affects growth of ESFT. In the hTERT-hMSC model, overexpression impacted both two-dimensional cell growth and anchorage-independent tumorsphere formation. In contrast, in the ESFT cell lines, knockdown of *HOTAIR* expression repressed only tumorsphere formation. This difference may be explained by the necessity of *HOTAIR* in ESFT. Using shmiRNA methods and strict selection conditions, knockdown below 25% was impossible. We additionally tried to knockout the *HOTAIR* alleles in ESFT cell lines by adapting an RNA-destabilizing element (RDE) (27) that also integrated green or red fluorescent proteins into the locus. While we could successfully insert one RDE into cells, we could not subsequently generate viable cells that lost expression of the second allele (data not shown). Our inability to completely silence *HOTAIR* expression suggests that a single copy of *HOTAIR* is both necessary and sufficient for viability of the ESFT cell lines we used. As such, repression of *HOTAIR* expression in our ESFT cell lines to a level that inhibits proliferation may simply make them nonviable. A conditional expression system may allow for titratable *HOTAIR* expression while genomic copies of the gene are knocked out, followed by phenotypic evaluation as *HOTAIR* expression is reduced. Nonetheless, the effects of *HOTAIR* loss in ESFT 3D colony and tumor formation phenocopy those effects found with inhibition of LSD1 in ESFT(19), supporting a pathophysiologic role of *HOTAIR* in ESFT tumor viability.

The gene expression analysis of our cell line models further defines how *HOTAIR* affects ESFT formation and viability. For the RNA-Seq analyses, we aimed to evaluate the maximal effect of *HOTAIR* expression. We generated hTERT-hMSC cell lines with stable *HOTAIR* and *EWS-FLI1* expression consistent with that seen in ESFT cell lines. However, to maximally reduce *HOTAIR* expression in ESFT cell lines, we used GapmeRs for transient knockdown. This approach may limit the ability to measure *HOTAIR's* activity in modulating epigenetic regulation, which has some rapidly altered features but also additional characteristics only seen over time. These different methods used to modulate *HOTAIR* expression likely contribute to the difference in the number of genes identified in the different models. In the stable hTERT-hMSC system, thousands of genes were found to be differentially expressed, while comparatively fewer were identified in the transient ESFT models. Nonetheless, key effects were still identified. In particular, there is a set of genes whose expression is regulated by *HOTAIR* independently of EWS-FLI1 and conversely another set regulated specifically by EWS-FLI1, suggesting complementary functions of the two genes in ESFT tumorigenesis.

In both ESFT and hTERT-hMSC models of *HOTAIR* and *EWS-FLI1* expression, differential expression of genes integral to tumor formation were identified, including cytoskeletal and adhesion proteins (collagens, keratins, cadherins and protocadherins) and matrix metalloproteases. The specific genes affected in each individual cell line are variable, consistent with data showing significant diversity in the gene expression profiles of ESFT in general (26, 28). Regardless, we identified key pathways affecting tumorigenesis, including cell motility and migration, across the different models. It is particularly noteworthy that these pathways were previously identified to be biomarkers of survival in patients(28).

We also identified other critical pathways that are differentially affected by *HOTAIR*, including DNA repair pathways and normal differentiation and development. ESFT are characteristically undifferentiated, which contributes to the inability to as yet define the cell of origin for these tumors. Additionally, errors in DNA repair pathways have been hypothesized to contribute to oncogenesis(29, 30). *HOTAIR* may function in the cell of origin to maintain the cells in an undifferentiated and EWS-FLI1 receptive state, tolerant of DNA damage induced by pathways activated by the fusion protein.

*HOTAIR* was originally demonstrated in fibroblasts to function as a scaffold for the LSD1/REST complex at its 3’ end and PRC2 at its 5’ end(9, 31). We confirmed the interaction of *HOTAIR* and LSD1, the correlated effect on histone methylation and gene expression in ESFT cell lines, and the necessity of this interaction for the anchorage-independent growth phenotype seen in the hTERT-hMSC model. It is important to acknowledge that the effects of *HOTAIR* varied among the disease models, with far greater effects seen on gene expression in the hMSC models than in ESFT cell lines. This may have been due to greater effects of stable *HOTAIR* expression in contrast to the transient effects of *HOTAIR* repression by the GapmeRs, as noted above. Alternatively, *HOTAIR* may induce gene expression changes during ESFT transformation but may not be required to maintain gene expression at all sites after transformation occurs. A comprehensive evaluation of *HOTAIR* binding across the genome, and a comparison of that binding to histone methylation, will elucidate the direct effects of *HOTAIR* in ESFT.

Additional work is warranted in investigating other functions of *HOTAIR* on gene expression, including its interaction with PRC2 and other regulatory mechanisms, such as effects on DNA methylation independent of LSD1 or PRC2, as recently described(32). We have begun additional work on the interaction of *HOTAIR* and PRC2, but these studies are much more complicated. A prior study showed a lack of effects on H3K27 methylation despite the presence of *HOTAIR(9)*, which may be due to additional cofactors that allow PRC2 binding but prevent methyltransferase activity(33). We felt it important to make our present findings available while we perform more expansive studies on *HOTAIR* in ESFT, particularly because of the demonstration of LSD1's importance in ESFT(34) and the development of novel LSD1 inhibitors(19) for use in this disease. These prior studies showed functions of LSD1 through the Nucleosome Remodeling and Demethylation (NuRD) Complex, but our studies would support evaluation of the effects of the LSD1 inhibitors on HOTAIR-directed H3K4 modifications.

A recent publication by Amandio *et al*., has called the biological relevance of *HOTAIR* into question(35). They examined the effect of the mouse *Hotair* ortholog on mouse developmental biology, in follow-up to prior studies showing discordant effects in different mouse strains. They concluded that *Hotair* had little effect on mouse morphology or on H3K27 methylation patterns, questioning the necessity of the lncRNA and its role in disease. While intriguing, this work by Amandio *et al*. has its limitations. They did not examine the effects of *Hotair* on LSD1 function or targeting, a key pathway identified in our studies in ESFT. Additionally, they examined gene expression effects of *Hotair* deletion at a single embryonic timepoint in whole tissues consisting of dozens of cell types. *HOTAIR* expression and its effects have been shown to be widely variable across cell types and degree of differentiation(36). Our studies also examined *HOTAIR* in ESFT tumors and models of disease, which while heterogeneous in their composition, are largely comprised of a single cancer cell type. Loss of *Hotair* may not be associated with premature differentiation, but *HOTAIR* expression is associated with inhibition of differentiation, a feature observed by Amandio *et al*. Finally, prior study by that group demonstrated poor sequence homology between *Hotair and HOTAIR*, with the mouse lncRNA showing poor conservation of the domains responsible for EZH2 and LSD1 interaction(37). As they state, “it nevertheless suggests that the function of this RNA in mice is not identical to that described for its human cognate(37).” This fact is further supported by the scores of reports on the expression of *HOTAIR* in multiple cancers, its correlation with disease outcome, and its biological functions in these cancers. While studies are needed to discriminate the direct function of *HOTAIR* from its indirect effects on downstream targets, our work supports these effects and merits such additional study.

The identification of *HOTAIR* in ESFT has implications on the understanding of the disease and on its treatment. The driver of *HOTAIR* expression has yet to be confirmed, but its expression was previously described in normal embryonic stem cells(11) and cancer stem cells(38, 39). The ESFT cell of origin may inherently have high *HOTAIR* expression as a normal part of development or an error in that process. *HOTAIR* may prime these cells for EWS-FLI1-mediated malignant transformation. We are currently evaluating what additional genetic, transcriptional,or epigenetic events can collaborate with *HOTAIR* and EWS-FLI1 to induce tumor formation *in vivo* in our 1^o^ and hTERT-hMSC models. Further study of the regulation and function of *HOTAIR* in these models and in primary ESFTs may elucidate the oncogenesis of these cancers.

*HOTAIR* offers novel therapeutic options for ESFT. Indirectly, its specific activity through LSD1 suggests that these pathways may be specifically targetable in ESFT in a fashion that would be synergistic to attacks on EWS-FLI1. LSD1 inhibitors are being developed against ESFT(19), though that work was based on LSD1 interaction with the NuRD complex. Our findings suggest LSD1 may have diverse functions in ESFT, and disruption of *HOTAIR-*dependent LSD1 function may augment antitumor effects. New therapeutics may also be developed against *HOTAIR* itself, such as antisense oligonucleotides that have been previously generated against lncRNAs(40). As *HOTAIR* is not normally expressed in most tissues, this strategy may have fewer adverse effects systemically. Additional drugs could be identified or designed to interrupt *HOTAIR's* interactions with the epigenetic complexes, avoiding the normal *HOTAIR*-independent functions that LSD1 has in nonmalignant tissues. These avenues have promise in the generation of more specific and effective therapies for ESFT and other HOTAIR-expressing cancers.

## MATERIALS AND METHODS

### Cell Culture and Plasmids

ESFT cell lines ES1, ES2, ES3, ES4, ES6, ES7, ES8, A673 and TC-71 were obtained from Peter Houghton (UT Health Science Center, San Antonio, TX), and 5838, RD-ES, SK-ES and TC-32 were obtained from Timothy Cripe (Nationwide Children’s Hospital, Columbus, OH). IMR90 cells were obtained from Ryan Roberts (NCH). Cell line identities were confirmed upon receipt, start of studies, and annually by short tandem repeat (STR) profiling, using the Promega Powerplex 16 system, last performed on October 18, 2016 (completion of studies). Cells were tested for Mycoplasma with the SouthernBiotech Mycoplasma detection kit (Birmingham, AL), tested every 3 months, last in October 2016. Mycoplasma-contaminated cells were either replaced with uninfected cells from prior stores or treated with Plasmocin (Invivogen, San Diego, CA). The immortalized human marrow-derived mesenchymal stem cell line (hTERT-hMSC) was purchased from Applied Biological Materials, Inc. (British Columbia, Canada). Primary hMSCs were given as a generous gift from Edwin Horwitz (NCH). Cell lines were maintained in DMEM media supplemented with 10% FBS and Penicillin-Streptomycin at 370C with 5% CO2.

pLenti-HOTAIR-GFP was constructed by inserting the *HOTAIR* sequence at the PmeI restriction site in the pLenti-CMV-GFP-2A-Puro vector (Applied Biological Materials, Inc.) using the Infusion HD kit (TakaraBioUSA, Mountain View, CA). The *HOTAIR* deletion variants were then cloned from this plasmid by inverse PCR and Gibson cloning. For pLenti-EWS-FLI1-GFP, *EWS-FLI1* coding sequence containing a self-cleaving peptide sequence E2A at the 3’ end was inserted 5’ to the GFP-2A-PURO sequence using inverse PCR and Gibson cloning. All cloning primers are listed in the supplementary methods.

*HOTAIR* shmiRNA construct shHOTAIR2 was obtained from Dharmacon (Lafayette, CO, USA). The shmiRNA sequences from the inducible TRIPZ vector were cloned into the constitutively active pGIPZ vector by Gibson cloning. Antisense LNA GapmeR for *HOTAIR* (300631–724) and nonsilencing control (300615–08) was purchased from Exiqon (Vedbaek, Denmark).

### siRNA-mediated EWS-FLI1 knockdown

The Luciferase-RNAi (Luc-RNAi) and EWS/FLI-RNAi (EF-2-RNAi) constructs are previously described(4, 41). ESFT cells were infected with Luc-RNAi or EF-2-RNAi. Cells were grown in appropriate selection media for 72 hours and total RNA was extracted using an RNAeasy Kit (Qiagen). RT-qPCR was performed using SYBR green fluorescence for quantitation. Fold-change of genes was determined relative to the control gene, *RPL19*.

### Transfection, Virus Production and Transduction

The NEON Transfection System (ThermoFisher Scientific, Waltham, MA) was used for plasmid transfection. GapmeRs were transfected using Lipofectamine 3000 (ThermoFisher Scientific) per manufacturer protocol.

Lentivirus was generated using HEK-293T cells. Briefly, cells were seeded onto a 10 cm tissue culture plate at 70% confluency, then transfected with shmiRNA plasmid using the calcium phosphate precipitation method, pCMV-dR8.2, and pMD2.G at a ratio of 4:3:1. Lentivirus-containing media were collected at 48 and 72 hours, pooled and concentrated using Lenti-X Concentrator (Takara Bio USA). Target cells were then transduced with lentiviral particles using polybrene (8 μg/ml), with exposure for 24 hours then selection with puromycin for 3–7 days.

### RNA extraction and RT-qPCR

Total RNA from cells was extracted using NucleoSpin RNA purification kit (Takara Bio USA) per manufacturer instructions, then used for cDNA synthesis using Maxima RT cDNA First Strand Synthesis kit (ThermoFisher Scientific). qPCR was performed using KiCqStart SYBR Green qPCR ReadyMix (Sigma-Aldrich, St. Louis, MO) using the ABI PRISM 7900HT thermal cycler (ThermoFisher Scientific). Primers used are listed in the Supplementary Methods.

### Soft Agar Assay

Colony formation assay was performed on soft agar as previously described(42), with 5×10^3^ cells seeded in soft agar/media. After 2 or 3 weeks, fluorescent or brightfield photomicrographs were taken of the wells, and colonies were counted with the ImageQuant TL software (GE), with a colony defined as a cohesive collection of viable cells > 50 mcm in diameter. Experiments were completed in independent triplicates.

### Cell Proliferation Assay

Cell proliferation was measured using the alamarBlue cell viability assay (ThermoFisher Scientific). 2.5 × 10^3^ cells of each type were seeded in triplicate wells of a 96-well plate and grown in complete media with a separate plate for each time point. At each time point, alamarBlue reagent was added to each well and incubated for 2 hours. Fluorescence intensity was then measured, normalized for background fluorescence, then compared to day 0 readings. Experiments were completed in independent triplicates.

### Flow cytometry analysis

hTERT-hMSCs overexpressing wild-type *HOTAIR* or deletion mutants were analyzed on a BD LSRII flow cytometer (BD Biosciences, San Jose, CA) using allophycocyanin-conjugated anti-human CD105 and phycoerythrin-conjugated anti-human CD73 (BD Biosciences). Data were analyzed using FlowJo software v7.6 (Tree Star, Inc., Ashland, OR).

### Osteogenic and Adipogenic Differentiation

5 × 10^4 cells/well were plated in 24-well plates and were cultured in DMEM supplemented with 10% FBS. At about 90% confluency, the medium was switched to either NH OsteoDiff Medium or AdipoDiff Medium (Miltenyi Biotec Inc, Auburn, CA) and the cultures were maintained for 21 days. Osteoblastic differentiation was detected by calcium deposition visualized by staining with 1% alizarin red S solution (Sigma-Aldrich). Adipogenic differentiation was detected by Oil Red O staining (Sigma-Aldrich).

### Tumor xenograft growth assay

1×10^6^ ESFT cells (ES2, SK-ES, TC32) or 1×10^7^ 1^o^ hMSC, hTERT-hMSC, or IMR90 HOTAIR-EWS-FLI1 expressing cells were resuspended in 50 mcL PBS and mixed with 50 mcL of Matrigel (Corning) and injected subcutaneously into the flanks of SCID mice (Envigo), with 5 mice per injection group. Mice were monitored twice weekly for weight, body condition, and tumor size. Tumor volumes were estimated using the formula V=(length × (width)^2^)/2, per institutional protocol. Mice with SK-ES tumor xenografts were monitored for 17 days; mice with ES2 and TC32 tumor xenografts were monitored for 21 days; mice with hTERT-hMSC, 1^o^ hMSC, or IMR90 cell tumor xenografts were monitored for 12 weeks. Mice were then euthanized at the time endpoint and tumors harvested and fixed in formalin. All studies were designed in accordance with Nationwide Children’s Hospital IACUC guidelines and performed under IACUC-approved protocols.

### RNA immunoprecipitation (RIP)

RIP was performed as described(43, 44). 2×10^7^ cells were grown to 80% confluency, cross-linked with 1% formaldehyde, quenched with 2.66 M glycine, and then washed and lysed in IP lysis buffer. Samples were sonicated by probe sonication. 10 mg of lysate was incubated rotating overnight with 5 mcg of anti-LSD1 antibody (C69G12), or IgG control, Cell Signaling) at 4^o^C then with Protein A/G+ Agarose Beads (TFS). Samples were then centrifuged, washed, and heated to 70^o^C to reverse crosslink. RNA was isolated with the Nucleospin XS kit (Clontech) including DNAse treatment. RNA was then used for reverse transcription with Superscript IV, then used for qPCR as above. RNA expression was compared to expression in input. RIP experiments were performed in independent triplicate.

### RNA-Seq and data analysis

Total RNA was isolated from cells using the Nucleospin kit as above, with RNA quality assessed using Agilent 2100 bioanalyzer and RNA NanoChip kit (Agilent Technologies, CA). RNA was DNase-treated then subjected to rRNA removal with the Ribo-Zero rRNA removal kit (Illumina). Direction-oriented libraries were constructed from first strand cDNA using ScriptSeq v2 RNA-Seq library preparation kit (Epicentre Biotechnologies, WI). Briefly, 50 ng of rRNA-depleted RNA was fragmented and reverse transcribed using random primers containing a 5’ tagging sequence, followed by 3’end tagging with a terminal-tagging oligo to yield di-tagged, single-stranded cDNA. Di-tagged cDNA was purified via magnetic beads then amplified by limit-cycle PCR using primer pairs that anneal to tagging sequences and add adaptor sequences required for sequencing cluster generation. Amplified RNA-seq libraries were purified using AMPure XP System (Beckman Coulter). Library quality was determined via Agilent 2200 Tapestation using High Sensitivity D1000 screen tape, and quantified by Qubit flourometer with dsDNA BR assay (Thermo Fisher Scientific). Paired-end 150 bp sequence reads were generated using the Illumina HiSeq 4000 platform.

A minimum of 69 million paired-end 150 bp RNA-Seq reads were generated for each sample (range 69–111 million). Each sample was aligned to the GRCh38.p5 assembly of the *H.Sapiens* reference from NCBI (http://www.ncbi.nlm.nih.gov/assembly/GCF_000001405.31/) using version 2.5.0c of the RNA-Seq aligner STAR (http://bioinformatics.oxfordjournals.org/content/early/2012/10/25/bioinformatics.bts635). Transcript features were identified from the GFF file provided with with the GRCh38.p5 assembly and raw coverage counts were calculated using HTSeq (http://www-huber.embl.de/users/anders/HTSeq/doc/count.html). The raw RNA-Seq gene expression data was normalized and post-alignment statistical analyses performed using *DESeq2* (http://genomebiology.com/2014/15/12/550) and custom analysis scripts written in R. Comparisons of gene expression and associated statistical analysis were made between different conditions of interest using the normalized read counts. All fold change values are expressed as test condition/control condition, where values less than one are denoted as the negative of its inverse. Transcripts were considered significantly differentially expressed using a 10% false discovery rate (DESeq2 adjusted *p* value <= 0.1) and a fold-change cut-off of 2 between the control and test samples. Complete dataset will be accessioned to GEO. Annotated gene lists of differentially expressed features are available in the Supplementary Data and were used for analysis of statistical overrepresentation of biological pathways using PANTHER (45), overrepresentation test release 20160715, GO database release 2016–11–30.

### Chromatin Immunoprecipitation-Sequencing (ChIP-Seq)

ChIP-Seq was performed as described(46). Chromatin was crosslinked with fresh 1% formaldehyde for 4 minutes and sheared by the Covaris ME220 for 12 minutes. Anti-H3K4Me1 (ab8895) and anti-H3K4Me2 (ab32356, AbCam) antibodies were used with M-280 Sheep Anti-Rabbit IgG Dynabeads™ (TFS) for ChIP. Isolated chromatin was amplified with the Kapa Library Amplification Kit (Kapa Biosystems) and samples analyzed by HiSeq Illumina Genome Analyzer. Peak calling was performed using the USeq software. Complete data analysis of ChIP-Seq data is included in the supplementary data.

### Statistical Analysis

All statistical analyses excluding the RNA-Seq data were completed using the GraphPad Prism 7 software (GraphPad Software, Inc.). Where appropriate, the two-tailed Student’s t-test was used to calculate significant differences between comparison groups in the experiments above. For multiple comparisons, one-way ANOVA was used with the Bonferroni correction for multiple comparisons against a single control.

## ACKNOWLEDGEMENTS

This work was supported by internal funding from the Research Institute at Nationwide Children’s Hospital.

---

**SI1:**
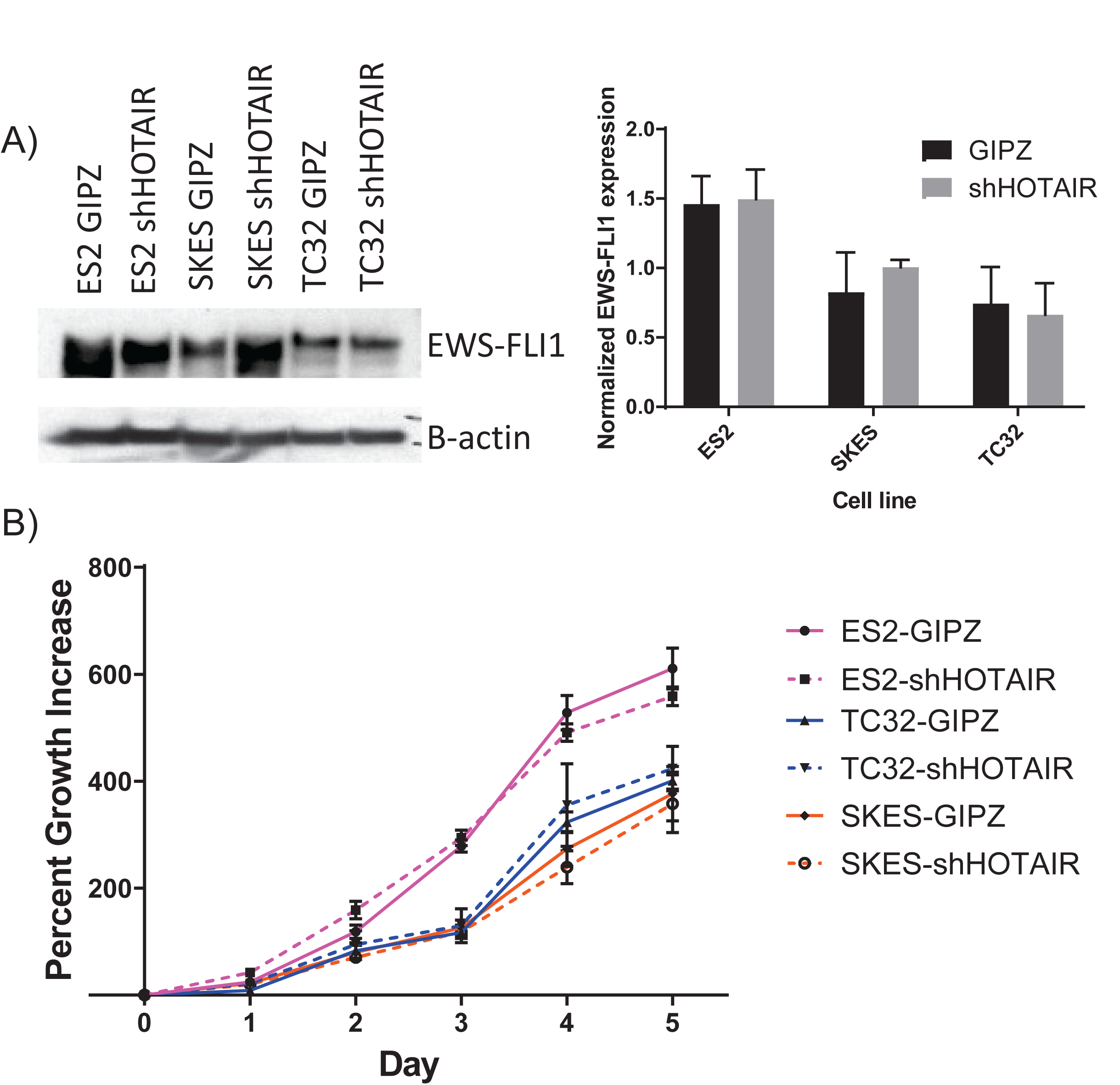
*HOTAIR* affects neither EWS-FLI1 expression nor two-dimensional proliferation in ESFT cell lines. A) Knockdown of *HOTAIR* has no significant effect on EWS-FLI1 protein expression. Protein lysate from ESFT cells expressing shmiRNA against *HOTAIR* or nonsilencing control (GIPZ) was used in western blot for EWS-FLI1 and β-actin. Representative western blots shown on left; normalized expression of EWS-FLI1/actin is shown in graph at right. Three independent western blots were performed for each lineage, error bars SD. B) Proliferation of ESFT cells lines expressing shmiRNA against *HOTAIR* or nonsilencing control (GIPZ). Knockdown of *HOTAIR* does not significantly impact two-dimensional proliferation.

**SI2:**
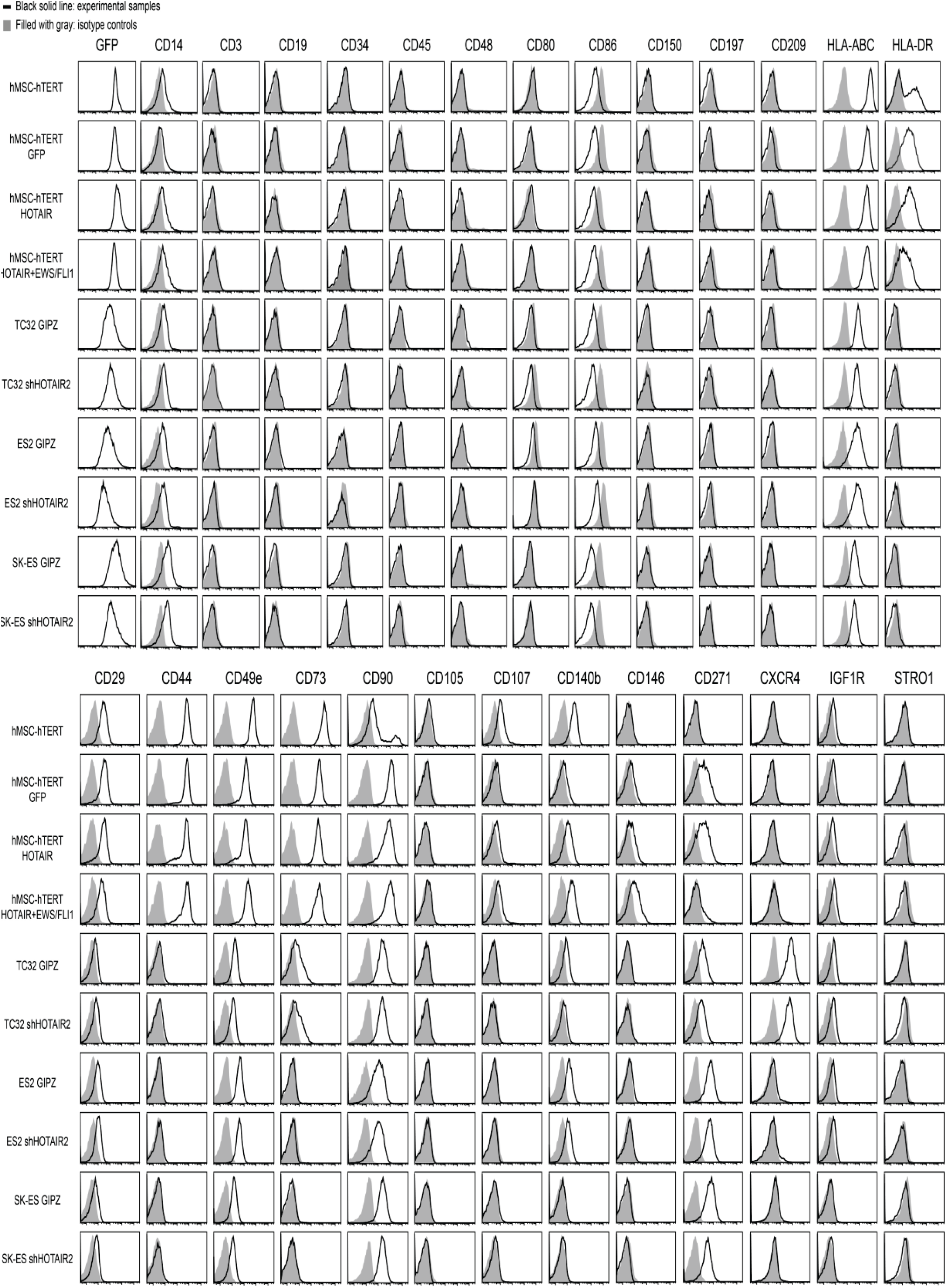
Flow cytometry analysis of hTERT-hMSC with overexpression of *HOTAIR, HOTAIR* and EWS-FLI1, or GFP control, and of ESFT cell lines expressing shmiRNA against *HOTAIR* or nonsilencing control (GIPZ). hTERT-hMSC cells all show similar expression patterns, including expression of the MSC markers CD73 and low expression of hematopoietic lineage markers. The ESFT shHOTAIR variants show similar expression to their matched GIPZ control, including unchanged expression of CD73 and CD90.

**SI3:**
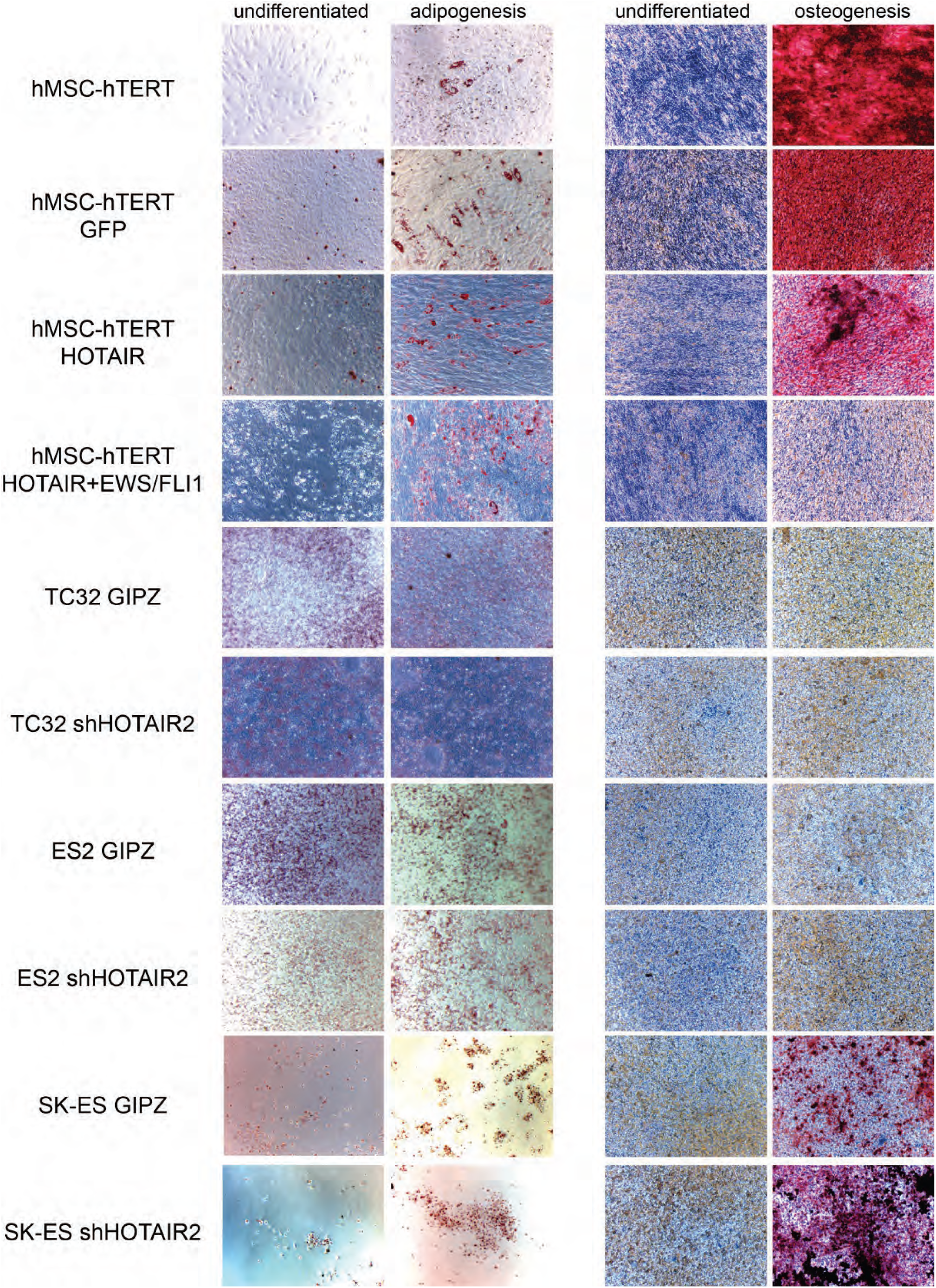
Differentiation analysis of hTERT-hMSC with overexpression of *HOTAIR, HOTAIR* and EWS-FLI1, or GFP control, and of ESFT cell lines expressing shmiRNA against *HOTAIR* or nonsilencing control (GIPZ). In the hTERT-hMSC variants, cells are still able to be differentiated into adipogenic (left) and osteogenic (right) lineages, although the hTERT-hMSC-*HOTAIR-EWS-FLI1* cells have reduced osteogenic capacity. In the ESFT cells, all cells maintain capacity for adipogenic differentiation; none of the TC32 or ES2 cells have osteogenic capacity, but both SK-ES GIPZ and shHOTAIR cells maintain osteogenic differentiation.

**SI4:**
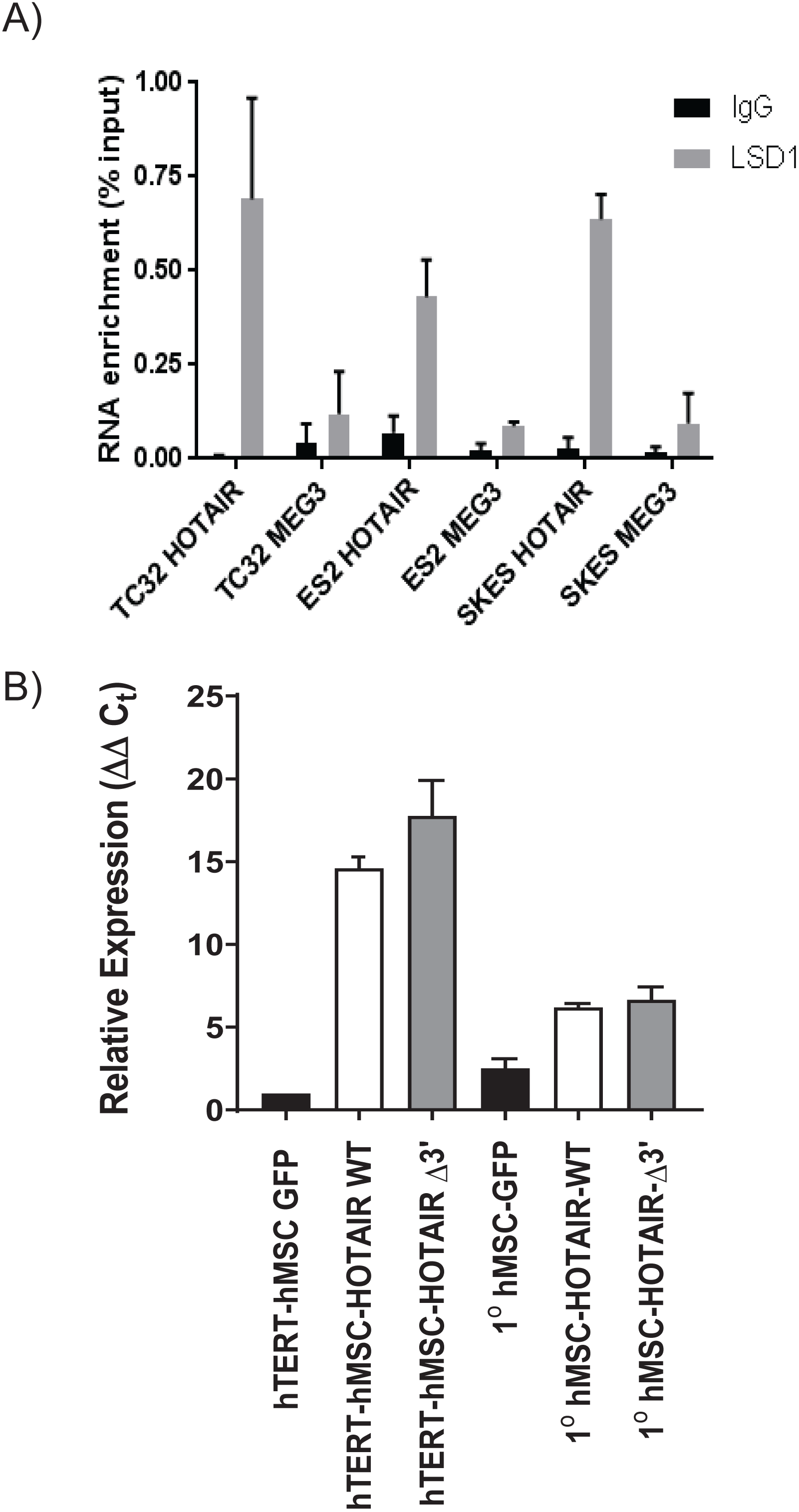
*HOTAIR* interaction with epigenetic complexes, and expression of mutant *HOTAIR* variants. A) RNA immunoprecipitation of LSD1 in the ESFT cell lines TC32, ES2, and SK-ES show enrichment for wild-type *HOTAIR* as compared to IgG control, with no enrichment of *MEG3* as a control. P<0.01 for all enrichment of *HOTAIR* vs IgG and vs *MEG3*. B) Quantitation of overexpression of wildtype *HOTAIR* and mutant variants in hTERT-hMSCs and 1^o^ hMSCs, as tested by RT-qPCR for a region of *HOTAIR* found in all variants. The hTERT-hMSC *HOTAIR* variants are not significantly different in their expression (p=0.17), not are the 1^o^ hMSC variants significantly different in their expression (p=.32), as compared by student's t-test.

**SI5:**
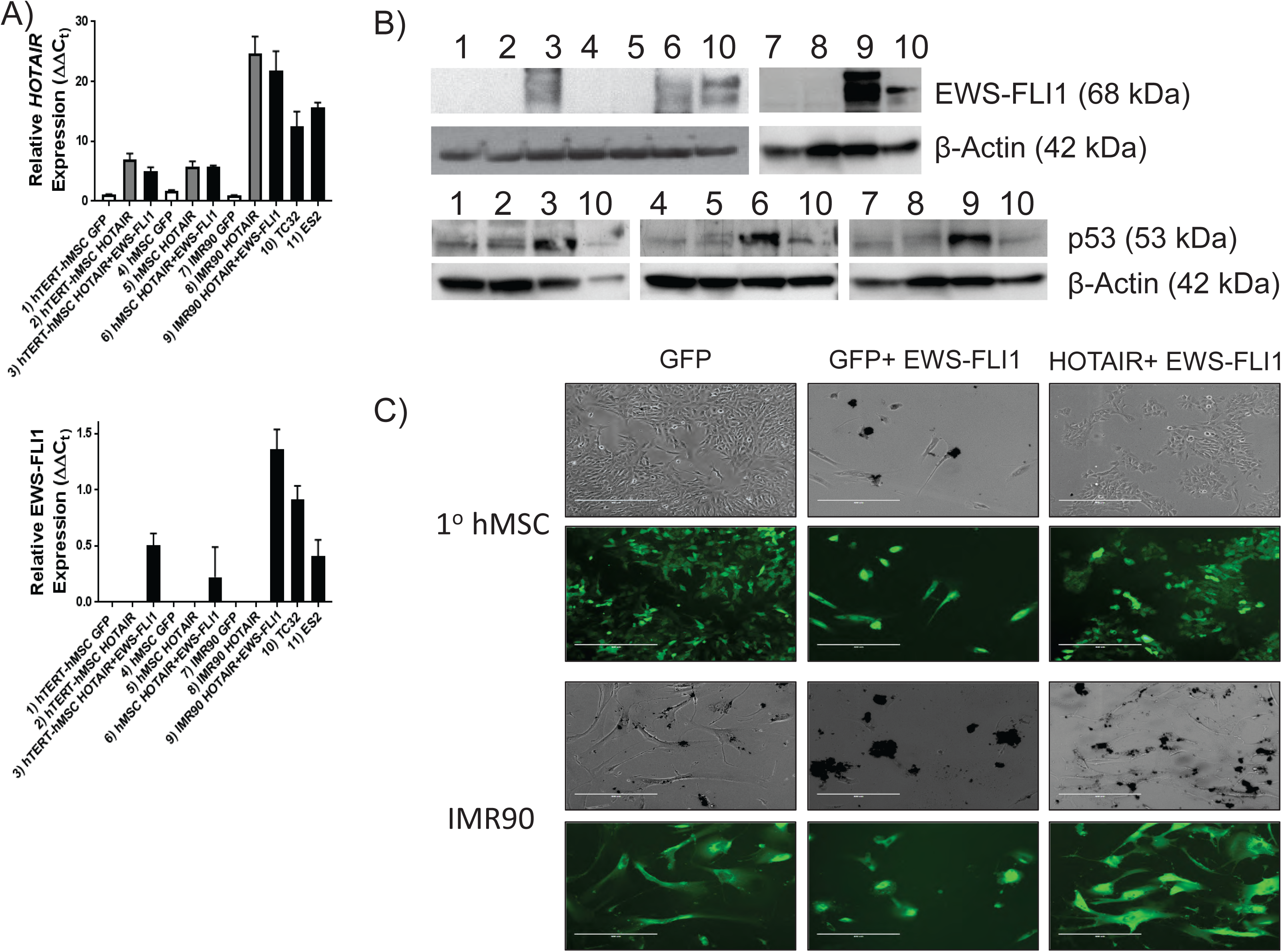
Stable *HOTAIR* expression allows for subsequent stable EWS-FLI1 expression in hMSCs and IMR90 cells. 1) mRNA expression of *HOTAIR* (top) and *EWS-FLI1* (bottom) in hTERT-hMSCs, 1^o^ hMSCs, and IMR90 cells with either GFP alone, *HOTAIR*, or *HOTAIR* then *EWS-FLI1* exogenously expressed, with comparisons to the ESFT cell lines TC32 and ES2. 2) Western blot analysis of EWS-FLI1 and p53 expression in these cell line models, with β-actin expression as controls. Number coding corresponds to the sample numbers in figure SI 5A. EWS-FLI1 is stably expressed in the hTERT-hMSC (#3), 1^o^ hMSC (#6) and IMR90 (#9) cell lines with stable *HOTAIR* and *EWS-FLI1* expression. p53 expression is stably demonstrated in all lineages, with upregulation in the *HOTAIR-EWS-FLI1* cell lines. TC32 cell lysate is used as comparison (#10). Note: the same western blot of IMR90 cell variants was used to evaluate expression of EWS-FLI1, p53, and β-actin (samples 7–10), and the same β-actin blot is shown for normalization of EWS-FLI1 (top) and p53 (below). C) Representative photomicrographs and fluroscent photomicrographs of 1^o^ hMSCs and IMR90 cells with exogenous expression of GFP alone (left), GFP+EWS-FLI1 (middle) or *HOTAIR* followed by EWS-FLI1 (right). Pictures taken 2 weeks after viral infection/expression and stable expression, as represented in figures A and B. In both cell lines, GFP expression alone was tolerated without toxicity, EWS-FLI1 expression alone induced cell death, but initial stable *HOTAIR* expression allowed subsequent stable EWS-FLI1 expression and cell viability.

**SI6:**
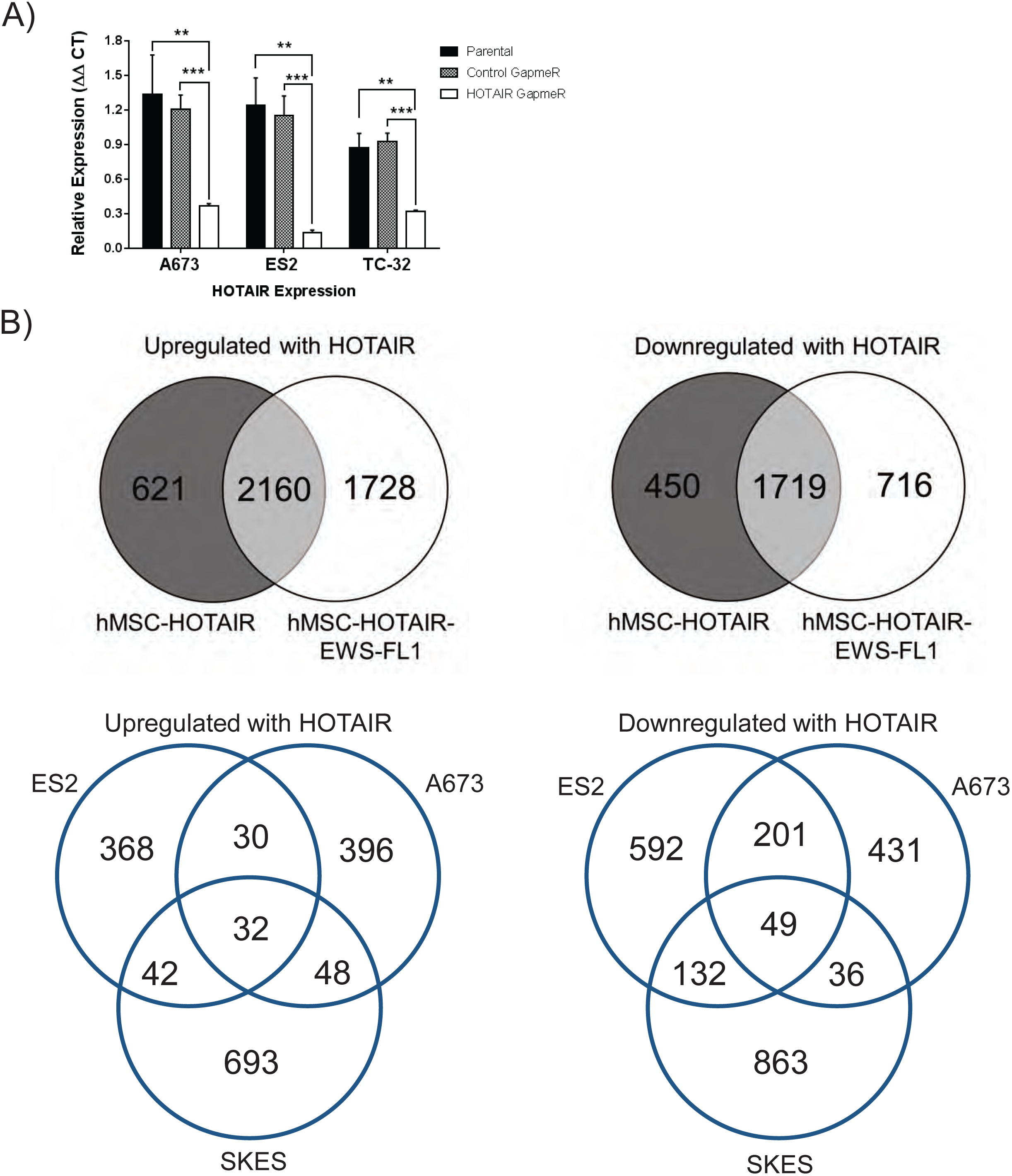
RNA-Seq analysis summaries. A) *HOTAIR* expression in ESFT cell lines treated with nonsilencing GapmeR control or HOTAIR-targeting GapmeR. All cell lines had repression of *HOTAIR* to 20–30% of baseline. C) Numbers of common and disparate genes differentially expressed (adjusted p-value ≤0.05 and at least two-fold relative change) in different cell line models. Upper panels are genes differentially expressed in hTERT-hMSC-HOTAIR or *hTERT-hMSC-HOTAIR-EWS-FLI1* versus GFP control. Lower panels are *HOTAIR-* GapmeR-treated ESFT cells vs nonsilencing control. Left panels identify genes whose expression directly correlates with *HOTAIR* expression; right panels identify genes whose expression inversely correlates with *HOTAIR* expression. Proportionately more genes were found to be upregulated than downregulated with overexpression of *HOTAIR* in hTERT-hMSC cells, with a majority of genes similarly affected regardless of *EWS-FLI1* expression. In contrast, more genes were found to be downregulated than upregulated in correlation with *HOTAIR* expression in ESFT cells.

**SI7:**
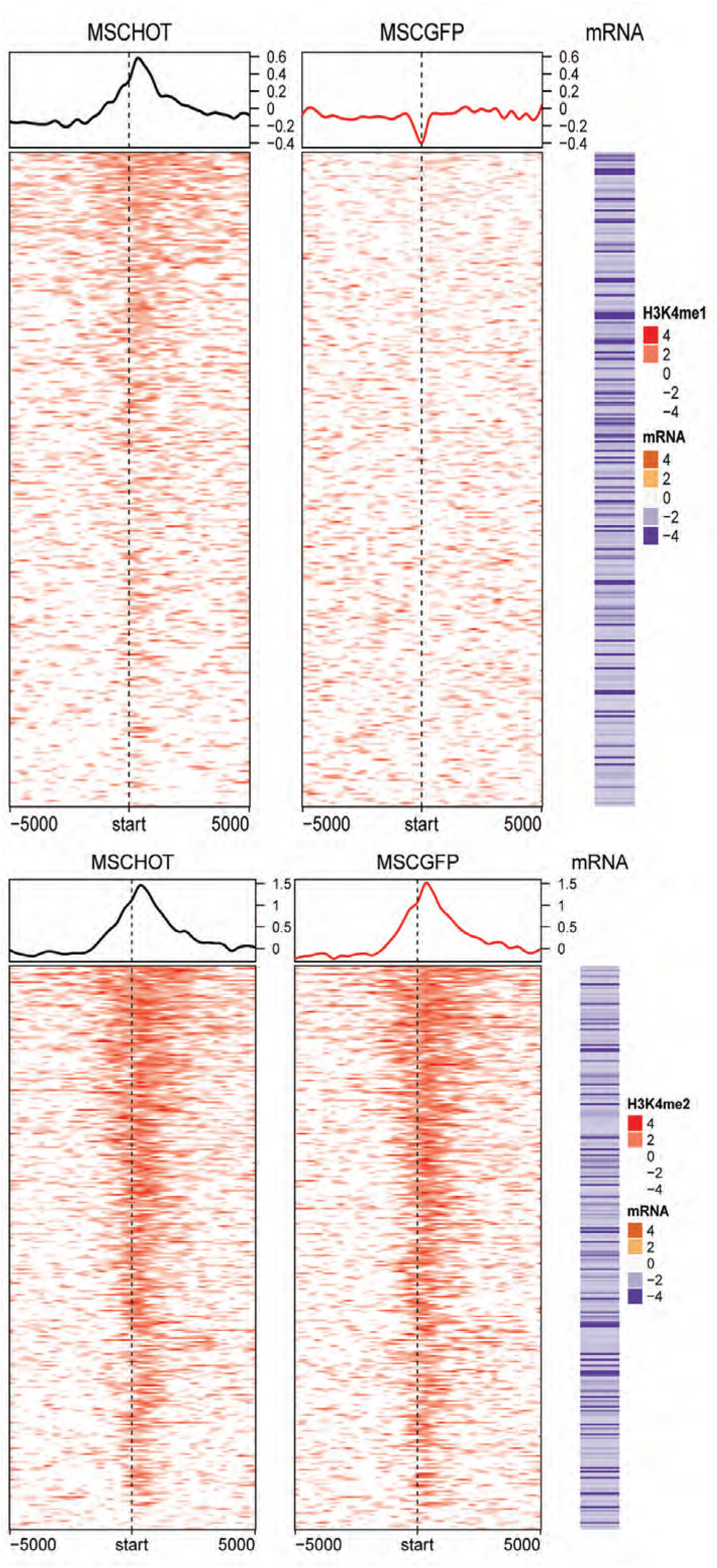
Chromatin Immunoprecipitation and Sequencing (ChIP-Seq) identified genomic regions with differential H3K4Me1 and H3K4Me2 binding in hTERT-hMSC-HOTAIR cells (“MSCHOT”, left) as compared to hTERT-hMSC-GFP (“MSCGFP”, right), with FDR<0.05. A) Profiles of HOTAIR-induced H3K4me1 modification around TSS of HOTAIR-repressed genes. Each row corresponds to HOTAIR-repressed genes (286 genes). Color for H3K4me1 profile represents log_2_ ratio of ChIP to input control. Positive (red) represents enrichment of H3K4me1. Mean H3K4me1 profile in each cell line are plotted on top. The corresponding mRNA expression (log2 ratio) is shown on the right panel. Negative (purple) represents decreased expression in hTERT-hMSC-HOTAIR cells compared to hTERT-hMSC-GFP controls (i.e. genes that are repressed by HOTAIR). H3K4Me1 was differentially bound adjacent to the TSS of these *HOTAIR-repressed* genes in the context of *HOTAIR* expression in hTERT-hMSC-HOTAIR cells as compared to control. B) Profiles of HOTAIR-repressed H3K4me2 binding in hTERT-hMSC-HOTAIR and hTERT-hMSC-GFP around TSS of HOTAIR-repressed genes. Each row corresponds to HOTAIR-repressed genes (287 genes). Color for H3K4me2 profile represents log_2_ ratio of ChIP to input control. Positive (red) represents enrichment of H3K4me2. Mean H3K4me2 profile in each cell line are plotted on top. The corresponding mRNA expression (log2 ratio) is shown on the right panel. Negative (purple) represents decreased expression in hTERT-hMSC-HOTAIR compared to hTERT-hMSC-GFP (i.e. genes that are repressed by HOTAIR). H3K4Me2 was differentially bound to adjacent to the TSS of HOTAIR-repressed genes in the context of low *HOTAIR* expression in hTERT-hMSC-GFP cells as compared to the hTERT-hMSC-HOTAIR cells (i.e., low *HOTAIR* expression correlates with increased H3K4Me2 binding and increased expression of these correlated genes). Differential H3K4Me2 expression is significant for each gene identified (FDR <0.05 by MACS2 analysis).

**SI8:**
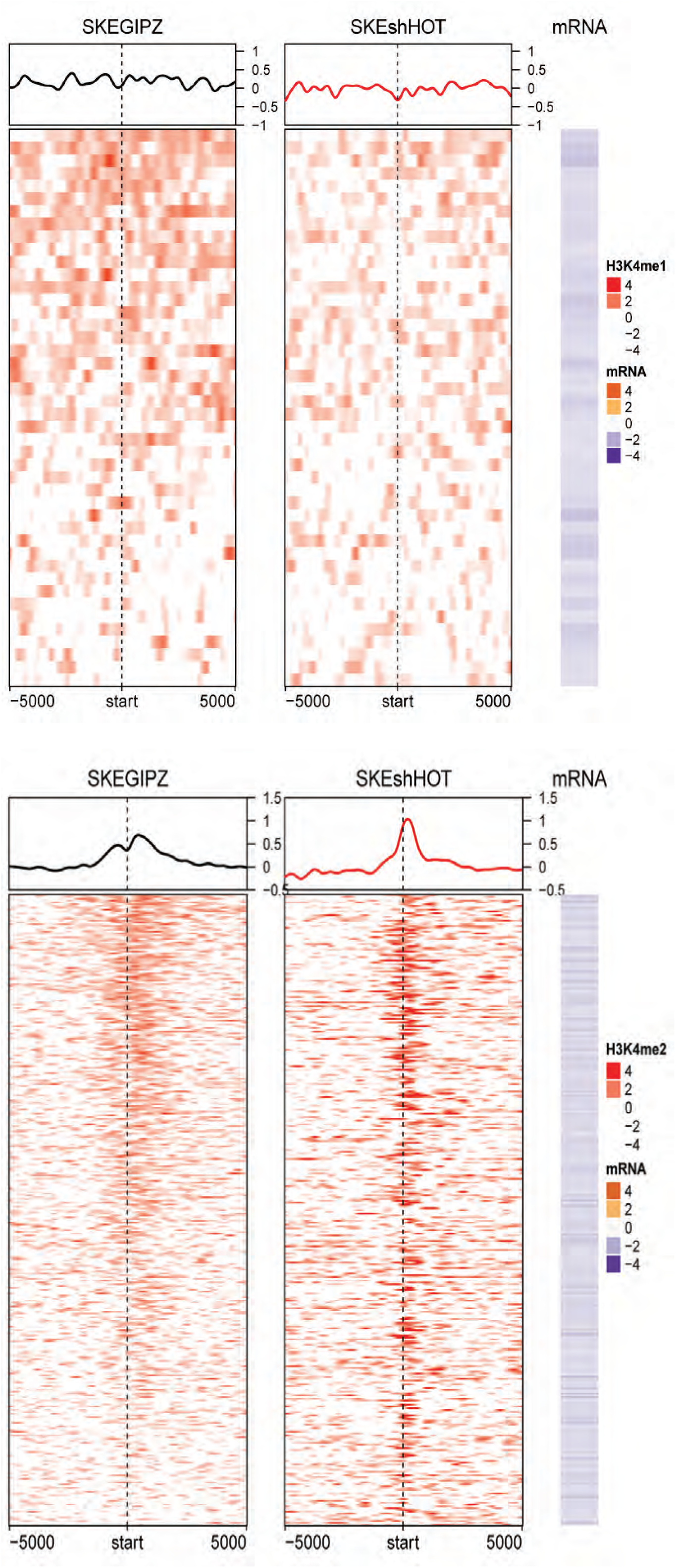
Chromatin Immunoprecipitation and Sequencing (ChIP-Seq) identified genomic regions with differential H3K4Me1 and H3K4Me2 binding in SKES-pGIPZ (“SKEGIPZ”, left) as compared to SKES-shHOTAIR (“SKEshHOT,” right), with FDR<0.05. A) Profiles of HOTAIR-induced H3K4me1 modification around TSS of HOTAIR-repressed genes. Each row corresponds to HOTAIR-repressed genes (44 genes). Color for H3K4me1 profile represents log_2_ ratio of ChIP to input control. Positive (red) represents enrichment of H3K4me1. Mean H3K4me1 profile in each cell line are plotted on top. The corresponding mRNA expression (log2 ratio) is shown on the right panel. Negative (purple) represents decreased expression in SKES-pGIPZ cells compared to SKES-shHOTAIR controls (i.e. genes that are repressed by HOTAIR). H3K4Me1 was differentially bound adjacent to the TSS of these HOTAIR-repressed genes in the context of *HOTAIR* expression SKES-pGIPZ cells as compared to SKES-shHOTAIR. B) Profiles of HOTAIR-repressed H3K4me2 binding in SKES-pGIPZ and SKES-shHOTAIR around TSS of HOTAIR-repressed genes. Each row corresponds to HOTAIR-repressed genes (287 genes). Color for H3K4me2 profile represents log_2_ ratio of ChIP to input control. Positive (red) represents enrichment of H3K4me2. Mean H3K4me2 profile in each cell line are plotted on top. The corresponding mRNA expression (log2 ratio) is shown on the right panel. Negative (purple) represents decreased expression in SKES-pGIPZ compared to SKES-shHOTAIR (i.e. genes that are repressed by HOTAIR). H3K4Me2 was differentially bound to adjacent to the TSS of HOTAIR-repressed genes in the context of loss of *HOTAIR* expression in SKES-shHOTAIR cells as compared to SKES-pGIPZ cells (i.e., loss of *HOTAIR* expression correlates with increased H3K4Me2 binding and increased expression of these correlated genes).

**SI9: Complete ChIP-Seq analysis summary.**

## H3K4me1 enrichment in ES2 in relation to HOTAIR expression

81,362 and 55,991 H3K4me1 broad peaks were identified at FDR < 0.05 using MACS2 in ES2GIPZ and ES2shHOTAIR, respectively. Peaks in each cell line were normalized to their corresponding input samples. 1,410 genes were found to be differentially regulated by HOTAIR at FDR < 0.05 (ES2Control vs ES2HOTAIRGapmer).

**Figure 1.**
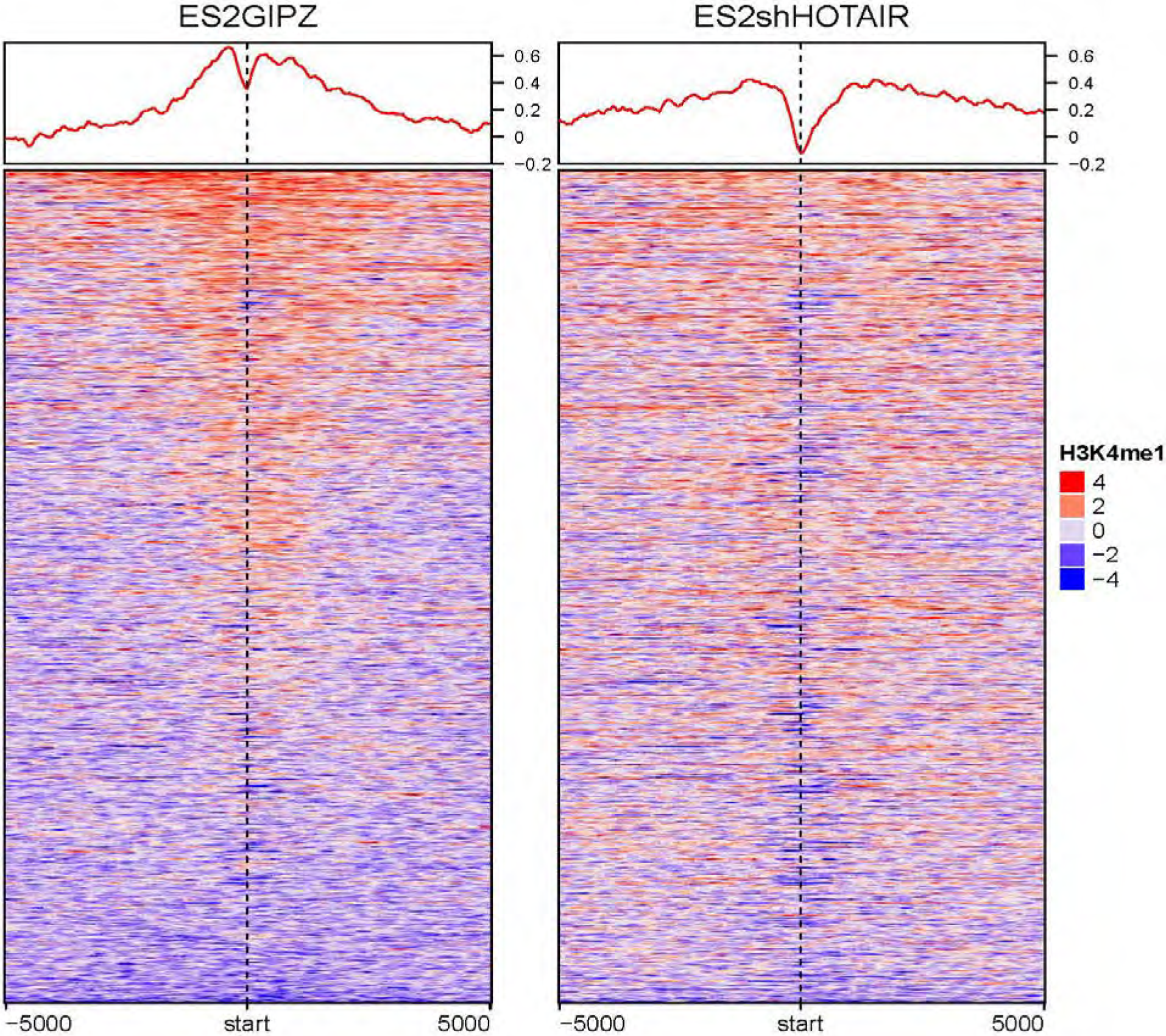
Heatmap of H3K4me1 profiles in ES2GIPZ and ES2shHOTAIR plotted ± 5000-bp of HOTAIR-regulated genes in ES2 (FDR < 0.05). Color represents log_2_ ratio of normalized read counts over input. Mean signals in each cell line are plotted on top. As shown in the heatmap, HOTAIR expression induces increased H3K4me1 binding around TSS of HOTAIR-regulated genes. (hm_ES2_H3K4M1_norm_diff_GIPZ_shHOT.pdf)

**Figure 2.**
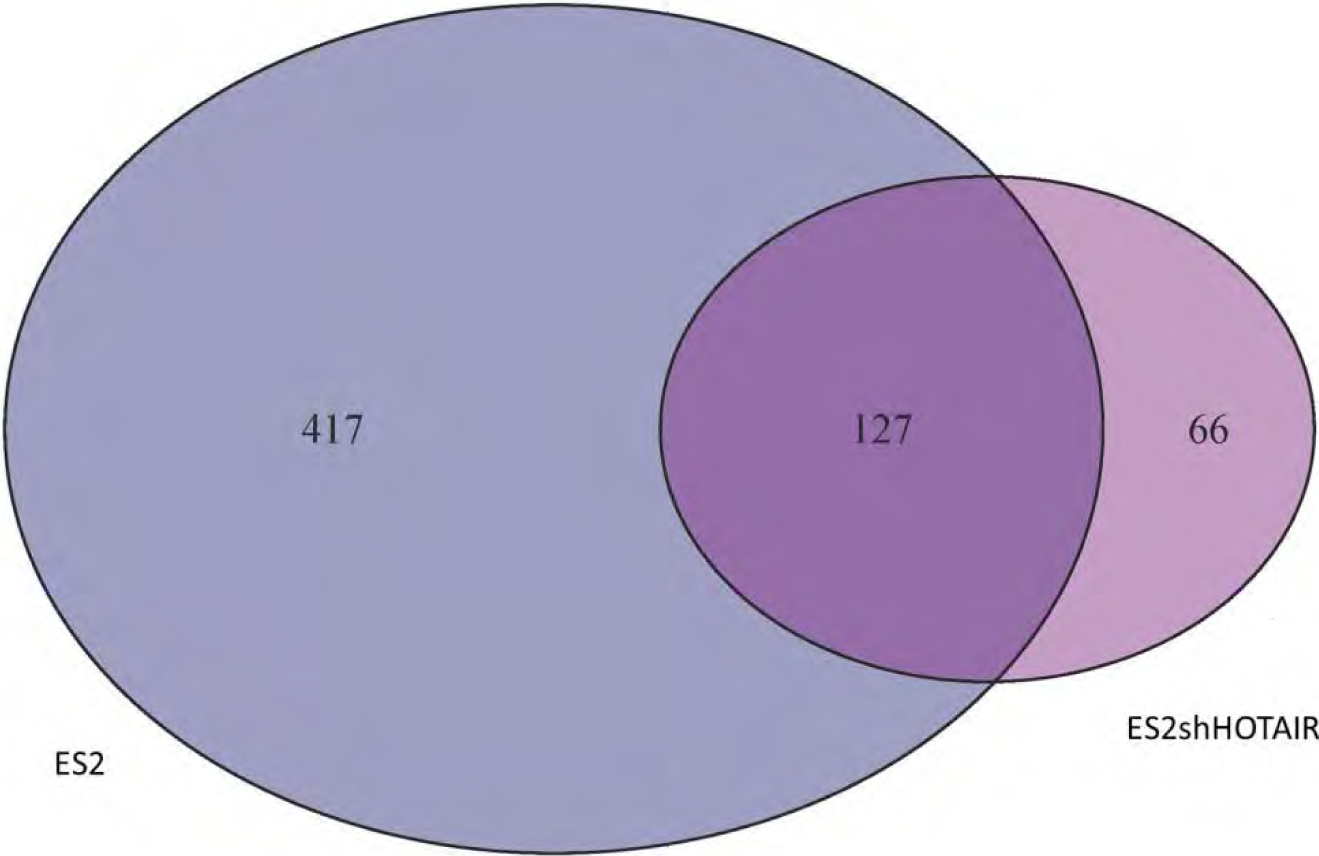
36,202 H3K4me1 binding peaks were found to be increased in ES2GIPZ and 12,333 peaks were found to be increased in ES2shHOTAIR. Regions with increased binding in ES2shHOTAIR were adjacent to 3,648 genes of which 193 is HOTAIR-regulated genes (40 up- and 153 down-regulated by HOTAIR). Regions with increased binding in ES2GIPZ were adjacent to 10,556 of which 544 is HOTAIR-regulated genes (141 up- and 403 down-regulated by HOTAIR). Venn diagram shows overlap between number of HOTAIR-regulated genes adjacent to significant increased H3K4me1 binding in ES2 (dark blue) and significant increased H3K4me1 binding in ES2shHOTAIR (magenta). Note that multiple differential binding may be adjacent to the same gene. venn_ES2GIPZvsshHOT_H3K4M1.pdf.

**Figure 3.**
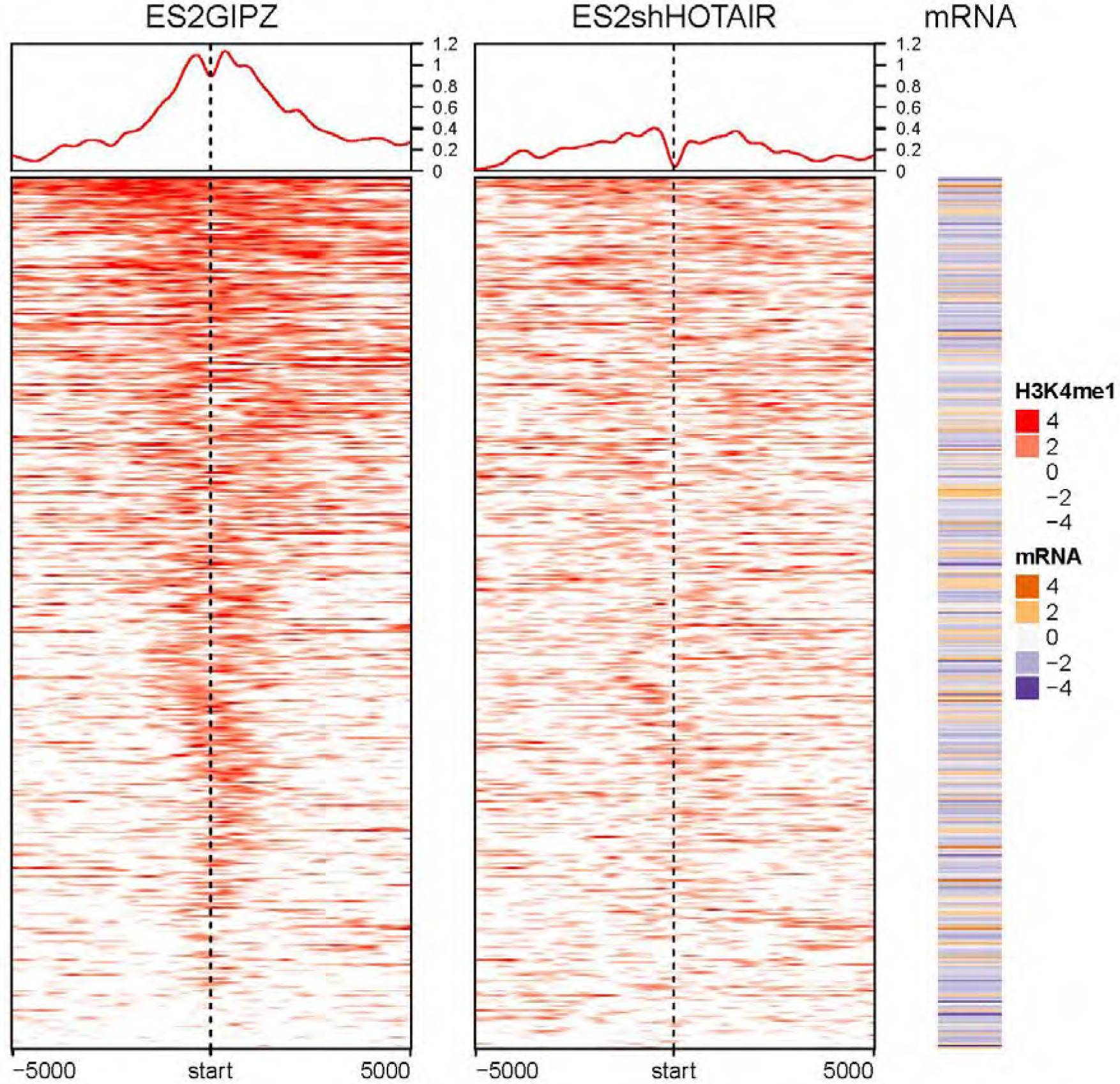
Plot of significantly increased H3K4me1 binding in ES2GIPZ compared to ES2shHOTAIR around TSS of HOTAIR-regulated genes (417 genes in venn diagram on Figure 2). Color for H3K4me1 ChIP-seq represents log_2_ ratio of ChIP to input samples. Positive (red) represents enrichment of H3K4me1. Mean H3K4me1 signals in each cell line are plotted on top. The corresponding mRNA expression (log2 ratio) is shown on the right panel. Negative (purple) represents decreased expression in ES2Control compared to ES2shHOTAIR (i.e. genes that are repressed by HOTAIR). “hm_ES2_H3K4M1_norm_GIPZenrich_shHOT.pdf”

**Figure 4.**
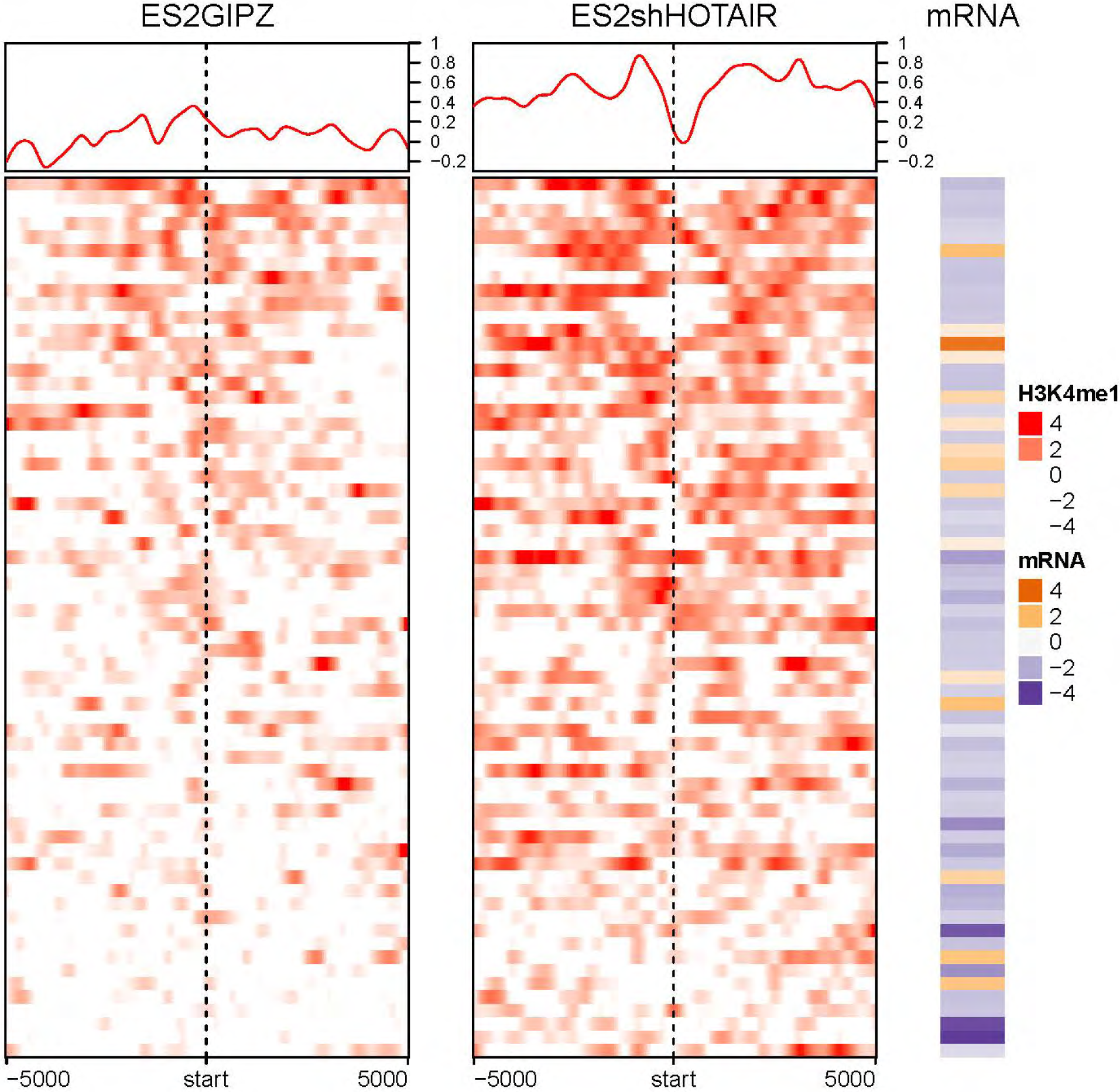
Plot of H3K4me1 significantly increased binding in ES2shHOTAIR compared to ES2GIPZ around TSS of HOTAIR-regulated genes (66 genes in venn diagram on Figure 2). Color represents log2 ratio of ChIP vs. input samples. Mean signals in each cell line are plotted on top. The expression of adjacent genes is plotted on the right panel (log_2_ ratio of ES2HOTAIRGapmer/ ES2Control). Negative (purple) represents decreased expression in ES2Control compared to ES2shHOTAIR (i.e. genes that are repressed by HOTAIR). hm_ES2_H3K4M1_norm_GIPZ_shHOTenrich.pdf

**Figure 5.**
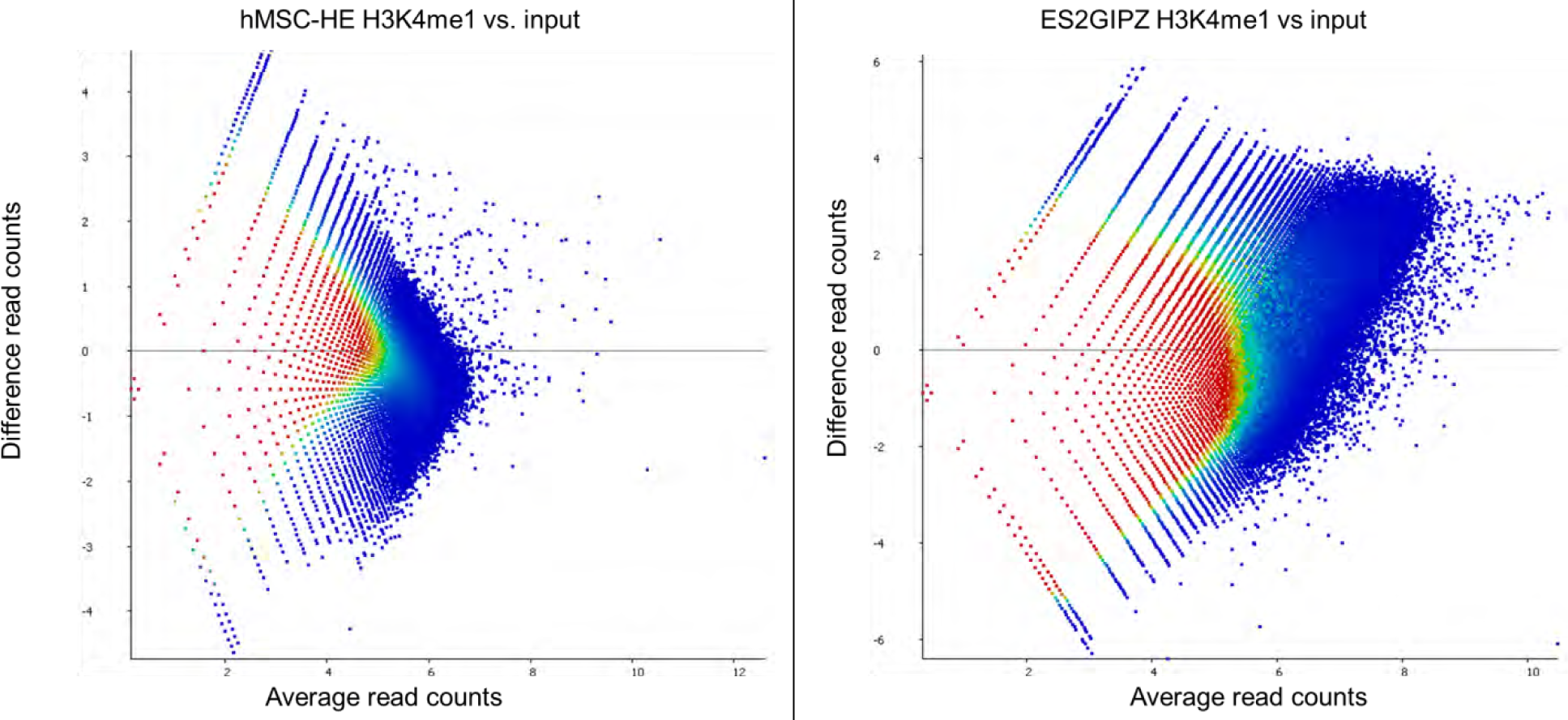
Plot of read counts difference between H3K4me1 and input in hMSC-HE samples (left panel) and in ES2GIPZ samples (right panel). As shown by the heatmap, there are more H3K4me1 enriched regions in ES2GIPZ compared to hMSC-HE. Color represents density of regions from high (red) to low (blue).

## H3K4me2 bindings in ES2GIPZ vs. ES2shHOTAIR

**Figure 6.**
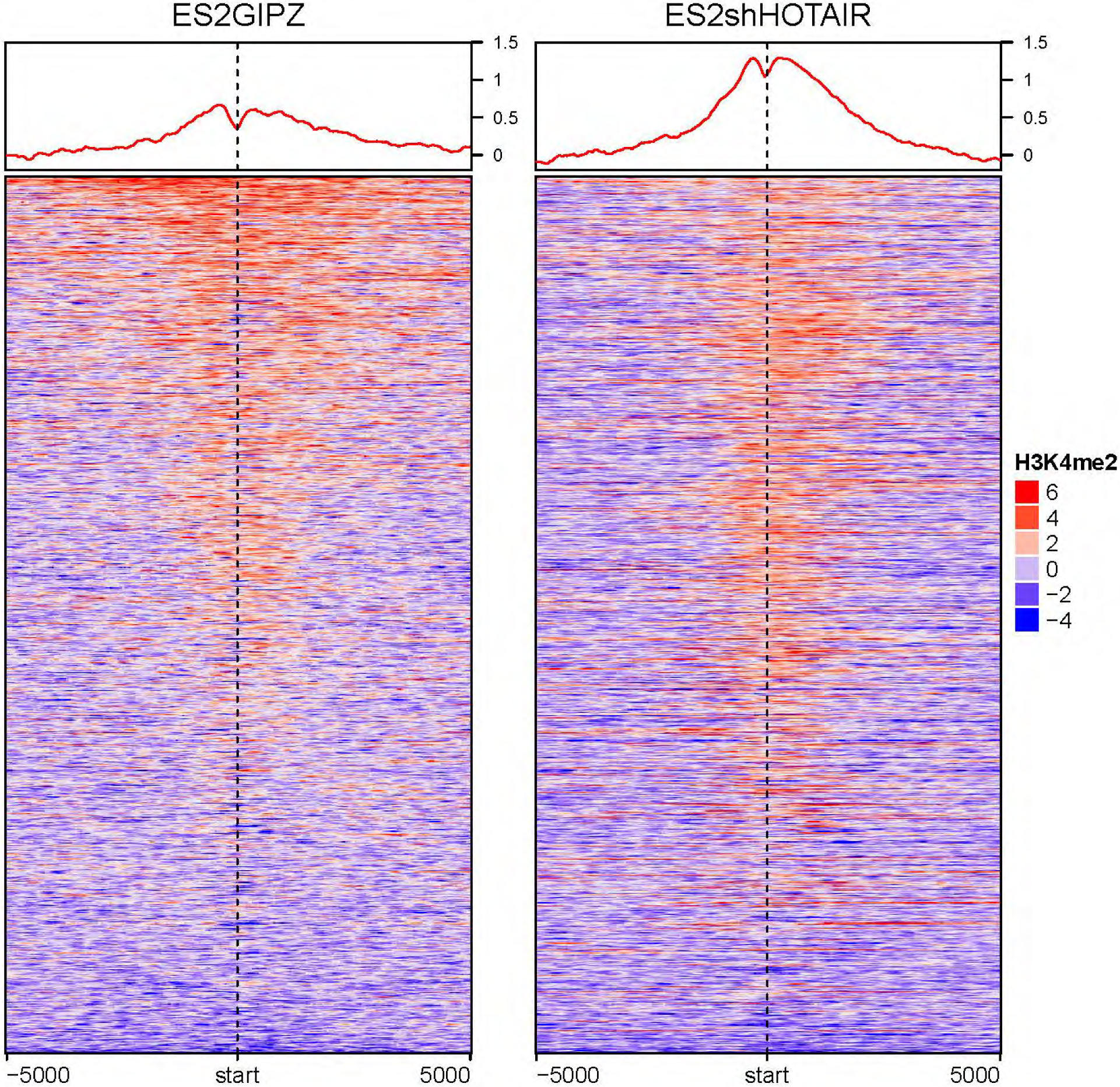
Heatmap of HBK4me2 profiles in ES2GIPZ and ES2shHOTAIR plotted ± 5000-bp of HOTAIR-regulated genes in ES2 (FDR < 0.05). Color represents log_2_ ratio of normalized read counts over input. Mean signals in each cell line are plotted on top. Overall, knockdown of HOTAIR induces increase of H3K4me2 binding around TSS of HOTAIR-regulated genes. (hm_ES2_H3K4M2_norm_diff_GIPZ_shHOT.pdf)

## Differential H3K4me2 binding in ES2GIPZ versus ES2shHOTAIR

There are 70,018 regions associated with significant increase of H3K4me2 binding in ES2shHOTAIR compared to ES2GIPz. These regions are adjacent to 17,171 unique genes and 750 of them are HOTAIR-regulated (544 down-regulated and 206 up-regulated by HOTAIR). In contrast, only 3 regions are associated with significant increase of H3K4me2 binding in ES2GIPZ compared to ES2shHOTAIR. These 3 regions are adjacent to 3 unique genes but none of them is HOTAIR-regulated genes.

**Figure 7.**
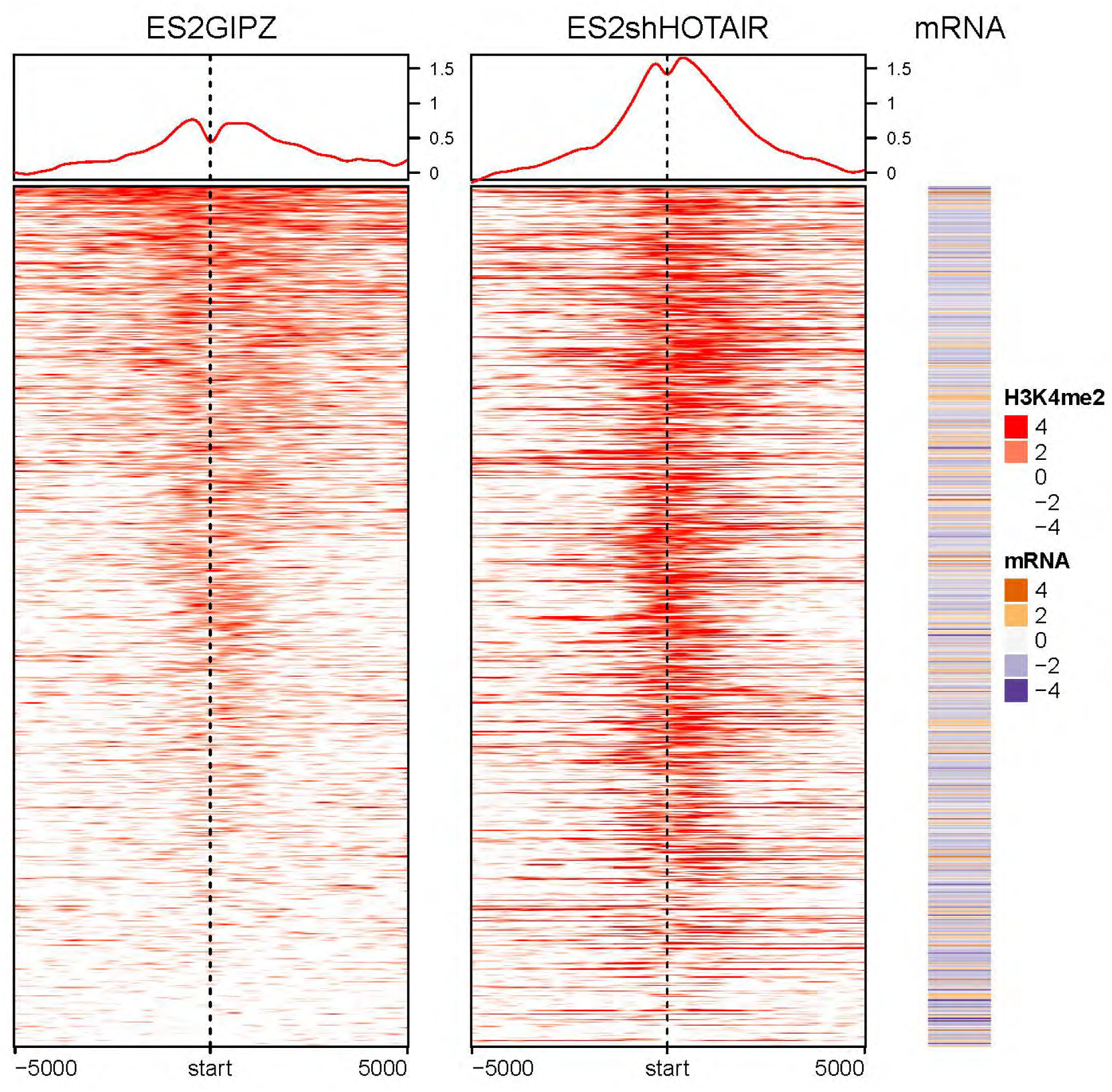
Plot of regions with significant increase of H3K4me2 binding in ES2shHOTAIR around TSS of HOTAIRregulated genes. hm_ES2_H3K4M2_norm_GIPZ_shHOTenrich.pdf

ES2_H3K4M1_norm_GIPZenrich_shHOT.xlsx contains list of regions with gain of H3K4me1 in ES2GIPZ compared to ES2shHOT with its adjacent gene.

ES2_H3K4M2_norm_GIPZ_shHOTenrich.txt contains list of regions with gain of H3K4me2 in ES2shHOTAIR compared to ES2GIPZ with its adjacent gene.

ES2_H3K4M2_norm_GIPZenrich_shHOT.txt contains list of regions with gain of H3K4me1 in ES2GIPZ compared to ES2shHOTAIR with its adjacent gene.

## Assessing batch effects with additional sequencing reads

As of 8/8/2017 there are a total of 53 ChIP-seq runs. 35 ChIP-seq ran previously using high output mode (_1 files), and new sets of sequencing run has 18 ChIP-seq using rapid mode ran on two lanes ( _2 and _3 files).

Read counts (Corrected for total counts) for random 7,000 windows of size 7,000bp in each ChIP-seq runs (53 runs) are evaluated and compared. Duplicated reads are only count as one. Counting and random selection were done using seqmonk (https://www.bioinformatics.babraham.ac.uk/projects/seqmonk/).

**Figure 8.**
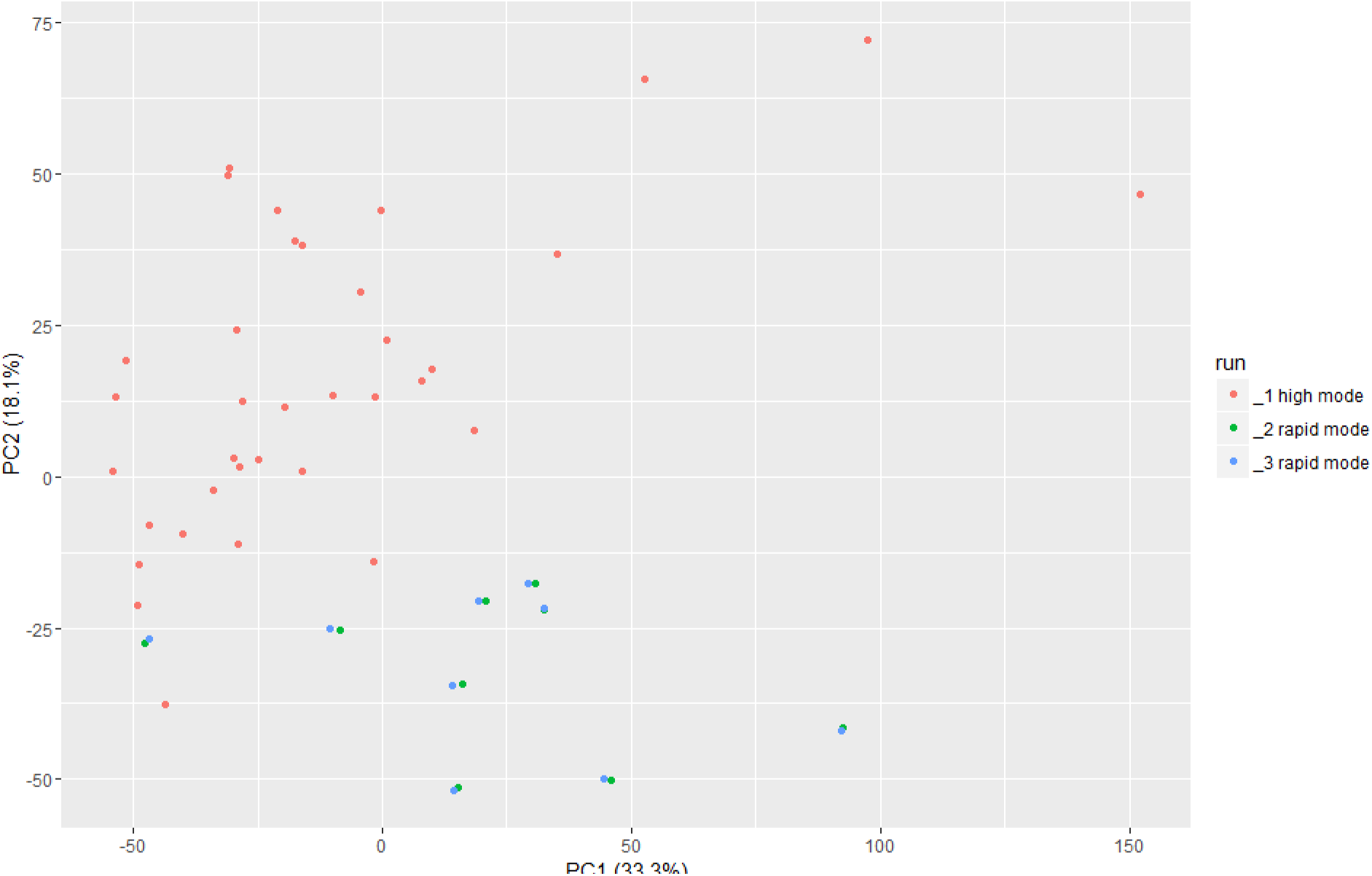
PCA plot suggesting there is a batch effect between rapid mode and high output mode runs. Colors represent different runs of ChIP-seq.

**Figure 9.**
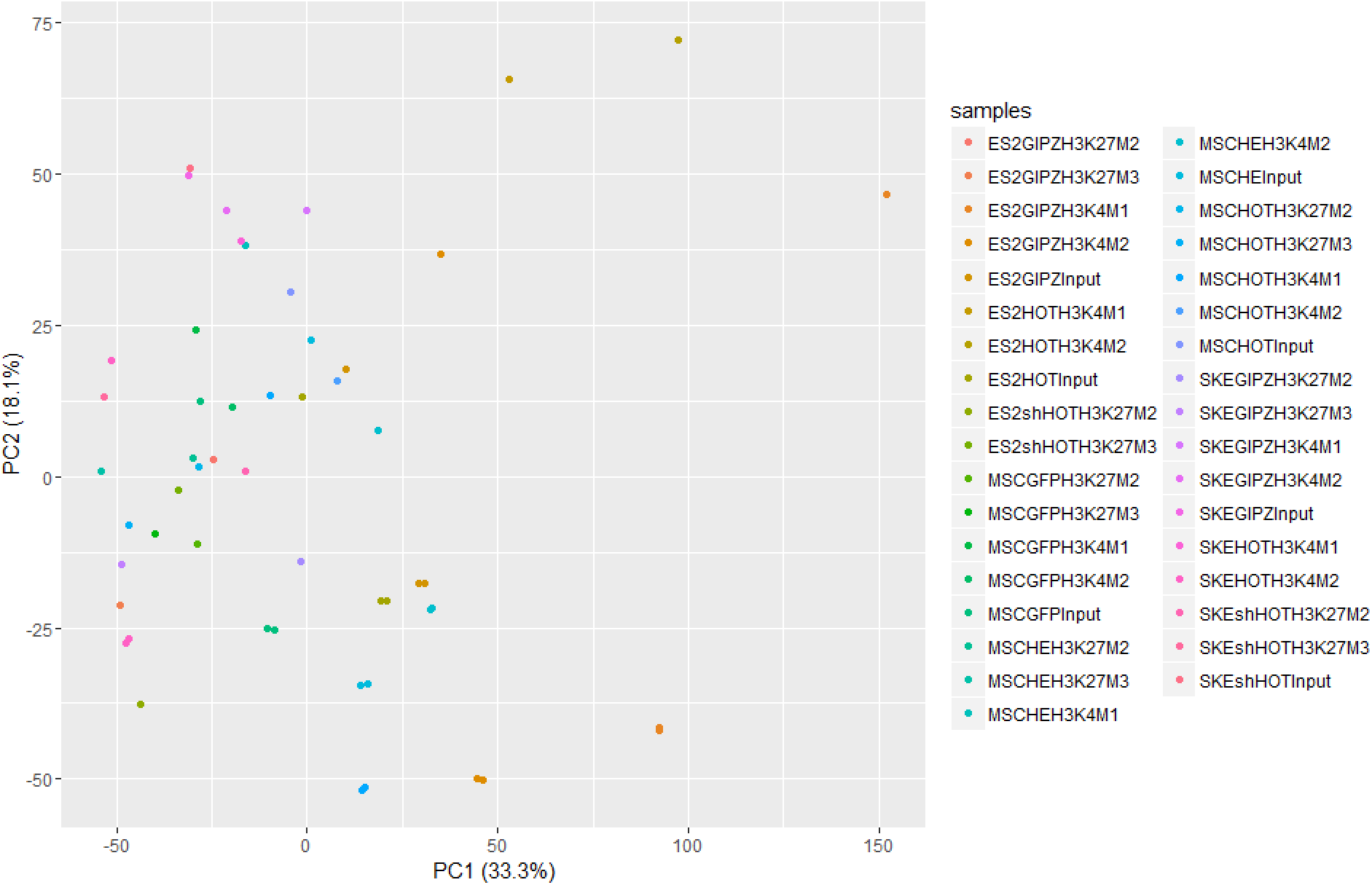
Figure 8 with colors represent the different ChIP-seq sample.

**Figure 10.**
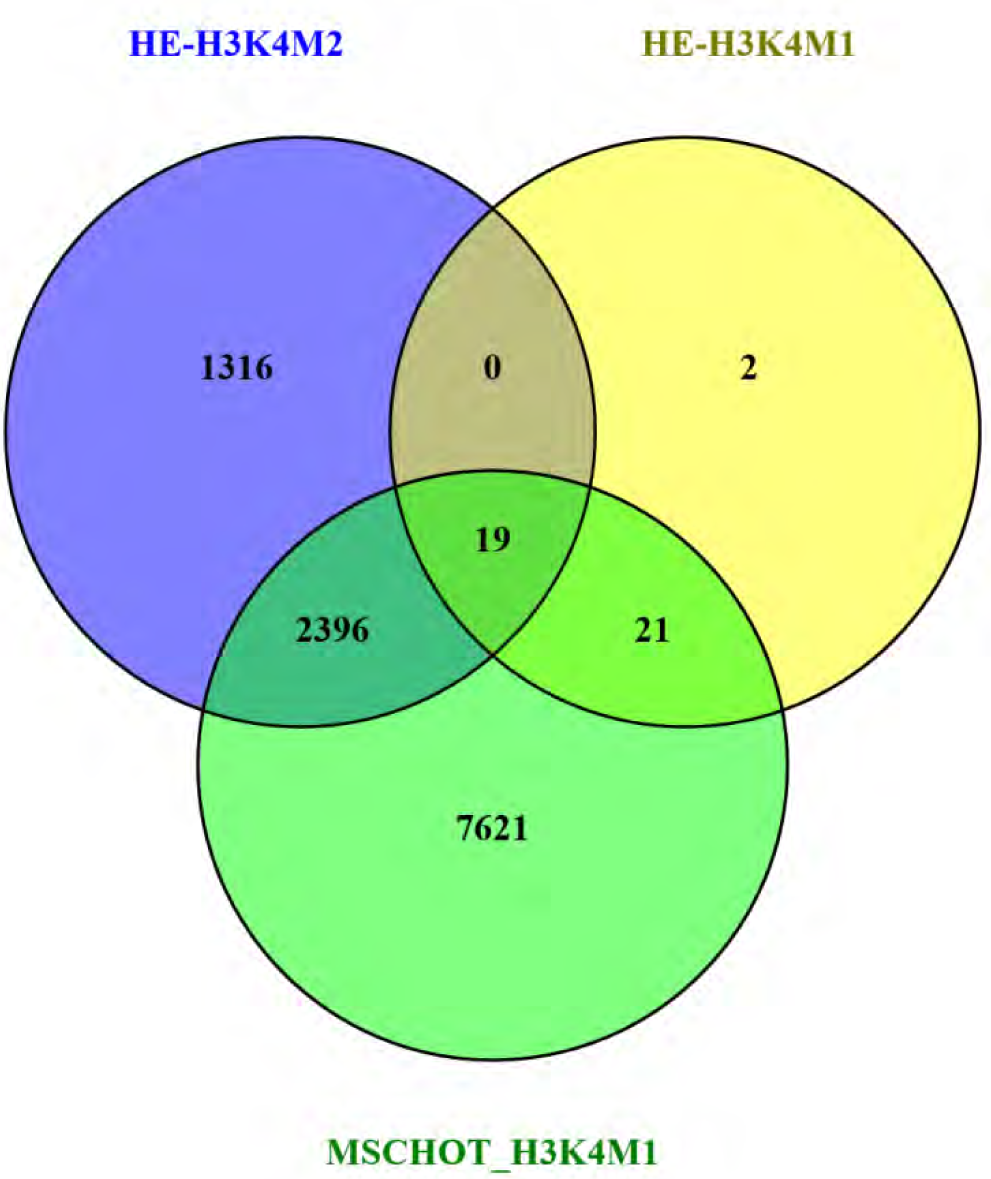
Overlap of genes near H3K4me1 peaks in hMSC-HOTAIR and hMSC-HE and genes near H3K4me2 peaks in hMSC-HE cells. Gene lists are shown in 'venn_gene_HE_HOTH3K4.xlsx'. The genes in HEH3K4me2 and HOTH3K4me1 have more overlap than expected by random chance (p-value < 10^−120^) The genes in HEH3K4me1 and HOTH3K4me1 have more overlap than expected by random chance (p-value < 10^−10^) P-values of overlap were calculated using Fisher's exact test implemented in R package GeneOverlap.

The number of genes in HEH3K4me1 is much less than the others make it looks they don't overlap much. However, there are only 42 genes in HEH3K4me1 and 40 of them overlap with HOTH3K4me1 (95%). While there are 3731 genes in HEH3K4me2 and 2396 of them overlap with HOTH3K4me1 (64%)

## From here on analysis are done on merged reads (from two different runs of ChIP-seq)

### H3K4me1 enrichment in ES2 in relation to HOTAIR expression

95,615 (adjacent to 19,061 unique genes) and 90,201 (adjacent to 17,498 unique genes) H3K4me1 broad peaks were identified at FDR ≤ 0.05 using MACS2 in ES2GIPZ and ES2shHOTAIR, respectively. Peaks in each cell line were normalized to their corresponding input. 1,181 genes were found to be differentially regulated by HOTAIR (FDR ≤ 0.05 and fold-change ≥ 2). Genes regulated by HOTAIR have positive fold-change. 665 and 617 HOTAIR-regulated genes are found adjacent to H3K4me1 peaks in ES2GIPZ and ES2shHOTAIR, respectively.

**Figure 11.**
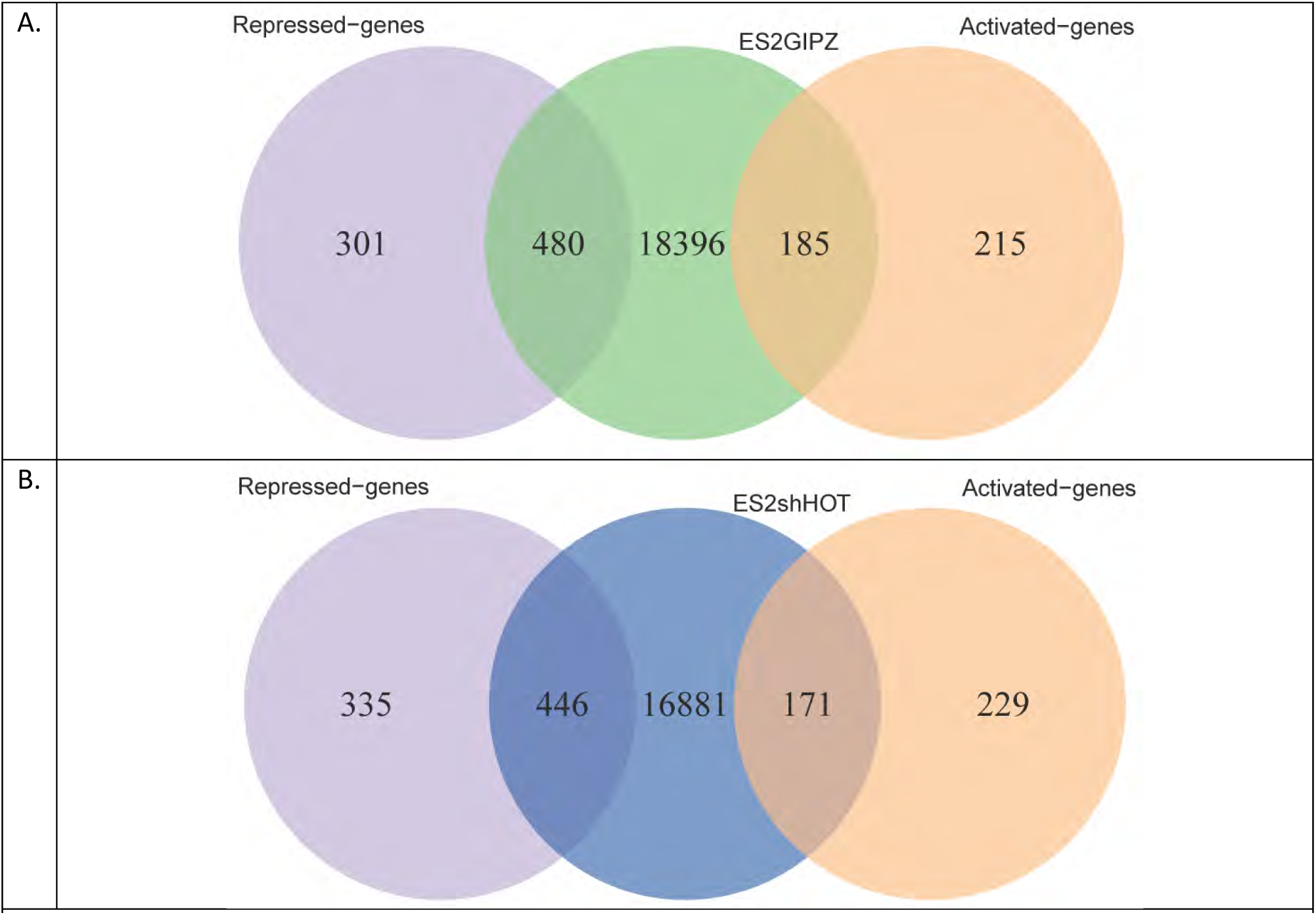
Overlap between genes adjacent to H3K4me1 peaks in (A) ES2GIPZ and (B) ES2shHOT with the HOTAIR-regulated genes. venn_ES2shHOT_H3K4M1_diff_genes.pdf and venn_ES2GIPZ_H3K4M1_diff_genes.pdf

Globally, HOTAIR expression in ES2 cell line induces increased H3K4me1 around TSS of HOTAIR-regulated genes with the characteristics dip at the TSS indicative of nucleosome-free regions (ENCODE project consortium 2012).

**Figure 12.**
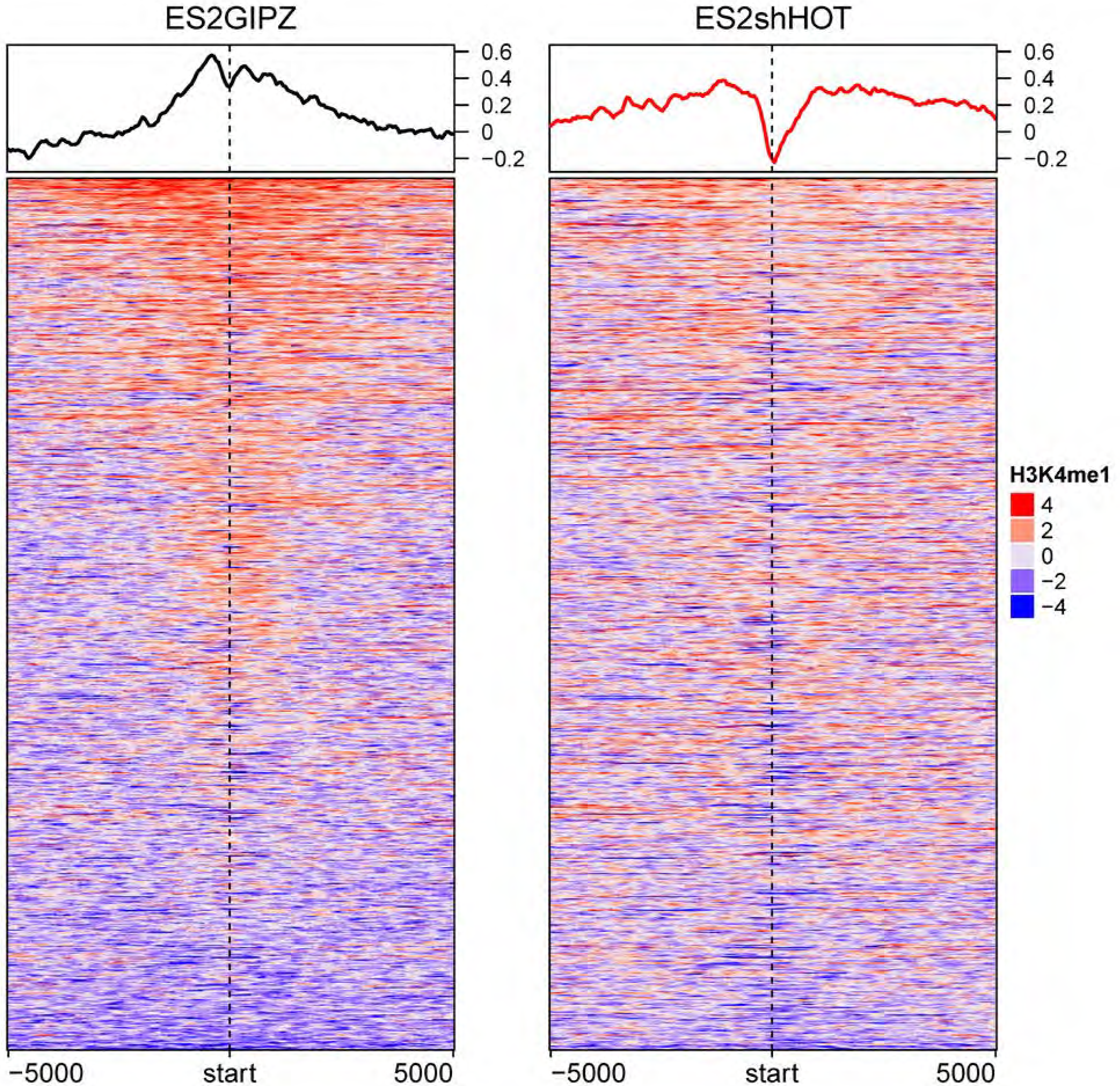
HOTAIR expression induces H3K4me1 binding around TSS of HOTAIR-regulated genes. Visualization of H3K4me1 profiles in ES2GIPZ (left) and ES2shHOTAIR (right) around ± 5000-bp of HOTAIR-regulated genes in ES2. Color represents log_2_ ratio of normalized read counts over input. Mean signals in each cell line are plotted on top. Each row represents HOTAIR-regulated gene. hm_ES2shHOT_ES2GIPZ_HBK4M1_norm_diff.pdf

H3K4me1 binding around TSS of non HOTAIR-regulated genes are shown in hm_ES2shHOT_ES2GIPZ_H3K4M1_norm_nodiff.pdf

### H3K4me1-induced and -repressed binding by HOTAIR expression

We found 66,387 H3K4me1 bindings that are induced by HOTAIR and 48, 859 H3K4me1 bindings that are repressed by HOTAIR. HOTAIR-induced H3K4me1 binding are adjacent to 544 HOTAIR-regulated genes, while HOTAIR-repressed H3K4me1 binding are adjacent to 318 HOTAIR-regulated genes. H3K4me1 bindings adjacent to 295 HOTAIR-regulated genes are exclusively induced by HOTAIR. On the other hand, 69 genes have H3K4me1 bindings that are exclusively repressed by HOTAIR.

#### Methods

66,387 regions with significant increase of H3K4me1 binding (HOTAIR-induced binding) in ES2GIPZ compared to ES2shHOTAIR were identified with FDR ≤ 0.05 using MACS2. 48, 859 regions with significant increase of H3K4me1 binding (HOTAIR-repressed) were identified in ES2shHOTAIR compared to ES2GIPZ.

**Figure 13.**
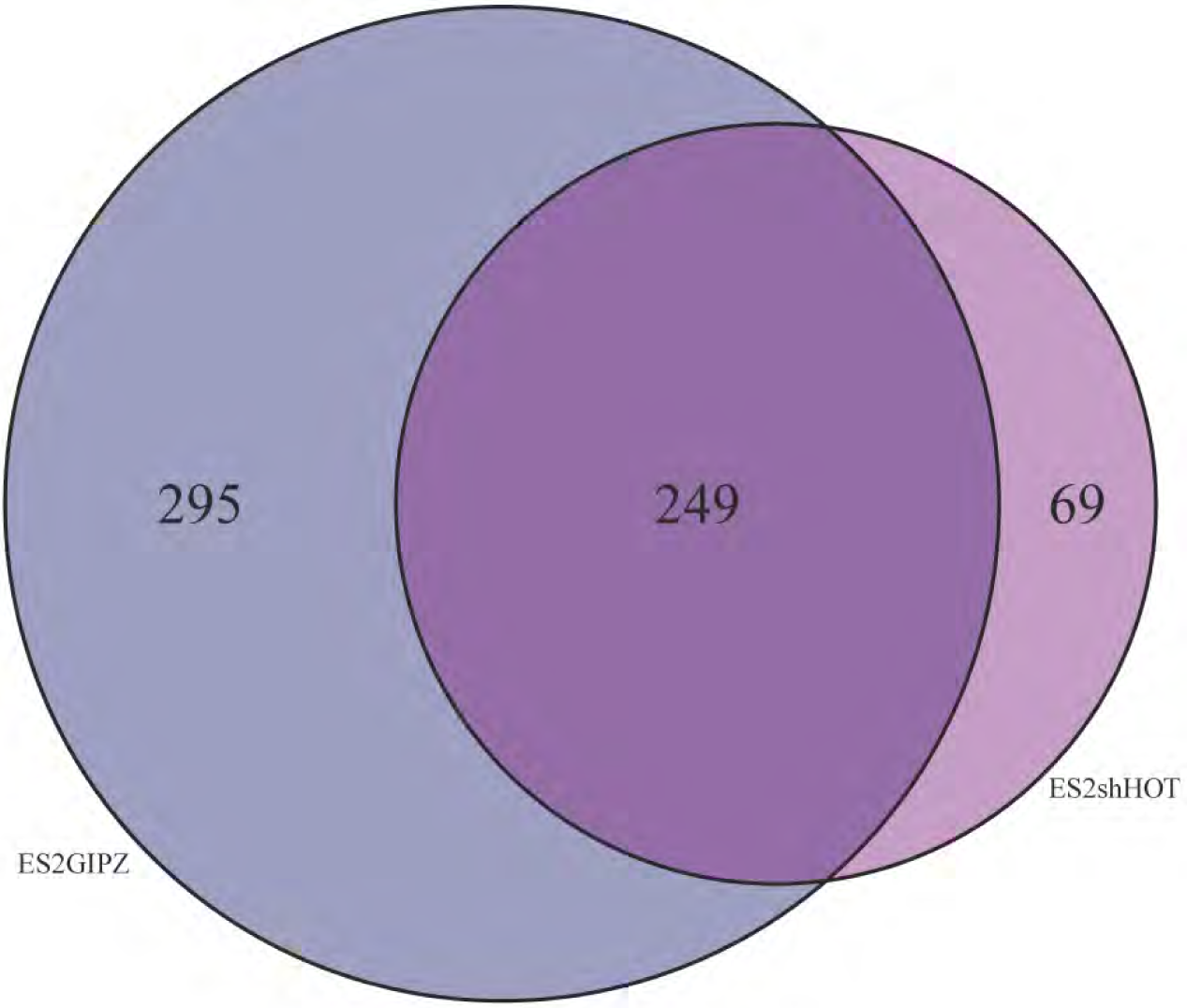
Overlap between HOTAIR-induced and -repressed H3K4me1 bindings. Venn diagram shows overlap between the number of HOTAIR-regulated genes adjacent to increased H3K4me1 binding in ES2GIPZ (dark blue) and increased H3K4me1 binding in ES2shHOTAIR (magenta). Note that multiple differential binding may be adjacent to the same gene. venn_ES2GIPZvsES2shHOT_H3K4M1.pdf.

**Figure 14.**
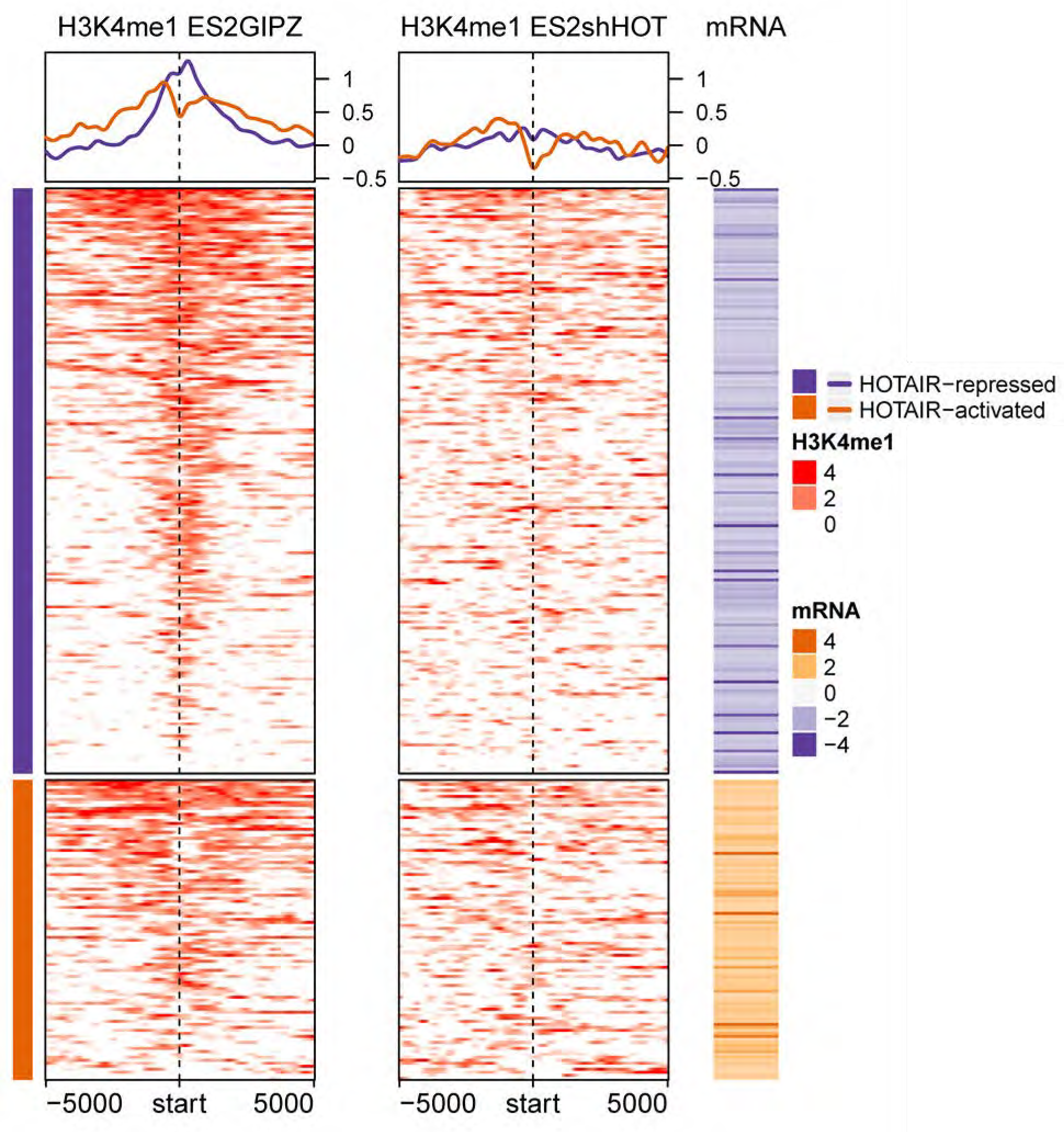
Profiles of HOTAIR-induced H3K4me1 modification around TSS of HOTAIR-regulated genes (295 genes in venn diagram on Figure 13). Each row corresponds to HOTAIR-regulated genes (195 down- and 100 up-regulated genes). Color for H3K4me1 profile represents log_2_ ratio of ChIP to input control. Positive (red) represents enrichment of H3K4me1. Mean H3K4me1 profile in each cell line are plotted on top. The corresponding mRNA expression (log2 ratio) is shown on the right panel. Negative (purple) represents decreased expression in ES2Control compared to ES2shHOTAIR (i.e. genes that are repressed by HOTAIR). “hm_clustES2GIPZenrich_H3K4M1_vsES2shHOT.pdf”

**Figure 14b.**
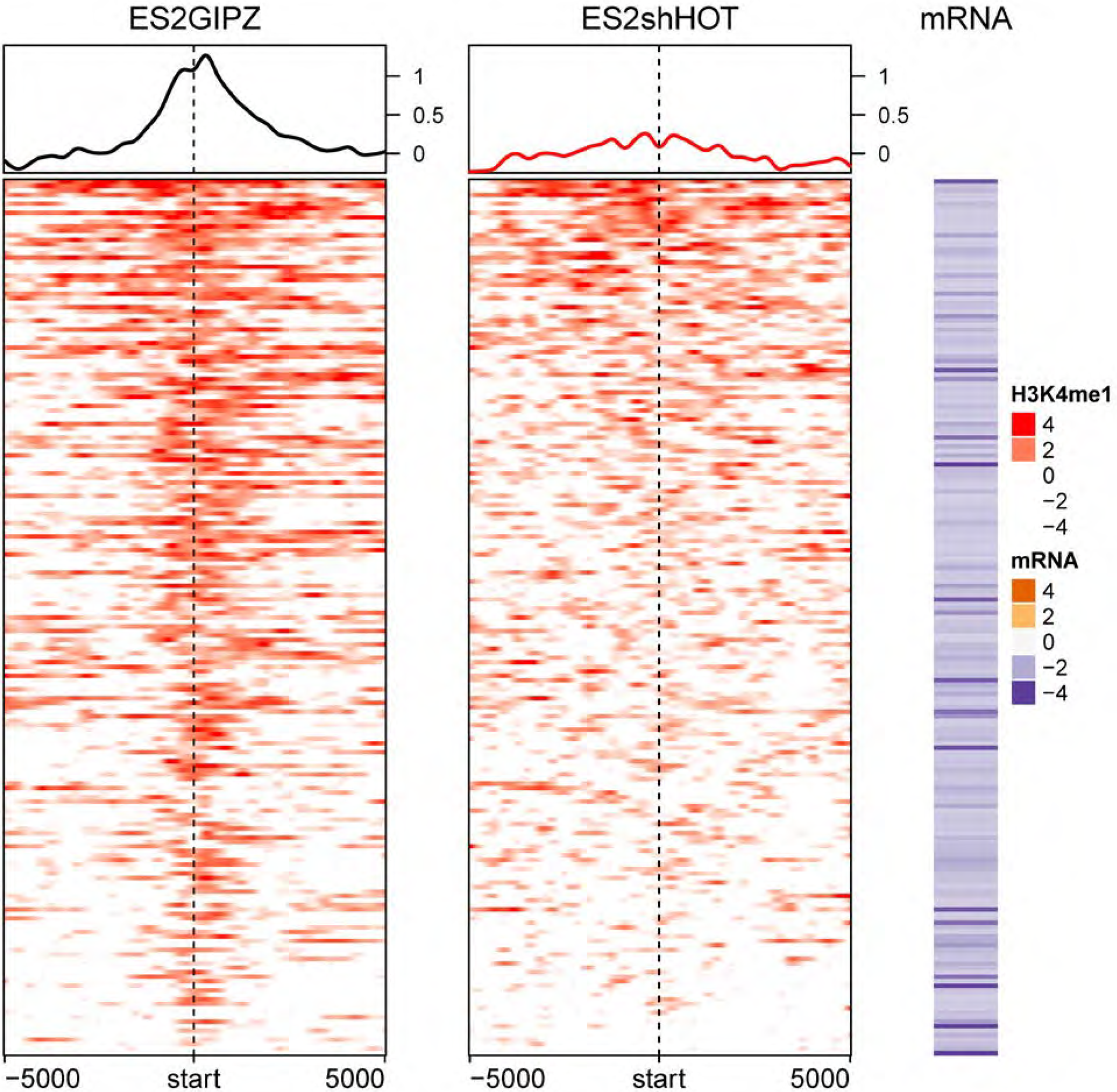
Profiles of HOTAIR-induced H3K4me1 modification around TSS of HOTAIR-repressed genes. Each row corresponds to HOTAIR-repressed genes (195 genes). Color for H3K4me1 profile represents log_2_ ratio of ChIP to input control. Positive (red) represents enrichment of H3K4me1. Mean H3K4me1 profile in each cell line are plotted on top. The corresponding mRNA expression (log2 ratio) is shown on the right panel. Negative (purple) represents decreased expression in ES2Control compared to ES2shHOTAIR (i.e. genes that are repressed by HOTAIR). “hm_clustES2GIPZenrich_H3K4M1_vsES2shHOT_rep_genes.pdf”

**Figure 15.**
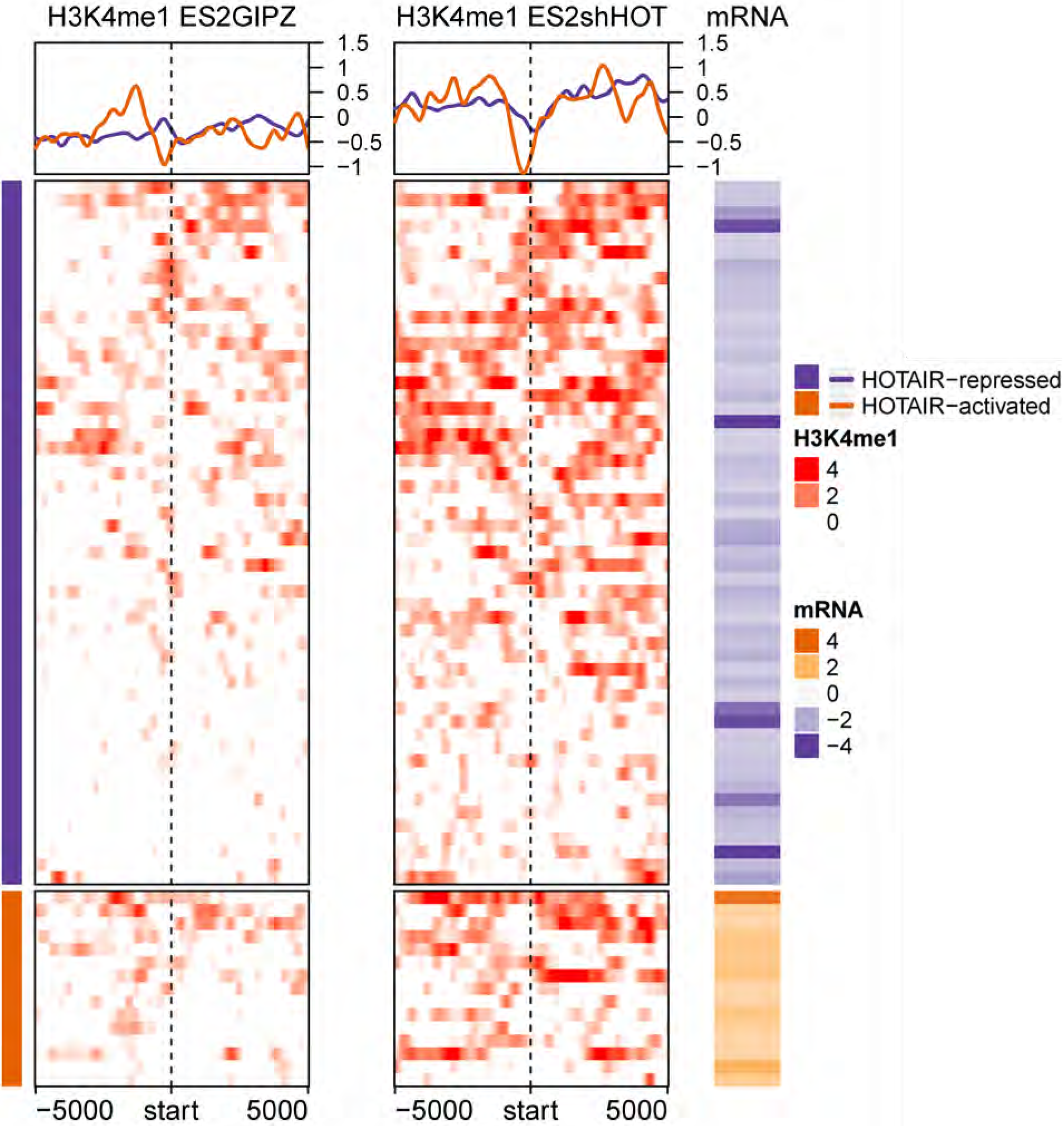
Profiles of HOTAIR-repressed H3K4me1 bindings in ES2GIPZ and ES2shHOTAIR around TSS of HOTAIR-regulated genes (69 genes in venn diagram on Figure 13). Each row corresponds to HOTAIR-regulated genes (54 down- and 15 up-regulated genes). Color for H3K4me1 profile represents log_2_ ratio of ChIP to input control. Positive (red) represents enrichment of H3K4me1. Mean H3K4me1 profile in each cell line are plotted on top. The corresponding mRNA expression (log2 ratio) is shown on the right panel. Negative (purple) represents decreased expression in ES2Control compared to ES2shHOTAIR (i.e. genes that are repressed by HOTAIR). “hm_clustES2shHOTenrich_H3K4M1_vsES2GIPZ.pdf”

## H3K4me2 modulation in ES2 cell line

28,843 (adjacent to 13,337 unique genes) and 109,565 (adjacent to 19,193 unique genes) H3K4me2 peaks were identified at FDR cut-off of 0.05 using MACS2 in ES2GIPZ and ES2shHOTAIR, respectively. Peaks in each cell line were normalized to their corresponding input. 416 and 657 HOTAIR-regulated genes are found adjacent to H3K4me2 peaks in ES2GIPZ and ES2shHOTAIR, respectively.

**Figure 16.**
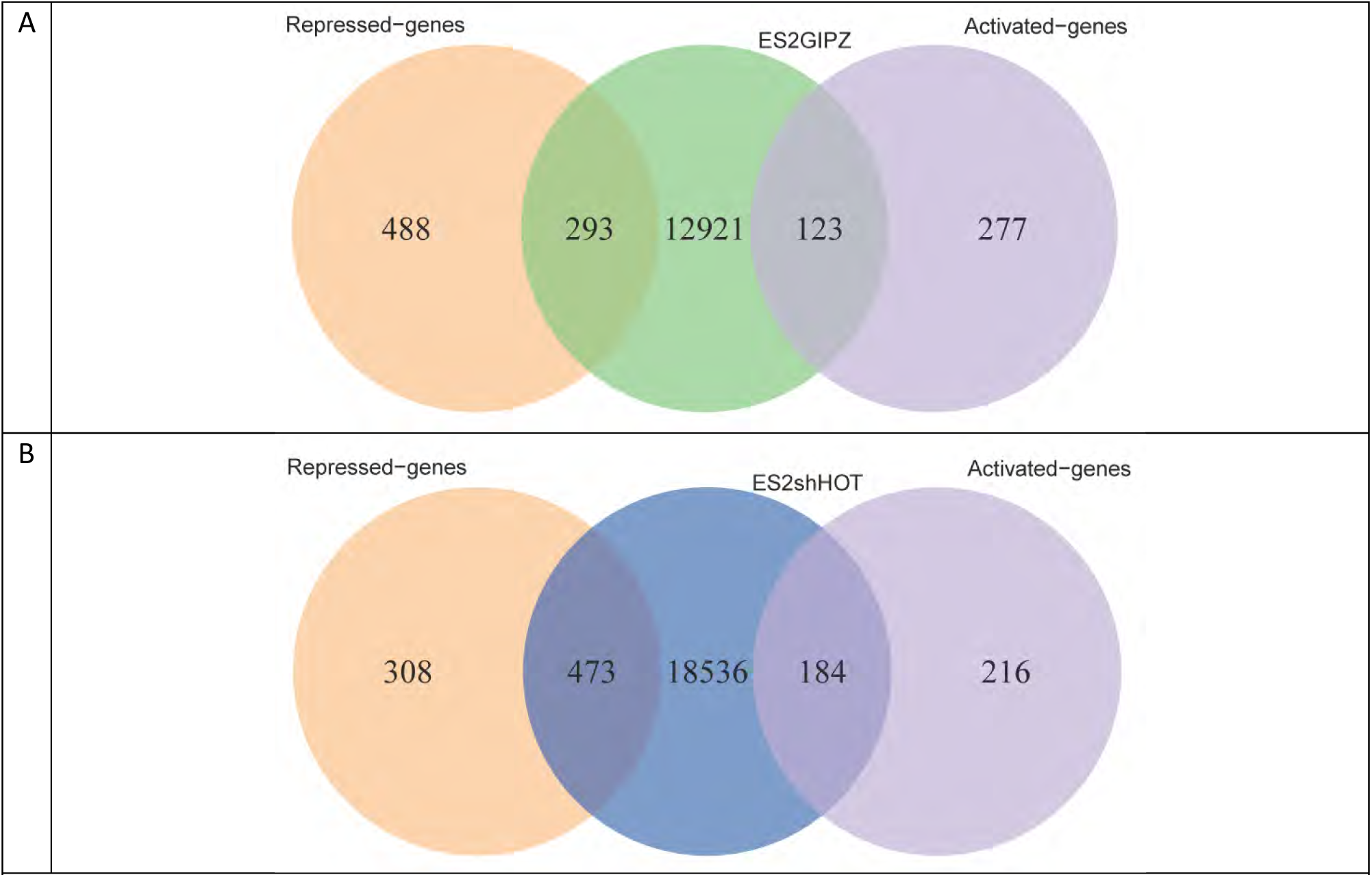
Overlap between genes adjacent to H3K4me1 peaks in (A) ES2GIPZ and (B) ES2shHOT with the HOTAIR-regulated genes.

In contrast to H3K4me1 mark, globally HOTAIR expression represses H3K4me2 modification around TSS of HOTAIR-regulated genes in ES2 cell line.

**Figure 17.**
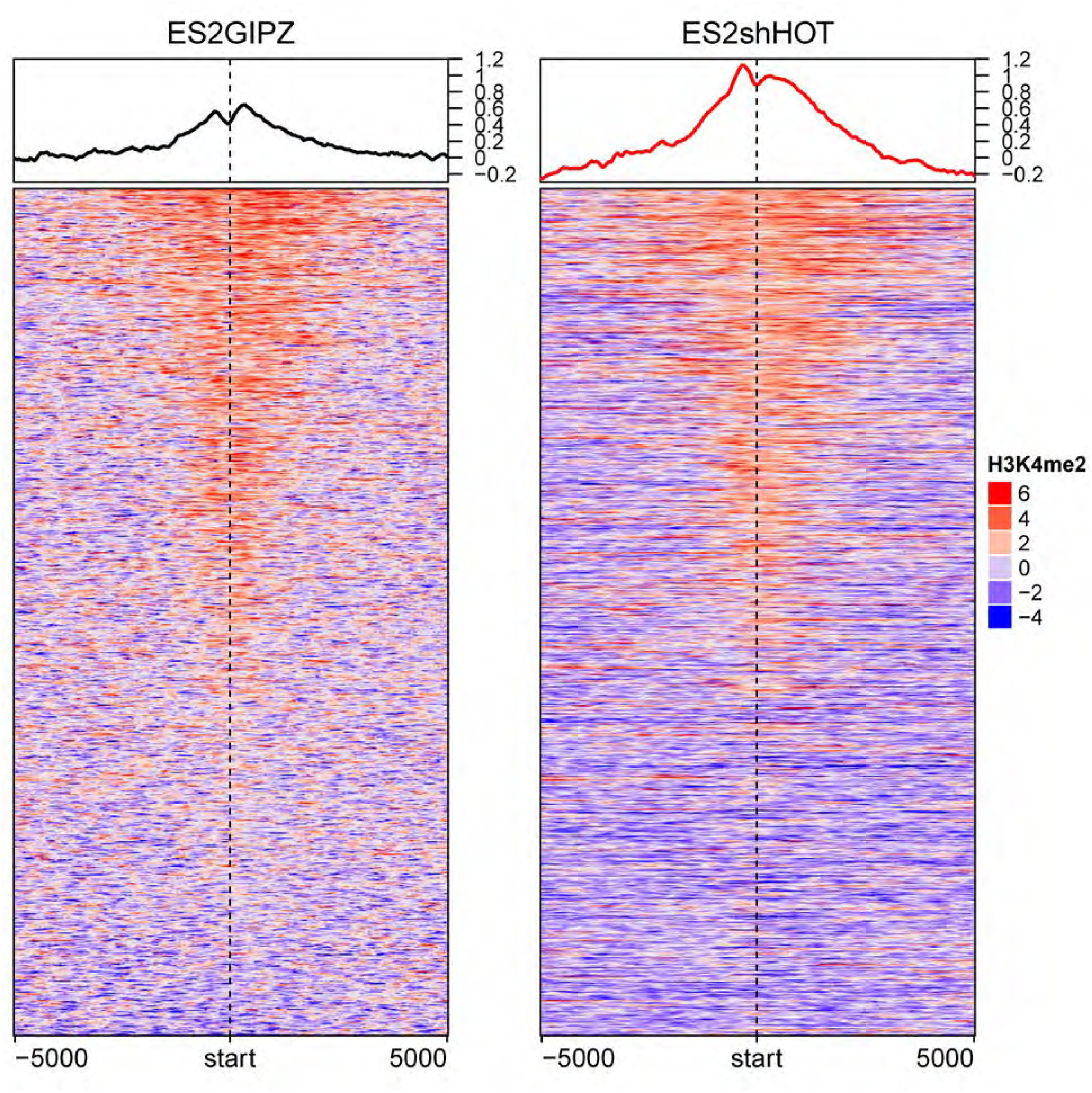
HOTAIR expression induces decreased H3K4me2 binding around TSS of HOTAIR-regulated genes. Visualization of H3K4me2 profiles in ES2GIPZ (left) and ES2shHOTAIR (right) around ± 5000-bp of HOTAIR-regulated genes in ES2. Color represents log_2_ ratio of normalized read counts over input. Mean signals in each cell line are plotted on top. Each row represents HOTAIR-regulated gene. hm_ES2shHOT_ES2GIPZ_H3K4M2_norm_diff.pdf

### H3K4me2- induced and -repressed binding by HOTAIR expression

We found 185 H3K4me2 bindings that are induced by HOTAIR and 92,516 H3K4me2 bindings that are repressed by HOTAIR. HOTAIR-induced H3K4me2 binding are adjacent only to 9 HOTAIR-regulated genes, while HOTAIR-repressed H3K4me2 binding are adjacent to 620 HOTAIR-regulated genes.

H3K4me2 bindings adjacent to only 1 HOTAIR-regulated gene are exclusively induced by HOTAIR. On the other hand, 612 genes have H3K4me2 bindings that are exclusively repressed by HOTAIR.

#### Methods

185 regions with significant increase of H3K4me2 binding (HOTAIR-induced binding) in ES2GIPZ compared to ES2shHOTAIR were identified with FDR ≤ 0.05 using MACS2. 92,516 regions with significant increase of H3K4me2 binding (HOTAIR-repressed) were identified in ES2shHOTAIR compared to ES2GIPZ.

**Figure 18.**
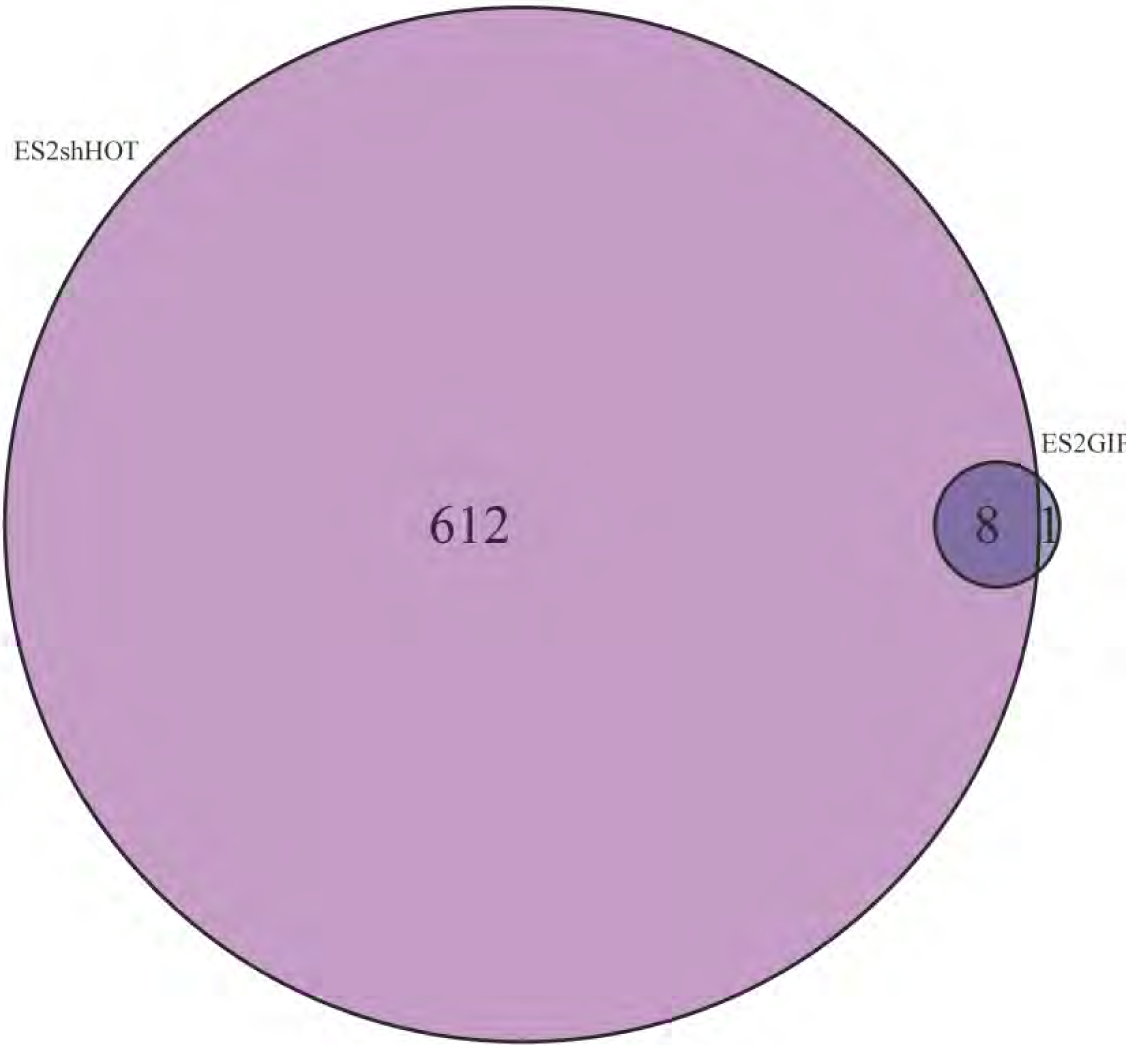
Overlap between HOTAIR-induced and -repressed H3K4me2 bindings. Venn diagram shows overlap between number of HOTAIR-regulated genes adjacent to increased H3K4me2 binding in ES2GIPZ (dark blue) and increased H3K4me2 binding in ES2shHOTAIR (magenta). Note that multiple differential binding may be adjacent to the same gene. venn_ES2GIPZvsES2shHOT_H3K4M2.pdf.

We found decreased H3K4me2 binding induced by HOTAIR in ES2 cell line around TSS of HOTAIR-regulated genes. Interestingly, although H3K4me2 bindings around some of the non-HOTAIR-regulated genes are also increased without HOTAIR expression, these increased bindings were not at the TSSs. “hm_ES2shHOT_ES2GIPZ_H3K4M2_norm_nodiff.pdf”

**Figure 19.**
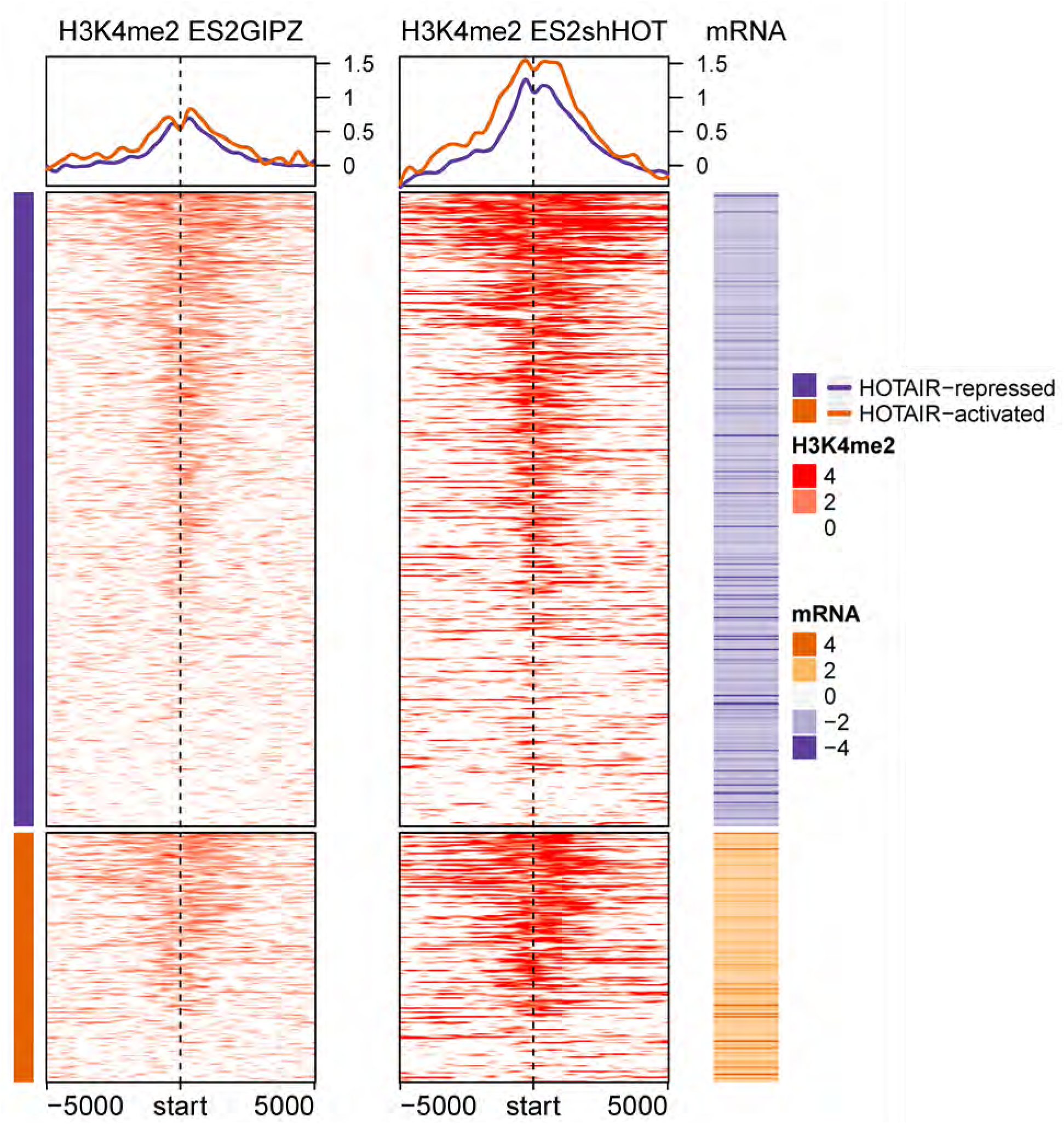
Profiles of HOTAIR-repressed H3K4me2 bindings in ES2GIPZ and ES2shHOTAIR around TSS of HOTAIR-regulated genes (612 genes in venn diagram on Figure 13). Each row corresponds to HOTAIR-regulated genes (439 down- and 172 up-regulated genes). Color for H3K4me2 profile represents log_2_ ratio of ChIP to input control. Positive (red) represents enrichment of H3K4me2. Mean H3K4me2 profile in each cell line are plotted on top. The corresponding mRNA expression (log2 ratio) is shown on the right panel. Negative (purple) represents decreased expression in ES2Control compared to ES2shHOTAIR (i.e. genes that are repressed by HOTAIR). “hm_clustES2shHOTenrich_H3K4M2_vsES2GIPZ.pdf”

**Figure 19b.**
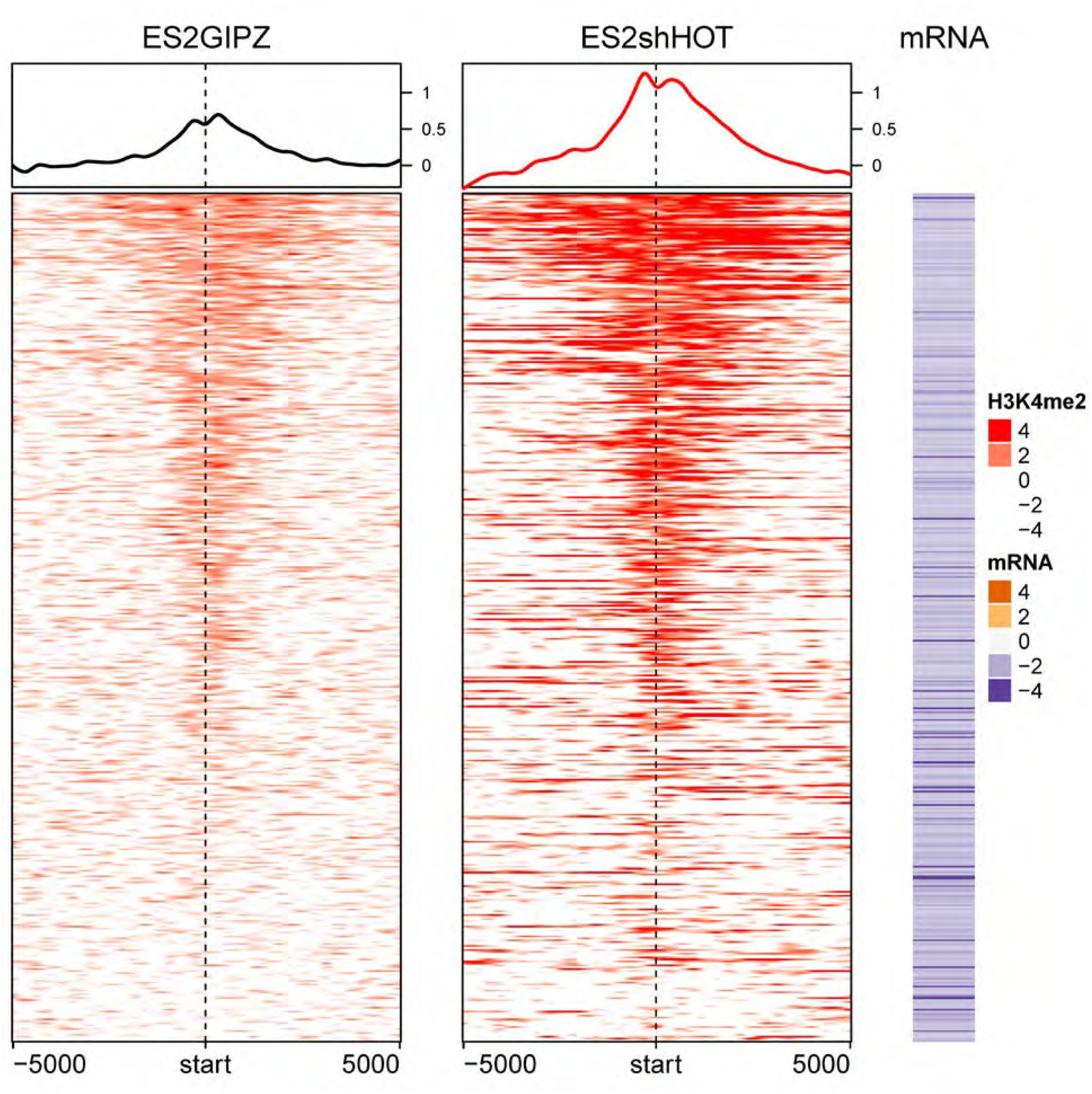
Profiles of HOTAIR-repressed H3K4me2 bindings in ES2GIPZ and ES2shHOTAIR around TSS of HOTAIR-repressed genes. Each row corresponds to **HOTAIR-repressed genes**. Color for H3K4me2 profile represents log_2_ ratio of ChIP to input control. Positive (red) represents enrichment of H3K4me2. Mean H3K4me2 profile in each cell line are plotted on top. The corresponding mRNA expression (log2 ratio) is shown on the right panel. Negative (purple) represents decreased expression in ES2Control compared to ES2shHOTAIR (i.e. genes that are repressed by HOTAIR). “hm_clustES2shHOTenrich_H3K4M2_vsES2GIPZ_rep_genes.pdf”

## hMSC cell line

### H3K4me1 enrichment in hMSC in relation to HOTAIR expression

48,017 (adjacent to 15,314 unique genes) and 1,885 (adjacent to 696 unique genes) H3K4me1 broad peaks were identified at FDR ≤ 0.05 using MACS2 in hMSCHOTAIR and hMSCGFP, respectively. Peaks in each cell line were normalized to their corresponding input. 3,657 genes were found to be differentially regulated by HOTAIR (FDR ≤ 0.05 and fold-change ≥ 2) (fold-change = hMSC-hTERTHOTAIR/hMSC-hTERTGFP, i.e. genes regulated by HOTAIR have positive fold-change). 15,314 and 696 HOTAIR-regulated genes are found adjacent to H3K4me1 peaks in hMSCHOTAIR and hMSCGFP, respectively.

**Figure 20.**
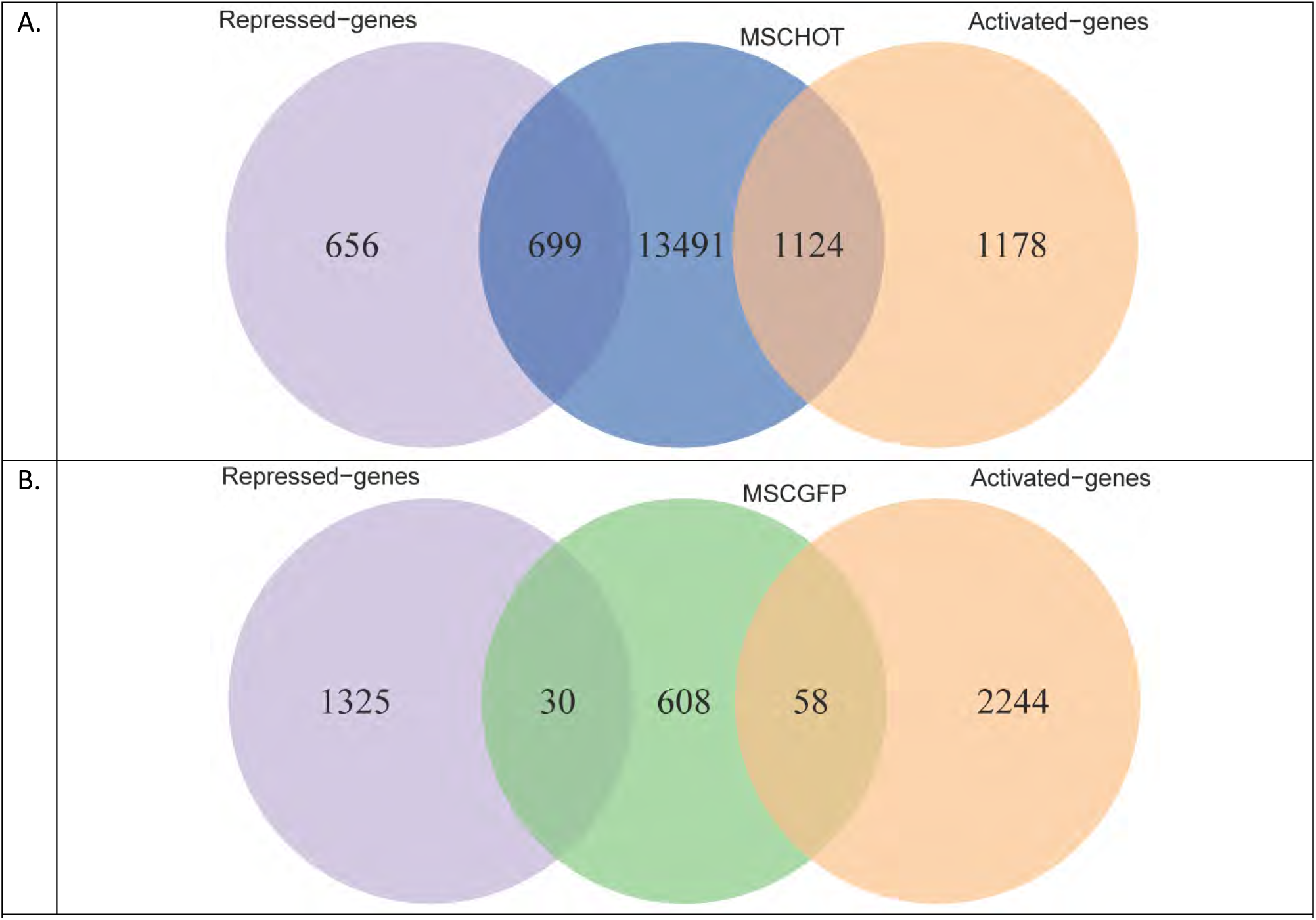
Overlap between genes adjacent to H3K4me1 peaks in (A) hMSCHOTAIR and (B) hMSCGFP with the HOTAIR-regulated genes. venn_MSCHOT_H3K4M1_diff_genes.pdf and venn_MSCGFP_H3K4M1_diff_genes.pdf

Globally, HOTAIR expression in hMSC cell line induces increased H3K4me1 around TSS of HOTAIR-regulated genes.

**Figure 21.**
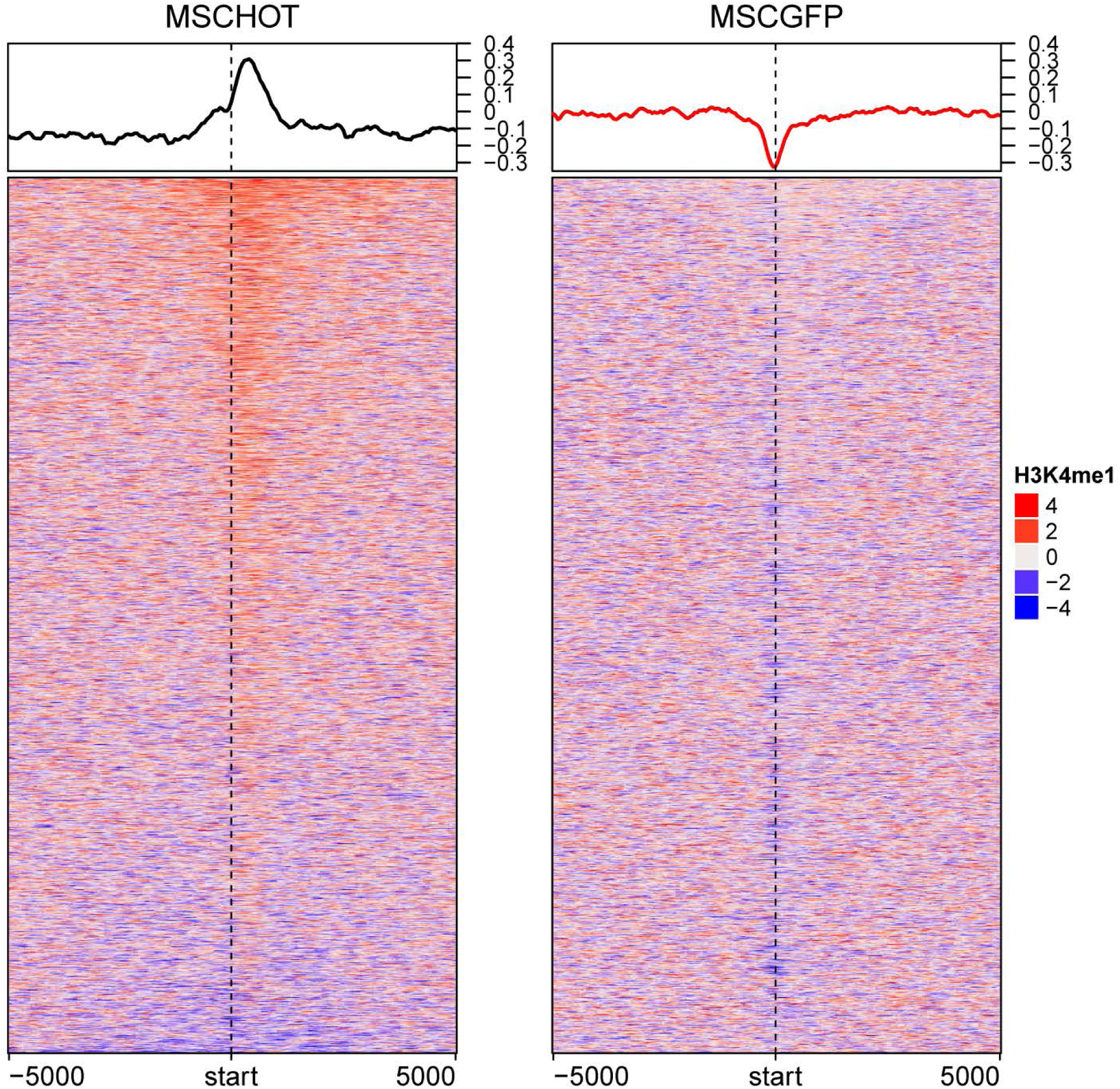
HOTAIR expression induces H3K4me1 binding around TSS of HOTAIR-regulated genes. Visualization of H3K4me1 profiles in hMSCHOTAIR (left) and hMSCGFP (right) around ± 5000-bp of HOTAIR-regulated genes. Color represents log_2_ ratio of normalized read counts over input. Mean signals in each cell line are plotted on top. Each row represents HOTAIR-regulated gene. hm_MSCGFP_MSCHOT_H3K4M1_norm_diff.pdf.

H3K4me1 binding around TSS of non HOTAIR-regulated genes are shown in hm_MSCGFP_MSCHOT_H3K4M1_norm_nodiff.pdf

### H3K4me1- induced and -repressed binding by HOTAIR expression

We found 12,089 H3K4me1 bindings that are induced by HOTAIR and 1,394 H3K4me1 bindings that are repressed by HOTAIR. HOTAIR-induced H3K4me1 binding are adjacent to 667 HOTAIR-regulated genes, while HOTAIR-repressed H3K4me1 binding are adjacent to 7 HOTAIR-regulated genes. H3K4me1 bindings adjacent to 662 HOTAIR-regulated genes are exclusively induced by HOTAIR. On the other hand, 7 genes have H3K4me1 bindings that are exclusively repressed by HOTAIR.

#### Methods

12,089 regions with significant increase of H3K4me1 binding (HOTAIR-induced binding) in hMSCHOTAIR compared to hMSCGFP were identified with FDR ≤ 0.05 using MACS2. 1,394 regions with significant increase of H3K4me1 binding (HOTAIR-repressed) were identified in hMSCGFP compared to hMSCHOTAIR.

**Figure 22.**
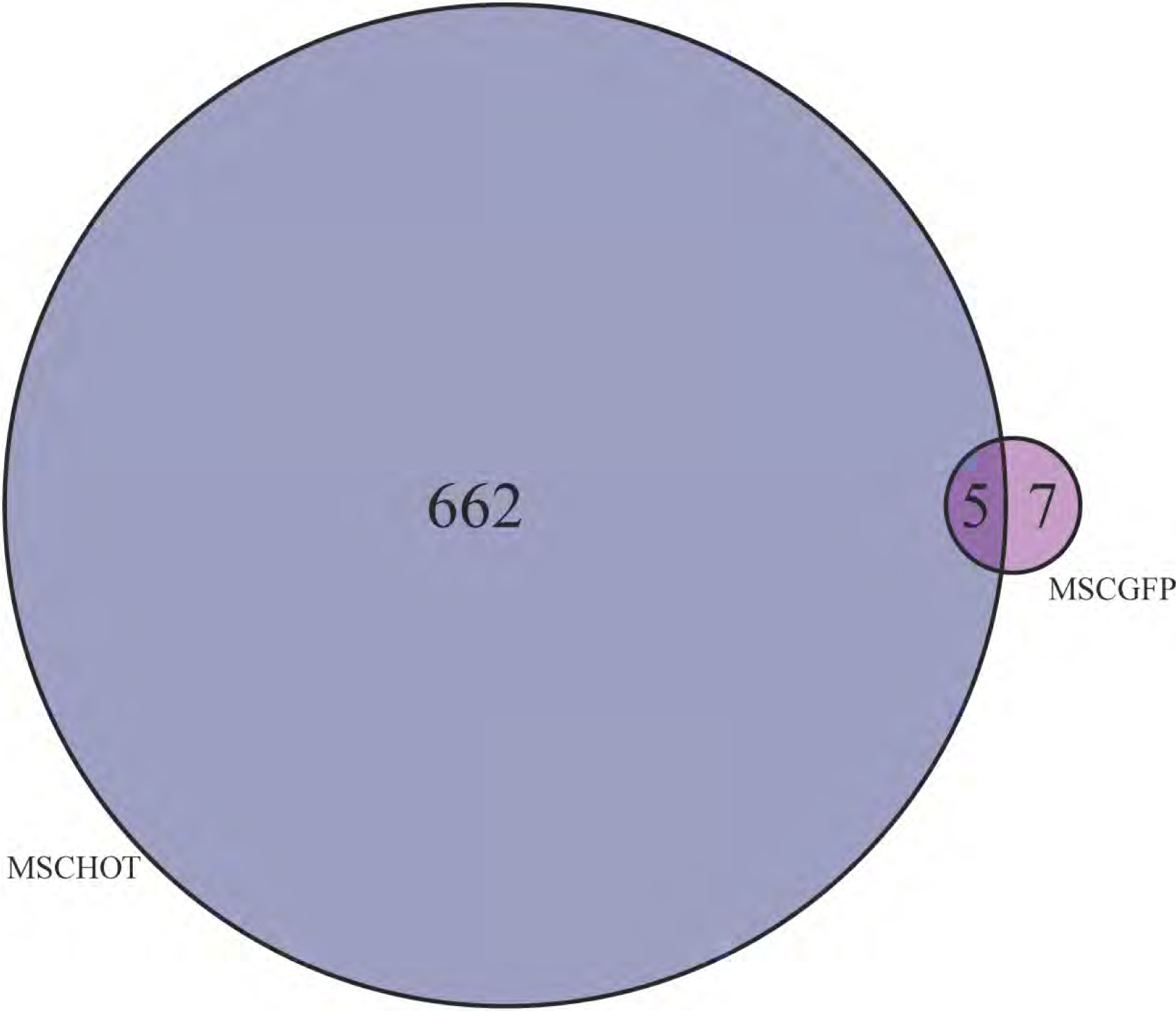
Overlap between HOTAIR-induced and –repressed H3K4me1 bindings. Venn diagram shows overlap between the number of HOTAIR-regulated genes adjacent to increased

H3K4me1 binding in hMSCHOTAIR (dark blue) and increased H3K4me1 binding in hMSCGFP (magenta). Note that multiple differential binding may be adjacent to the same gene. venn_MSCHOTvsMSCGFP_H3K4M1.pdf

**Figure 23.**
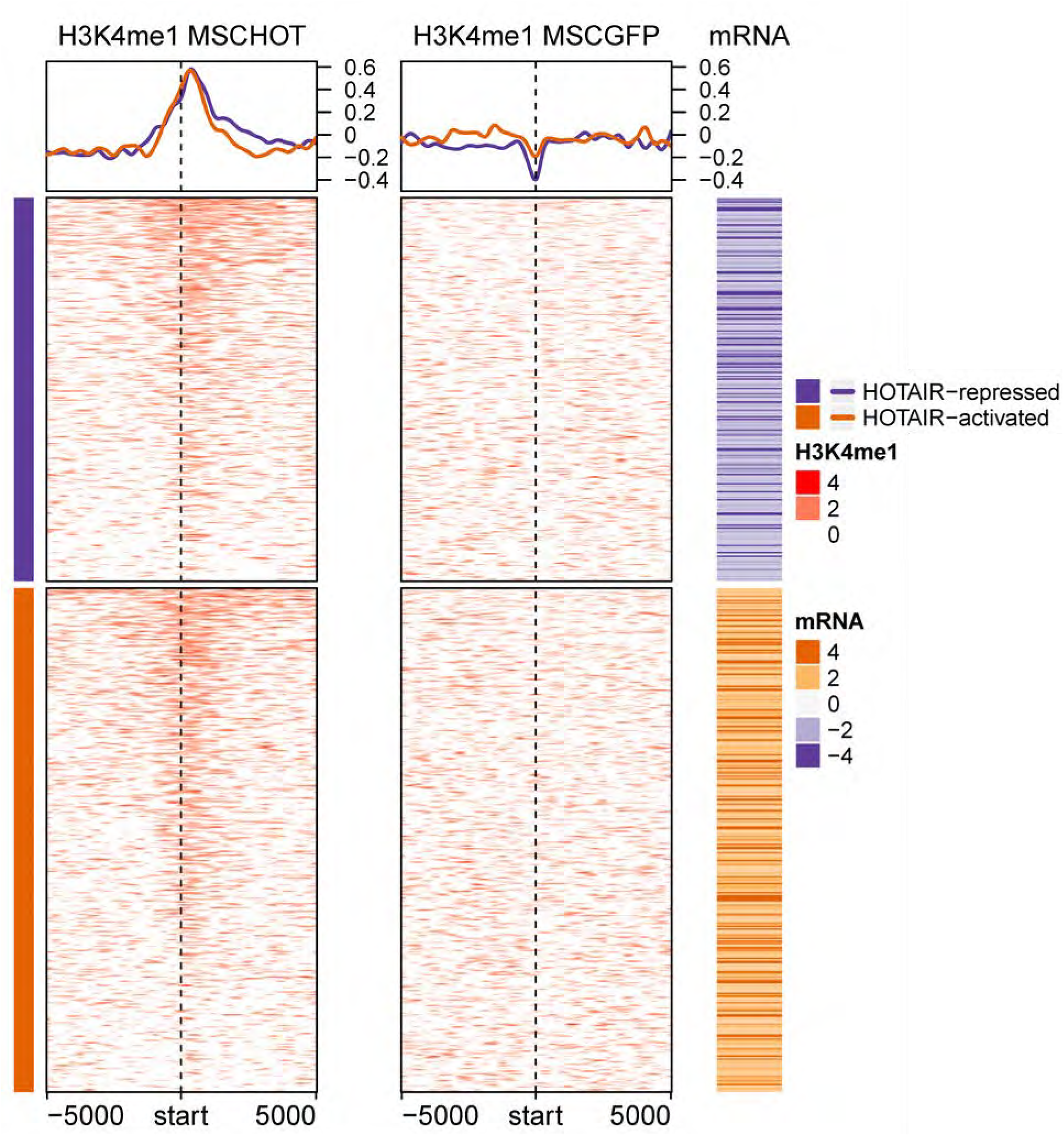
Profiles of HOTAIR-induced H3K4me1 modification around TSS of HOTAIR-regulated genes (662 genes in venn diagram on Figure 22). Each row corresponds to HOTAIR-regulated genes (286 down- and 376 up-regulated genes). Color for H3K4me1 profile represents log_2_ ratio of ChIP to input control. Positive (red) represents enrichment of H3K4me1. Mean H3K4me1 profile in each cell line are plotted on top. The corresponding mRNA expression (log2 ratio) is shown on the right panel. Negative (purple) represents decreased expression in hMSC-hTERTHOTAIR compared to hMSC-hTERTGFP (i.e. genes that are repressed by HOTAIR). “hm_clustMSCHOTenrich_H3K4M1_vsMSCGFP.pdf”

**Figure 23b.**
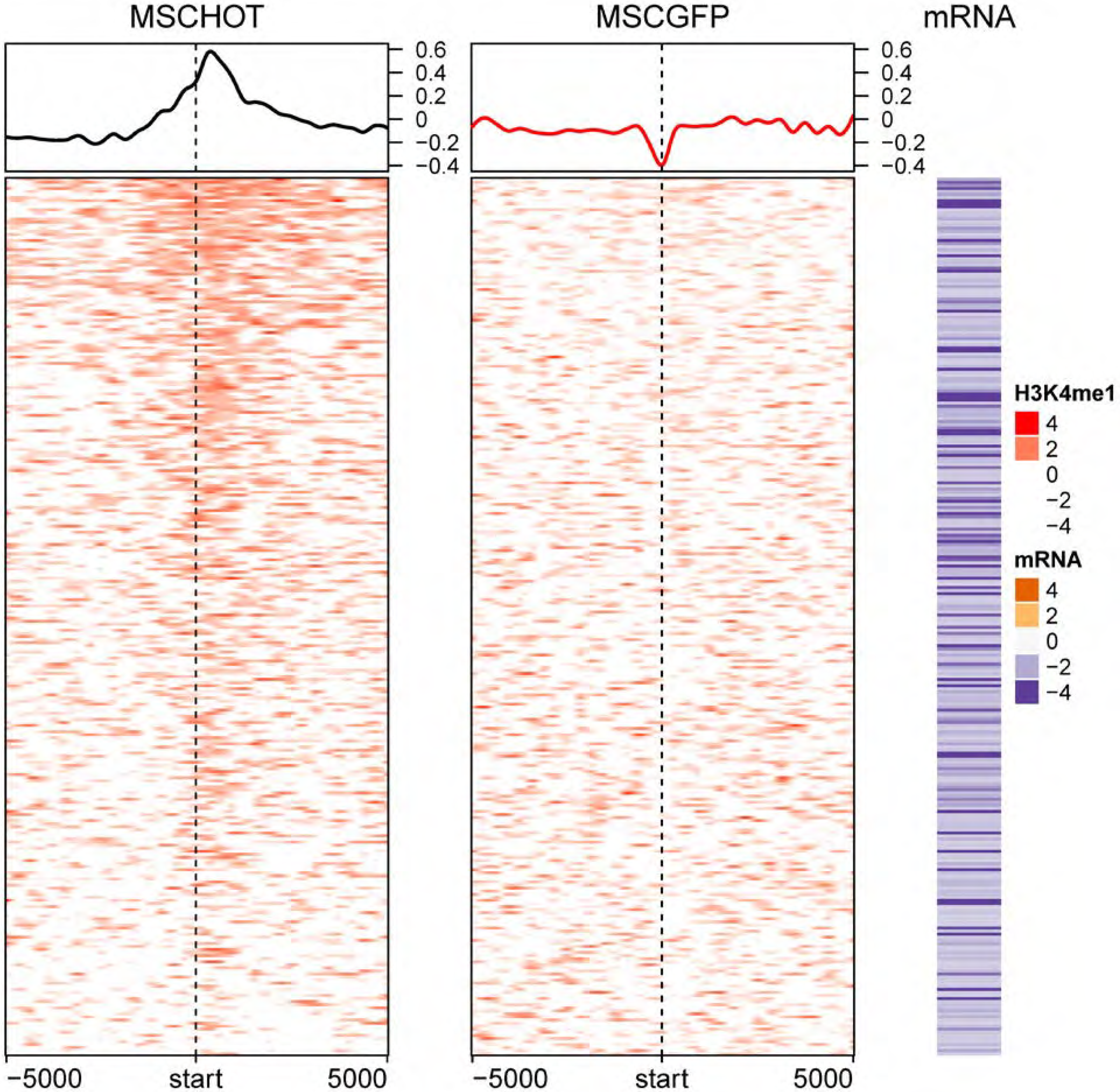
Profiles of HOTAIR-induced H3K4me1 modification around TSS of HOTAIR-repressed genes. Each row corresponds to HOTAIR-regulated genes (286 downregulated genes). Color for H3K4me1 profile represents log_2_ ratio of ChIP to input control. Positive (red) represents enrichment of H3K4me1. Mean H3K4me1 profile in each cell line are plotted on top. The corresponding mRNA expression (log2 ratio) is shown on the right panel. Negative (purple) represents decreased expression in hMSC-hTERTHOTAIR compared to hMSC-hTERTGFP (i.e. genes that are repressed by HOTAIR). “hm_clustMSCHOTenrich_H3K4M1_vsMSCGFP_rep_genes.pdf”

**Figure 24.**
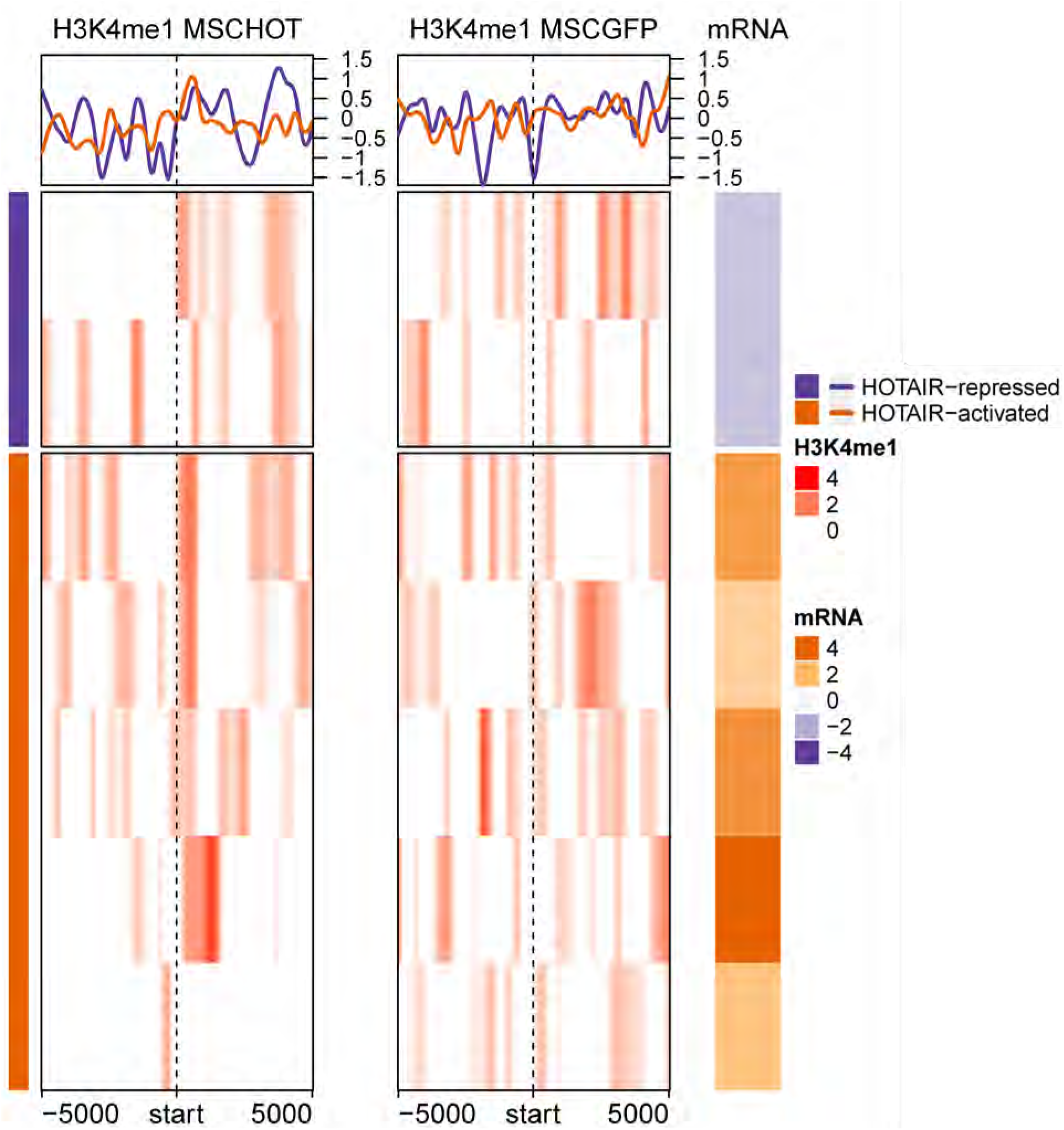
Profiles of HOTAIR-repressed H3K4me1 binding around TSS of HOTAIR-regulated genes (7 genes in venn diagram on Figure 22). Each row corresponds to HOTAIR-regulated genes (2 down- and 5 up-regulated genes). Color for H3K4me1 profile represents log_2_ ratio of ChIP to input control. Positive (red) represents enrichment of H3K4me1. Mean H3K4me1 profile in each cell line are plotted on top. The corresponding mRNA expression (log2 ratio) is shown on the right panel. Negative (purple) represents decreased expression in hMSC-hTERTHOTAIR compared to hMSC-hTERTGFP (i.e. genes that are repressed by HOTAIR). “hm_clustMSCGFPenrich_H3K4M1_vsMSCHOT.pdf”

### H3K4me2 modulation in hMSC cell lines

37,543 (adjacent to 15,427 unique genes) and 35,678 (adjacent to 14,251 unique genes) H3K4me2 peaks were identified at FDR cut-off of 0.05 using MACS2 in hMSCHOTAIR and hMSCGFP, respectively. Peaks in each cell line were normalized to their corresponding input. 1,823 and 1,709 HOTAIR-regulated genes are found adjacent to H3K4me2 peaks in hMSCHOTAIR and hMSCGFP, respectively.

**Figure 25.**
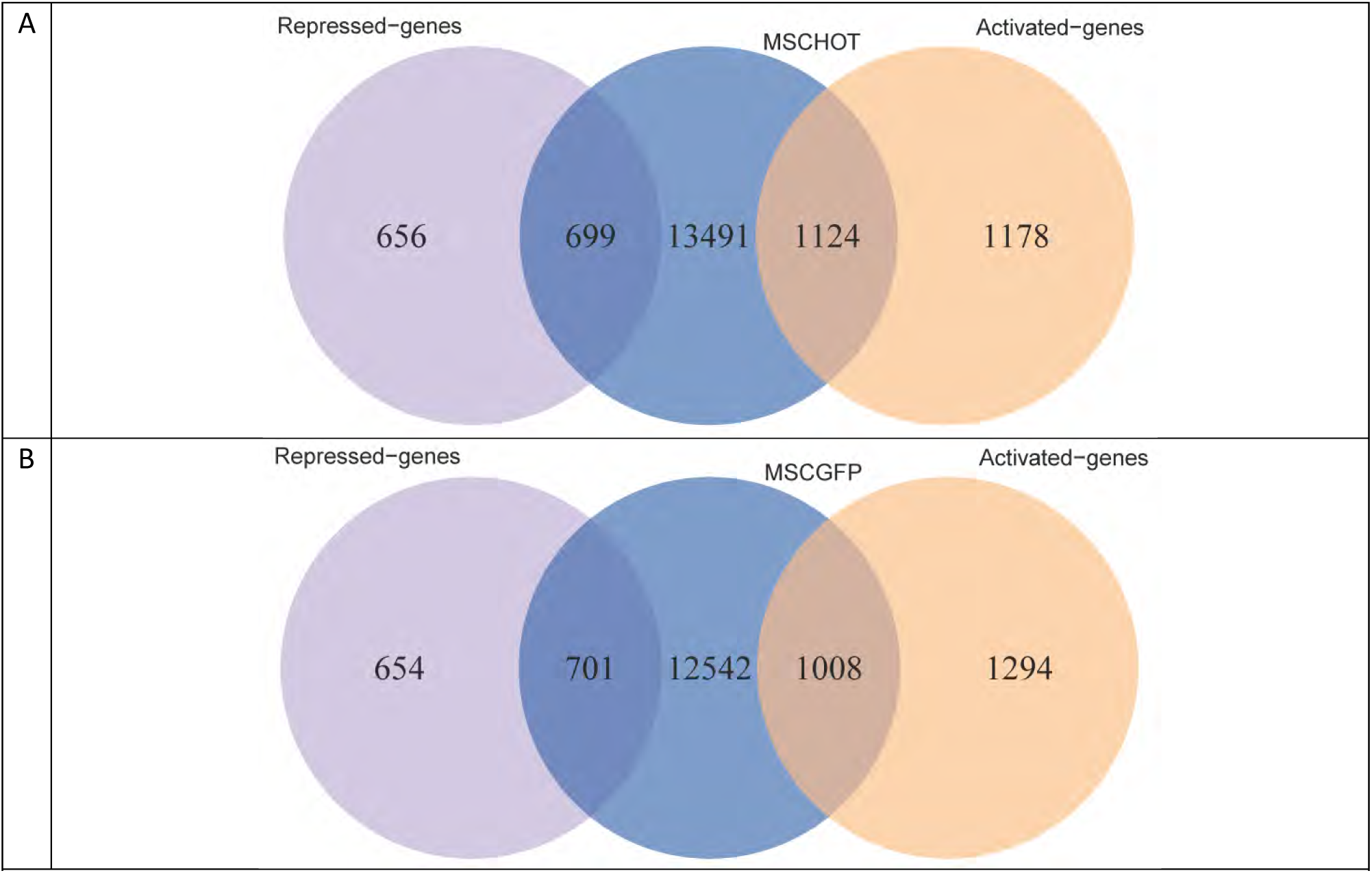
Overlap between genes adjacent to H3K4me2 peaks in (A) MSCHOTAIR and (B) MSCGFP with the HOTAIR-regulated genes.

While the H3K4me2 bindings around HOTAIR-regulated genes in ES2 cell line are globally repressed by HOTAIR, in hMSC cell line the repression by HOTAIR is very minimal (Figure 26).

**Figure 26.**
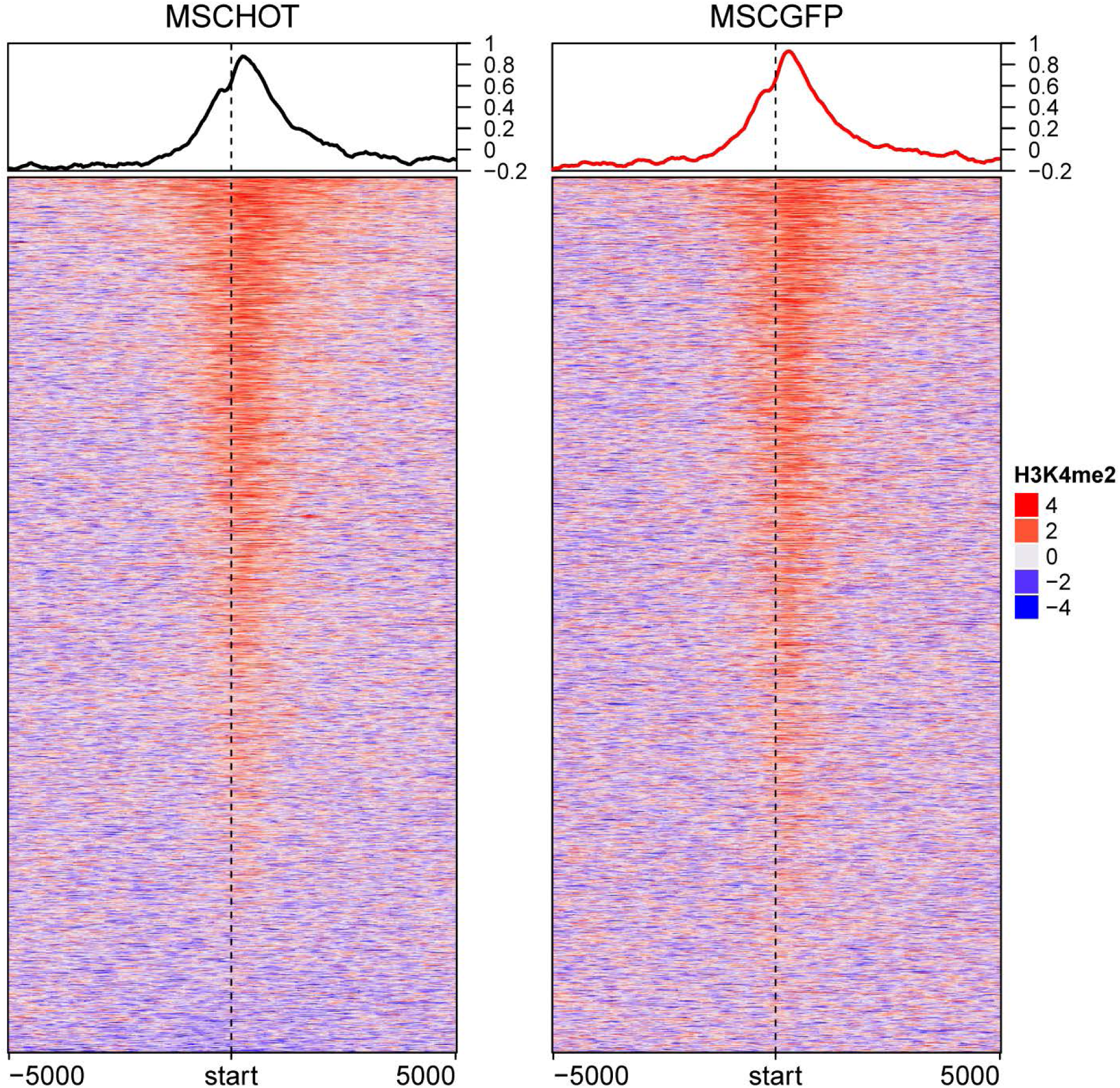
HOTAIR expression induces a slight decrease of H3K4me2 binding around TSS of HOTAIR-regulated genes. Visualization of H3K4me2 profiles in hMSCHOTAIR (left) and hMSCGFP (right) around ± 5000-bp of HOTAIR-regulated genes. Color represents log_2_ ratio of normalized read counts over input. Mean signals in each cell line are plotted on top. Each row represents HOTAIR-regulated gene. hm_MSCGFP_MSCHOT_H3K4M2_norm_diff.pdf.

H3K4me2 bindings around TSS of non-differentially expressed genes is shown in hm_MSCGFP_MSCHOT_H3K4M2_norm_nodiff.pdf

### H3K4me2- induced and -repressed binding by HOTAIR expression

We found 1,876 H3K4me2 bindings that are significantly induced by HOTAIR and 8,417 H3K4me2 bindings that are repressed by HOTAIR. HOTAIR-induced H3K4me2 binding are adjacent to 186 HOTAIR-regulated genes, while HOTAIR-repressed H3K4me2 binding are adjacent to 748 HOTAIR-regulated genes. H3K4me2 bindings adjacent to only 106 HOTAIR-regulated gene are exclusively induced by HOTAIR. On the other hand, 668 genes have H3K4me2 bindings that are exclusively repressed by HOTAIR.

#### Methods

1,876 regions with significant increase of H3K4me2 binding (HOTAIR-induced binding) in hMSCHOTAIR compared to hMSCGFP were identified with FDR ≤ 0.05 using MACS2. 8,417 regions with significant increase of H3K4me2 binding (HOTAIR-repressed) were identified in hMSCGFP compared to hMSCHOTAIR.

**Figure 27.**
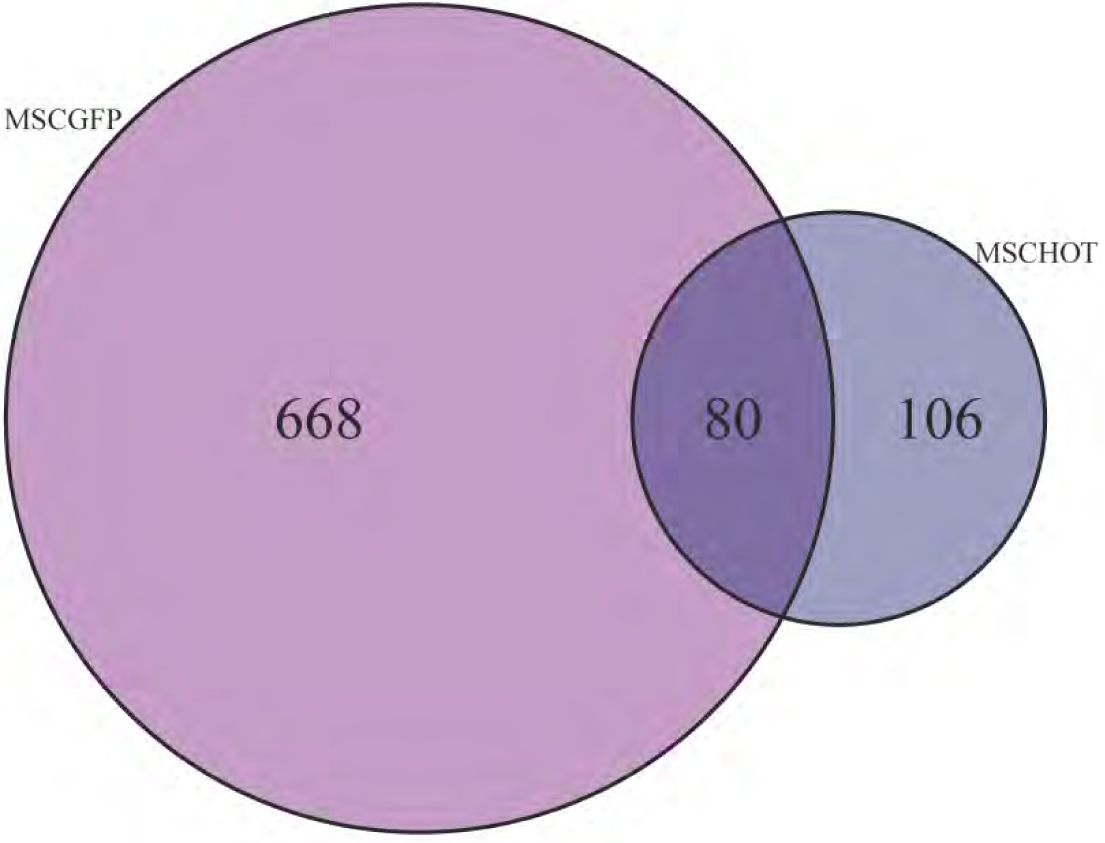
Overlap between HOTAIR-induced and -repressed H3K4me2 bindings. Venn diagram shows overlap between numbers of HOTAIR-regulated genes adjacent to increased H3K4me2 binding in hMSCHOTAIR (dark blue) and increased H3K4me2 binding in hMSCGFP (magenta). Note that multiple differential binding may be adjacent to the same gene. venn_MSCHOTvsMSCGFP_H3K4M2.pdf.

**Figure 28.**
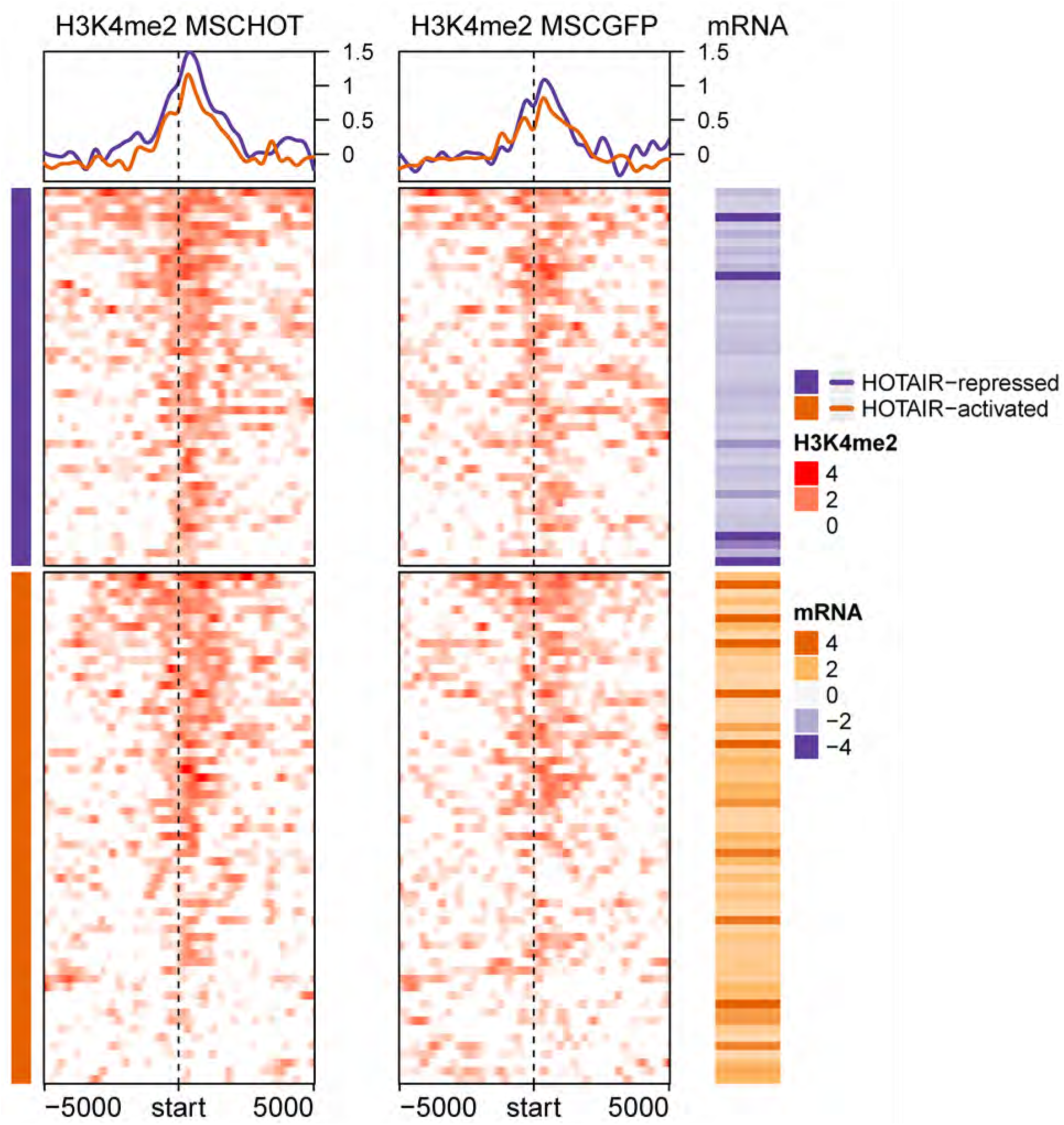
Profiles of HOTAIR-induced H3K4me2 bindings in hMSC cell lines around TSS of HOTAIR-regulated genes (106 genes in venn diagram on Figure 27). Each row corresponds to HOTAIR-regulated genes (45 down- and 61 up-regulated genes). Color for H3K4me2 profile represents log_2_ ratio of ChIP to input control. Positive (red) represents enrichment of H3K4me2. Mean H3K4me2 profile in each cell line are plotted on top. The corresponding mRNA expression (log2 ratio) is shown on the right panel. Negative (purple) represents decreased expression in hMSC-hTERTHOTAIR compared to hMSC- hTERTGFP (i.e. genes that are repressed by HOTAIR). “hm_clustMSCHOTenrich_H3K4M2_vsMSCGFP.pdf”

**Figure 29.**
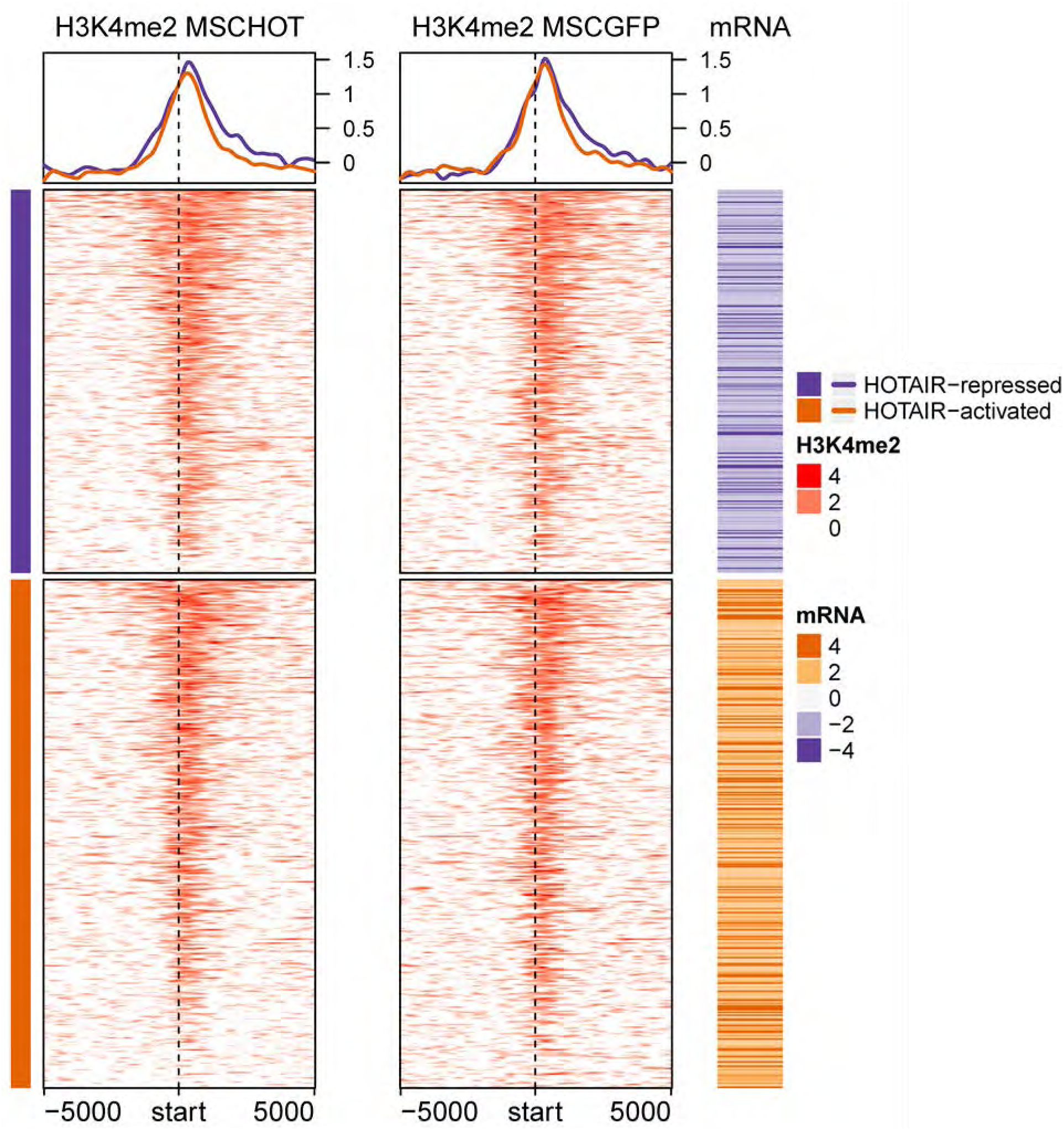
Profiles of HOTAIR-repressed H3K4me2 bindings in hMSC cell lines around TSS of HOTAIR-regulated genes (668 genes in venn diagram on Figure 13). Each row corresponds to HOTAIR-regulated genes (287 down- and 381 up-regulated genes). Color for H3K4me2 profile represents log_2_ ratio of ChIP to input control. Positive (red) represents enrichment of H3K4me2. Mean H3K4me2 profile in each cell line are plotted on top. The corresponding mRNA expression (log2 ratio) is shown on the right panel. Negative (purple) represents decreased expression in hMSC-hTERTHOTAIR compared to hMSC- hTERTGFP (i.e. genes that are repressed by HOTAIR). “hm_clustMSCGFPenrich_H3K4M2_vsMSCHOT.pdf”

**Figure 29b.**
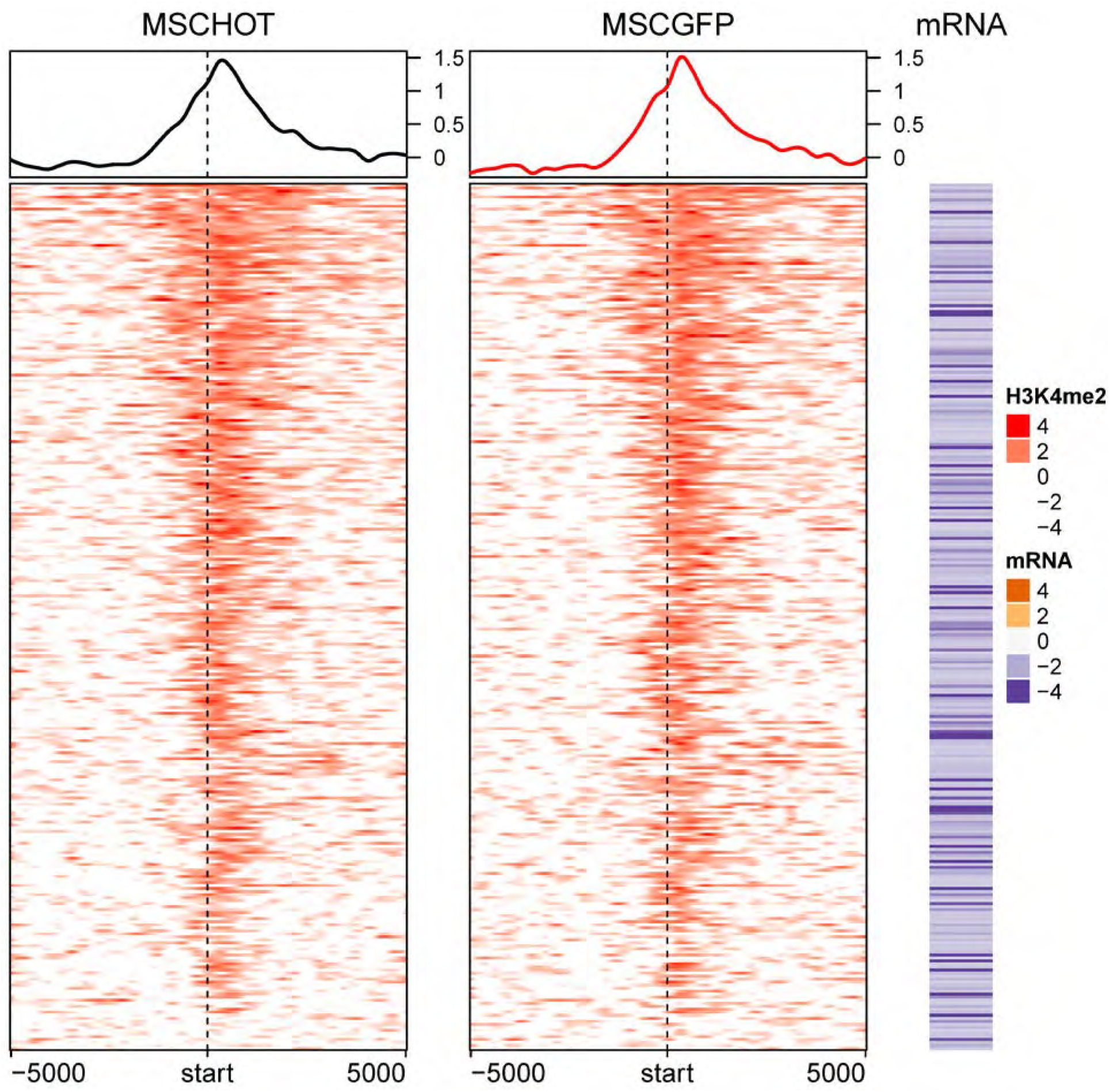
Profiles of HOTAIR-repressed H3K4me2 bindings in hMSC cell lines around TSS of **HOTAIR-repressed genes**. Each row corresponds to HOTAIR-repressed genes (287 downregulated genes). Color for H3K4me2 profile represents log_2_ ratio of ChIP to input control. Positive (red) represents enrichment of H3K4me2. Mean H3K4me2 profile in each cell line are plotted on top. The corresponding mRNA expression (log2 ratio) is shown on the right panel. Negative (purple) represents decreased expression in hMSC-hTERTHOTAIR compared to hMSC-hTERTGFP (i.e. genes that are repressed by HOTAIR). “hm_clustMSCGFPenrich_H3K4M2_vsMSCHOT.pdf”

## SKES Cell Lines

### H3K4me1 enrichment in SKES in relation to HOTAIR expression

67,906 (adjacent to 16,872 genes) and 359 (adjacent to 243 genes) H3K4me1 broad peaks were identified at FDR ≤ 0.05 using MACS2 in SKESGIPZ and SKESshHOTAIR, respectively. Peaks in each cell line were normalized to their corresponding input. 301 genes were found to be differentially regulated by HOTAIR (FDR ≤ 0.05 and absolute fold-change ≥ 2). Genes regulated by HOTAIR have positive fold-change. 194 and 2 HOTAIR-regulated genes are found adjacent to H3K4me1 peaks in SKESGIPZ and SKESshHOTAIR, respectively.

**Figure 30.**
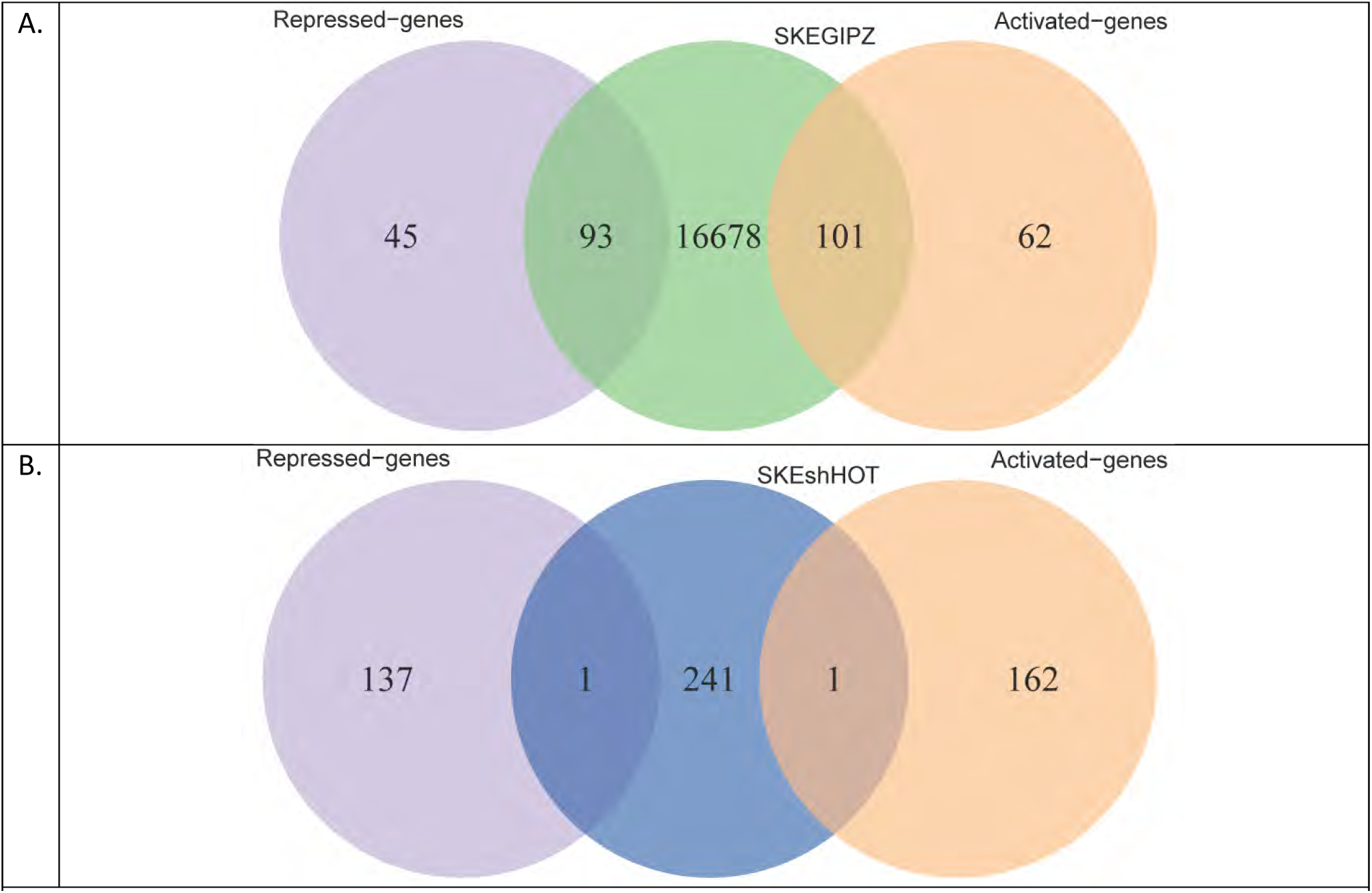
Overlap between genes adjacent to H3K4me1 peaks in (A) SKESGIPZ and (B) SKESshHOT with the HOTAIR-regulated genes. venn_SKEshHOT_H3K4M2_diff_genes.pdf and venn_SKEGIPZ_H3K4M2_diff_genes.pdf

In SKES cell line, HOTAIR expression seems to induce increased H3K4me1 in broad regions surrounding TSS of HOTAIR-regulated genes. Without HOTAIR expression, there is loss of H3K4me1 binding at their TSSs.

**Figure 31.**
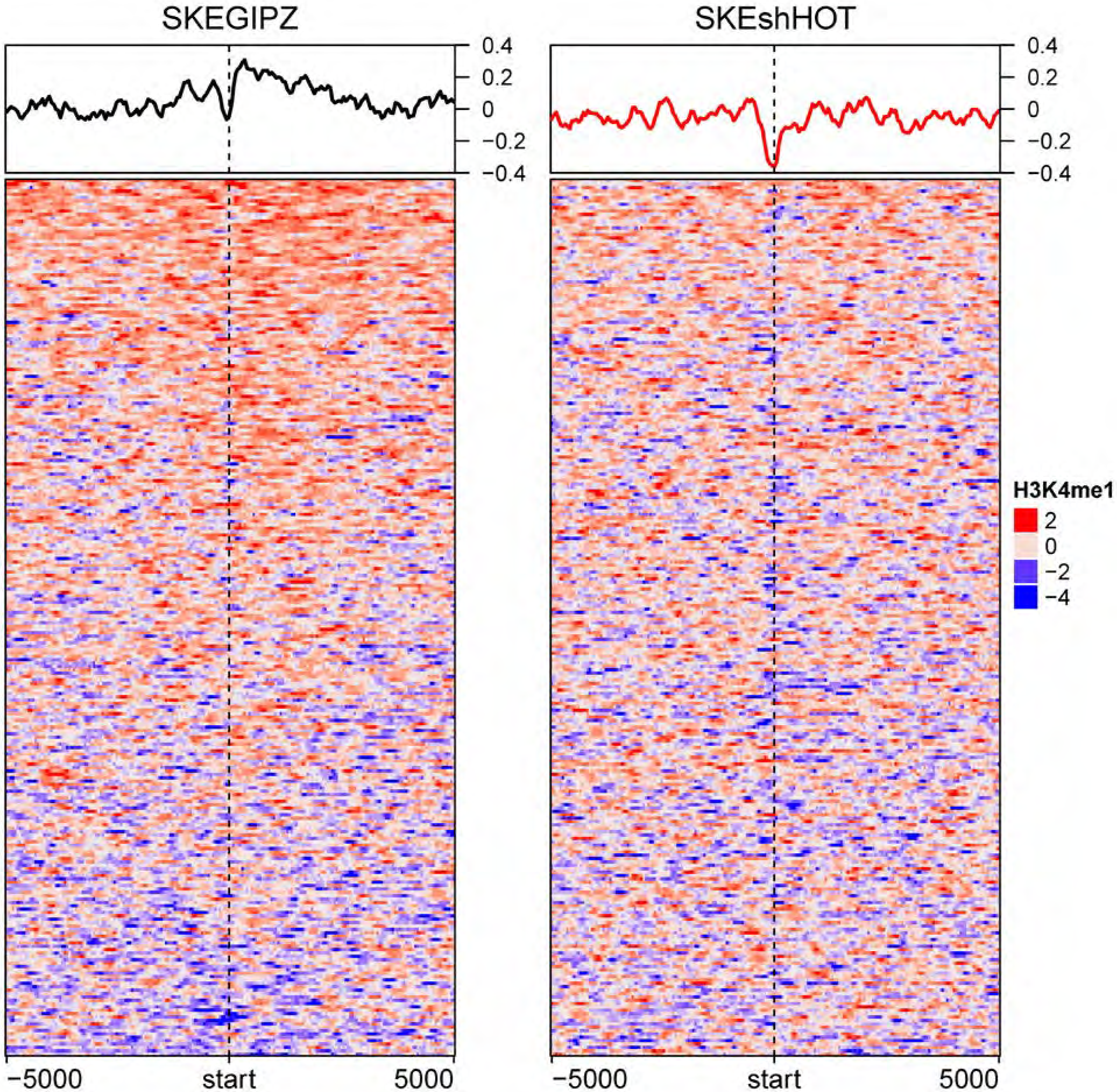
HOTAIR expression induces H3K4me1 binding around TSS of HOTAIR-regulated genes. Visualization of H3K4me1 profiles in SKESGIPZ (left) and SKESshHOTAIR (right) around ± 5000-bp of HOTAIR-regulated genes in ES2. Color represents log_2_ ratio of normalized read counts over input. Mean signals in each cell line are plotted on top. Each row represents HOTAIR-regulated gene. hm_SKEshHOT_SKEGIPZ_H3K4M1_norm_diff.pdf

H3K4me1 binding profile around TSS of non-differentially expressed genes is shown in hm_SKEshHOT_SKEGIPZ_H3K4M1_norm_nodiff.pdf

### H3K4me1- induced and -repressed binding by HOTAIR expression

We found 1,757 H3K4me1 bindings that are induced by HOTAIR and 51 H3K4me1 bindings that are repressed by HOTAIR. HOTAIR-induced H3K4me1 binding are adjacent to 21 HOTAIR-regulated genes, while there are no HOTAIR-regulated genes near HOTAIR-repressed H3K4me1 binding.

#### Methods

1,757 regions with significant increase of H3K4me1 binding (HOTAIR-induced binding) in SKESGIPZ compared to SKESshHOTAIR were identified with FDR ≤ 0.05 using MACS2 diff. 51 regions with significant increase of H3K4me1 binding (HOTAIR-repressed) were identified in SKESshHOTAIR compared to SKESGIPZ.

**Figure 32.**
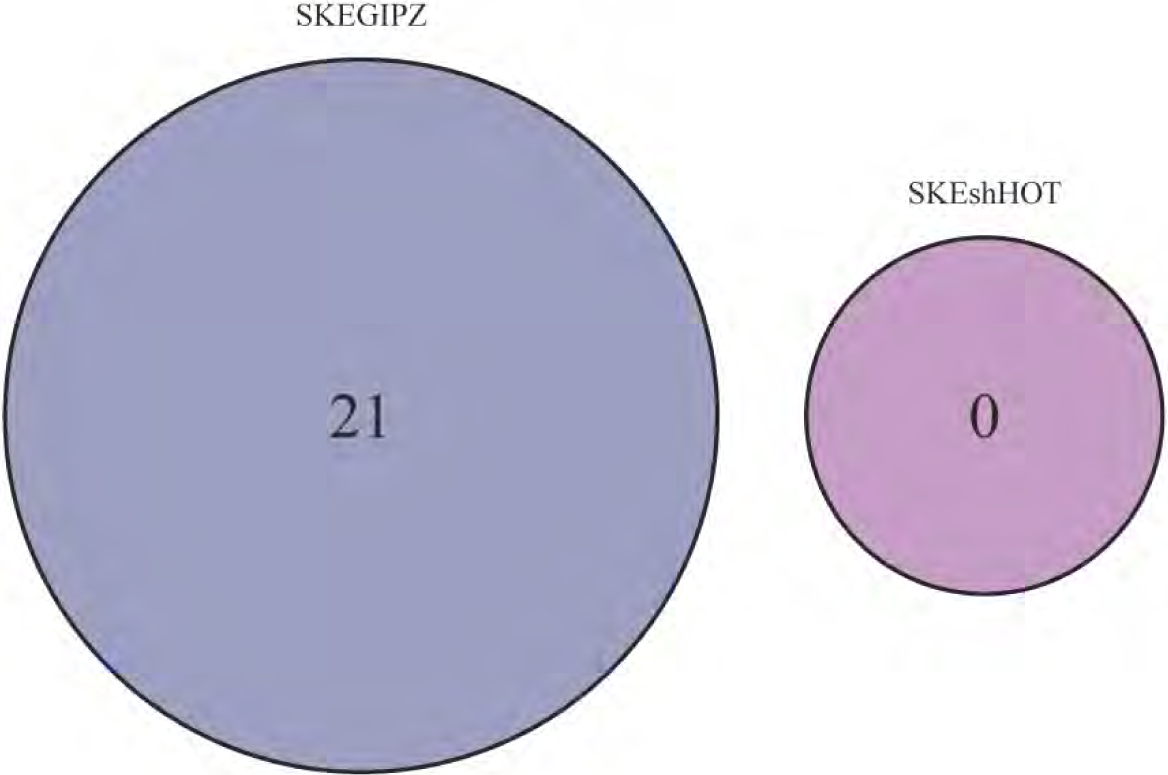
Venn diagram shows the number of HOTAIR-regulated genes adjacent to increased H3K4me1 binding in SKESGIPZ (dark blue) and increased H3K4me1 binding in SKESshHOTAIR (magenta). Note that multiple differential binding may be adjacent to the same gene. venn_SKEGIPZvsSKEshHOT_H3K4M1.pdf.

**Figure 33.**
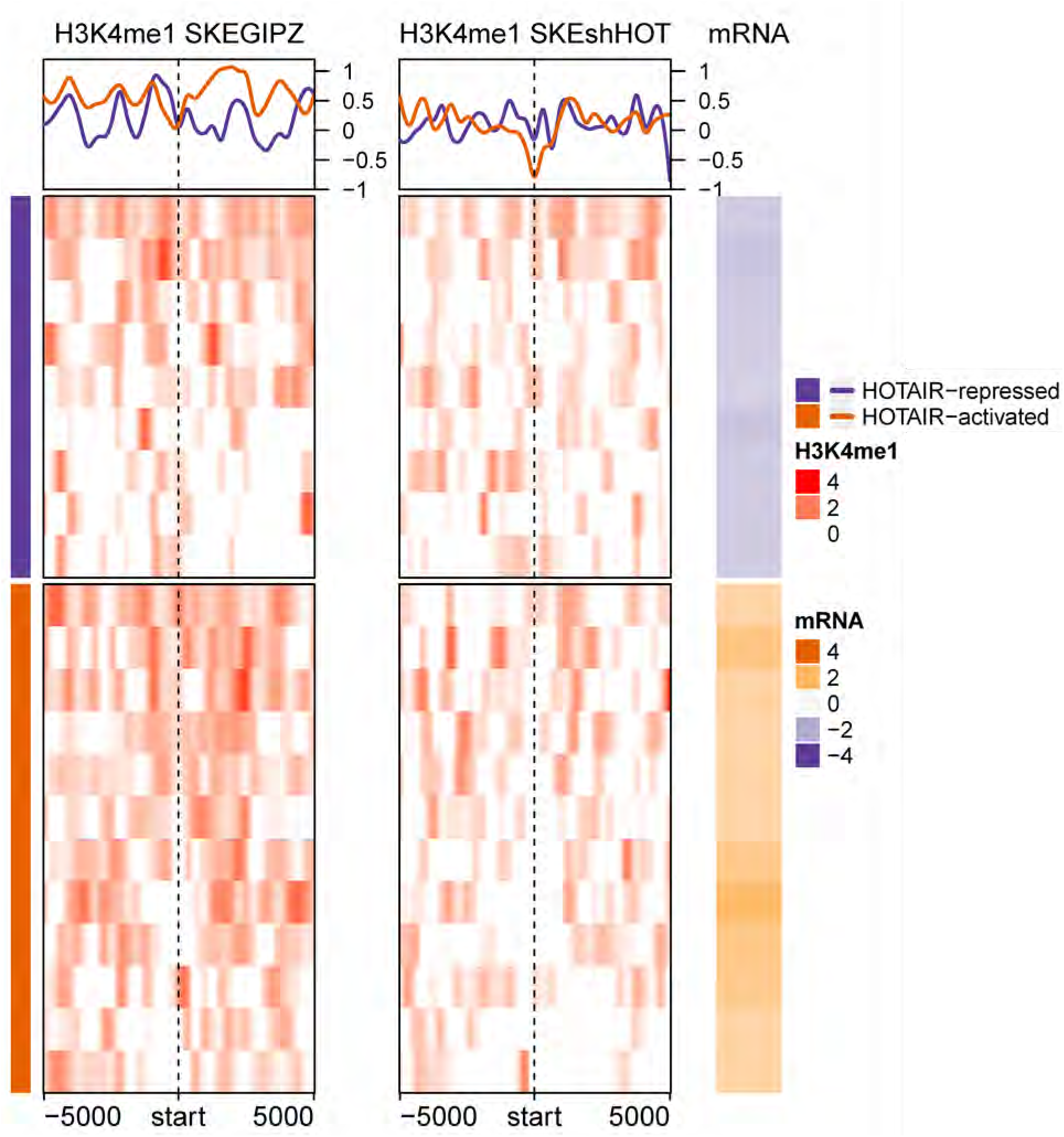
Profiles of HOTAIR-induced H3K4me1 modification around TSS of HOTAIR-regulated genes (21 genes in venn diagram on Figure 13). Each row corresponds to HOTAIR-regulated genes (9 down- and 12 up-regulated genes). Color for H3K4me1 profile represents log_2_ ratio of ChIP to input control. Positive (red) represents enrichment of H3K4me1. Mean H3K4me1 profile in each cell line are plotted on top. The corresponding mRNA expression (log2 ratio) is shown on the right panel. Negative (purple) represents decreased expression in SKESGIPZ compared to SKESshHOTAIR (i.e. genes that are repressed by HOTAIR). “hm_clustSKEGIPZenrich_H3K4M1_vsSKEshHOT.pdf”

### H3K4me2 modulation in SKES cell line

2,646 (adjacent to 2,090 genes) and 19,586 (adjacent to 10,265 genes) H3K4me2 peaks were identified at FDR cut-off of 0.05 using MACS2 in ES2GIPZ and ES2shHOTAIR, respectively. Peaks in each cell line were normalized to their corresponding input. 416 and 657 HOTAIR-regulated genes are found adjacent to H3K4me2 peaks in ES2GIPZ and ES2shHOTAIR, respectively.

**Figure 34.**
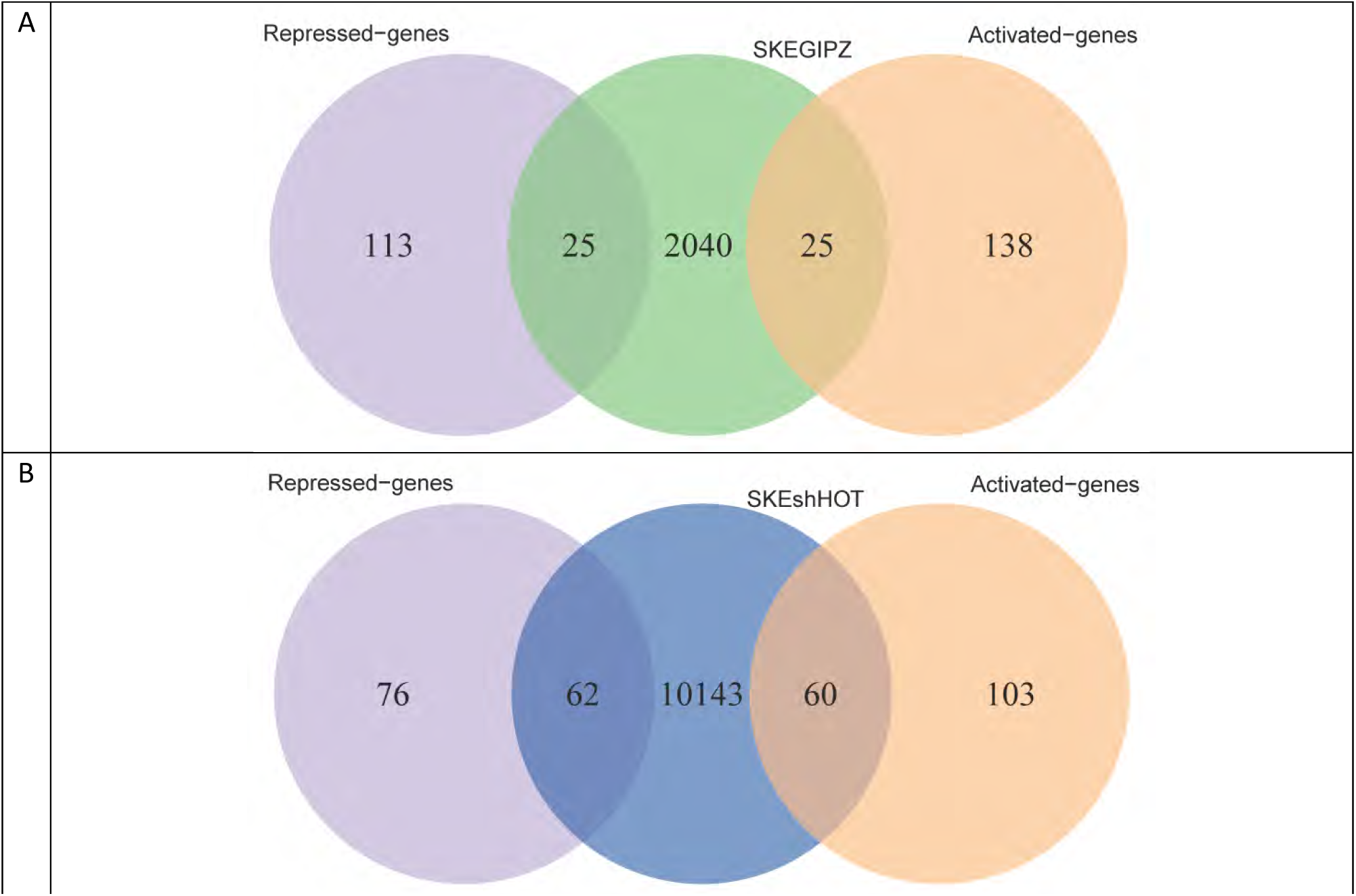
Overlap between genes adjacent to H3K4me2 peaks in (A) SKESGIPZ and (B) SKESshHOT with the HOTAIR-regulated genes.

Globally, HOTAIR expression induces H3K4me2 modification at TSS of HOTAIR-regulated genes and the broad region surrounding the TSS in SKES cell line.

**Figure 35.**
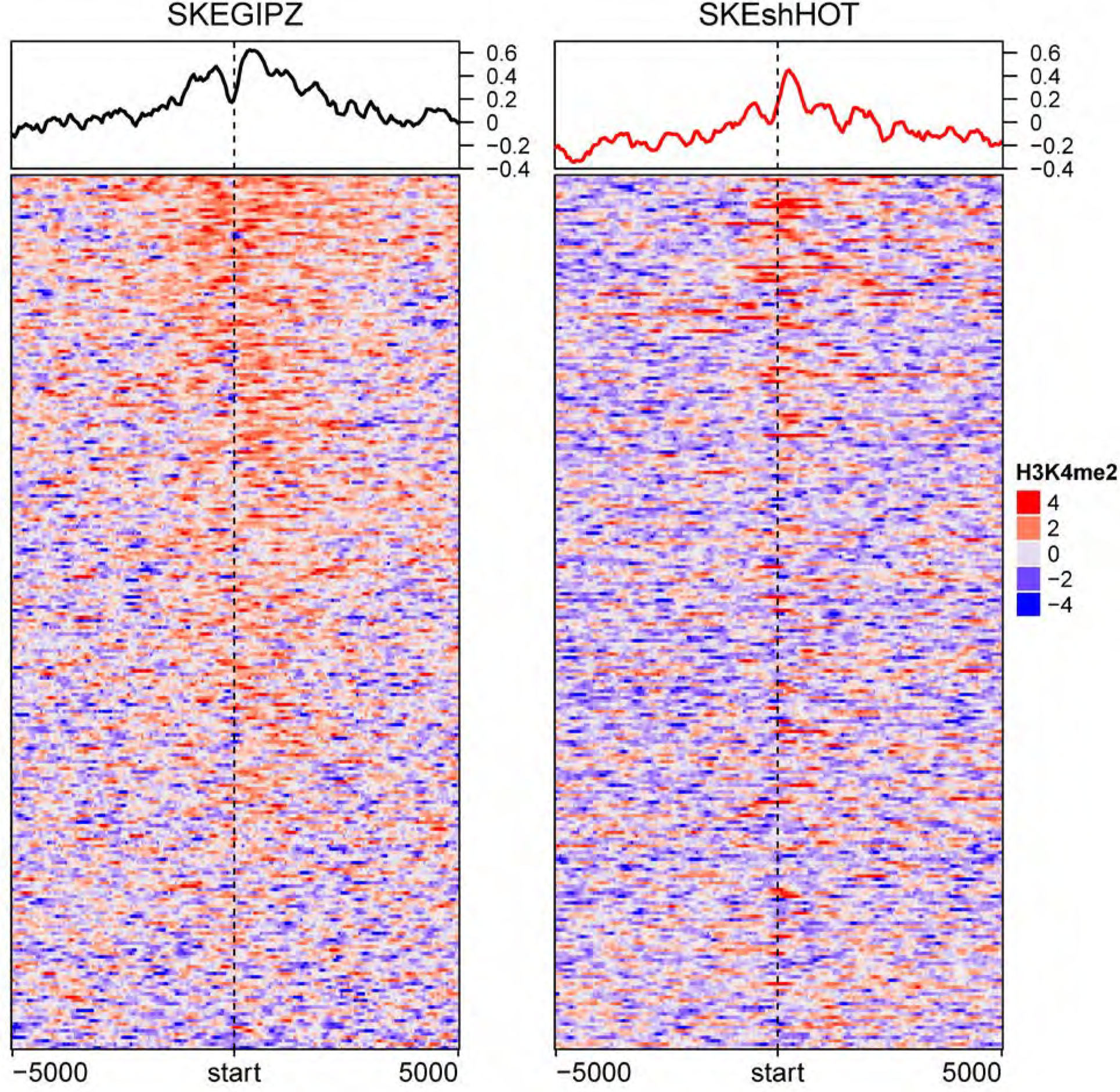
HOTAIR expression induces decreased H3K4me2 binding around TSS of HOTAIR-regulated genes. Visualization of H3K4me2 profiles in SKESGIPZ (left) and SKESshHOTAIR (right) around ± 5000-bp of HOTAIR-regulated genes. Color represents log_2_ ratio of normalized read counts over input. Mean signals in each cell line are plotted on top. Each row represents HOTAIR-regulated gene. hm_SKEshHOT_SKEGIPZ_H3K4M2_norm_diff.pdf

Profiles of H3K4me2 binding around non-differentially expressed genes are shown in “hm_SKEshHOT_SKEGIPZ_H3K4M2_norm_nodiff.pdf”

### H3K4me2- induced and -repressed binding by HOTAIR expression

We found 1,197 H3K4me2 bindings that are induced by HOTAIR and 17,969 H3K4me2 bindings that are repressed by HOTAIR. HOTAIR-induced H3K4me2 binding are adjacent to 22 HOTAIR-regulated genes, while HOTAIR-repressed H3K4me2 binding are adjacent to 115 HOTAIR-regulated genes. H3K4me2 bindings adjacent to only 4 HOTAIR-regulated gene are exclusively induced by HOTAIR. On the other hand, 97 genes have H3K4me2 bindings that are exclusively repressed by HOTAIR.

#### Methods

1,197 regions with significant increase of H3K4me2 binding (HOTAIR-induced binding) in SKESGIPZ compared to SKESshHOTAIR were identified with FDR ≤ 0.05 using MACS2 diff. 17,969 regions with significant increase of H3K4me2 binding (HOTAIR-repressed) were identified in SKESshHOTAIR compared to SKESGIPZ.

**Figure 36.**
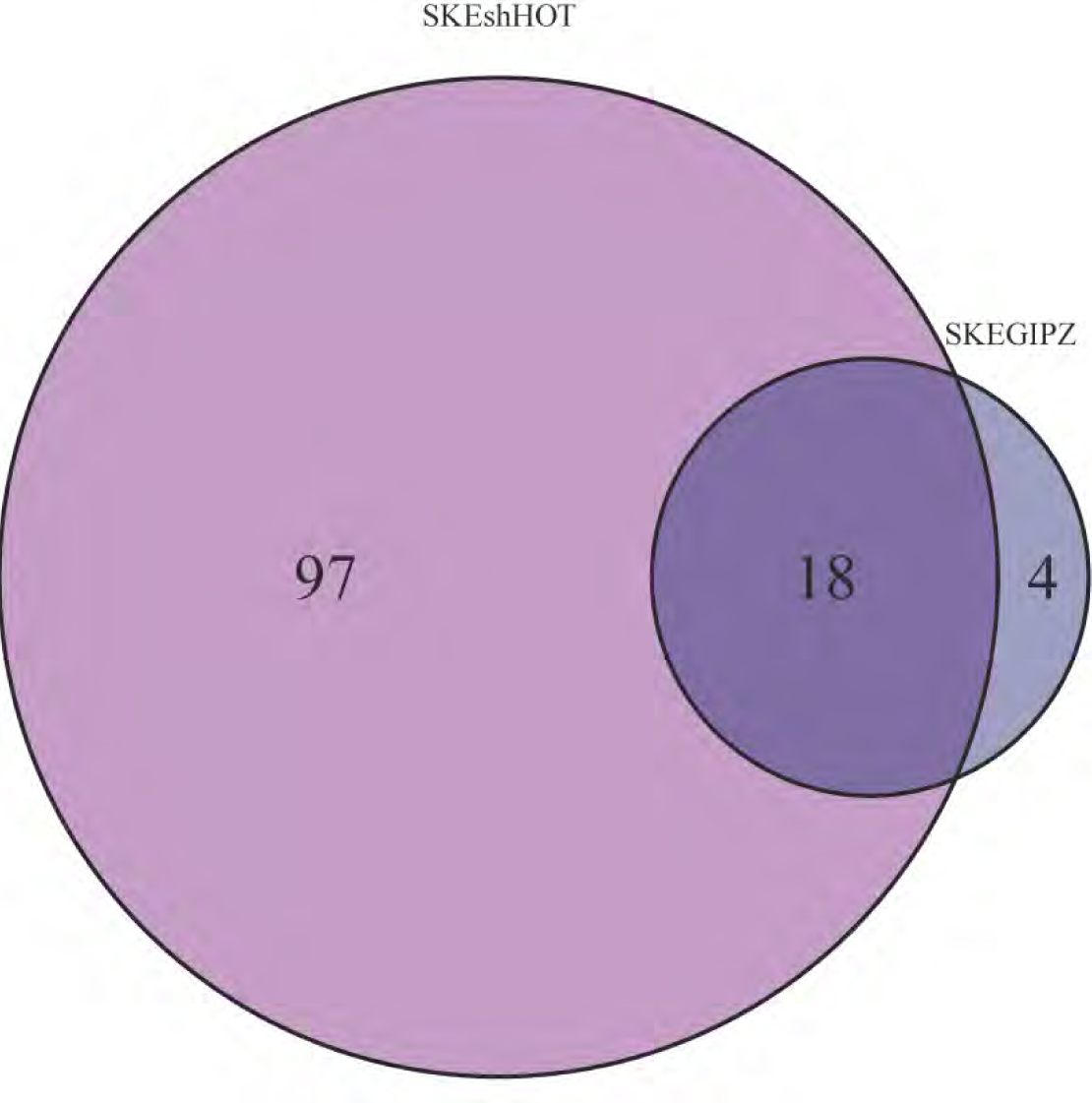
Overlap between HOTAIR-induced and -repressed H3K4me2 bindings. Venn diagram shows overlap between numbers of HOTAIR-regulated genes adjacent to increased H3K4me2 binding in SKESGIPZ (dark blue) and increased H3K4me2 binding in SKESshHOTAIR (magenta). Note that multiple differential binding may be adjacent to the same gene. venn_SKEGIPZvsSKEshHOT_H3K4M2.pdf.

**Figure 37.**
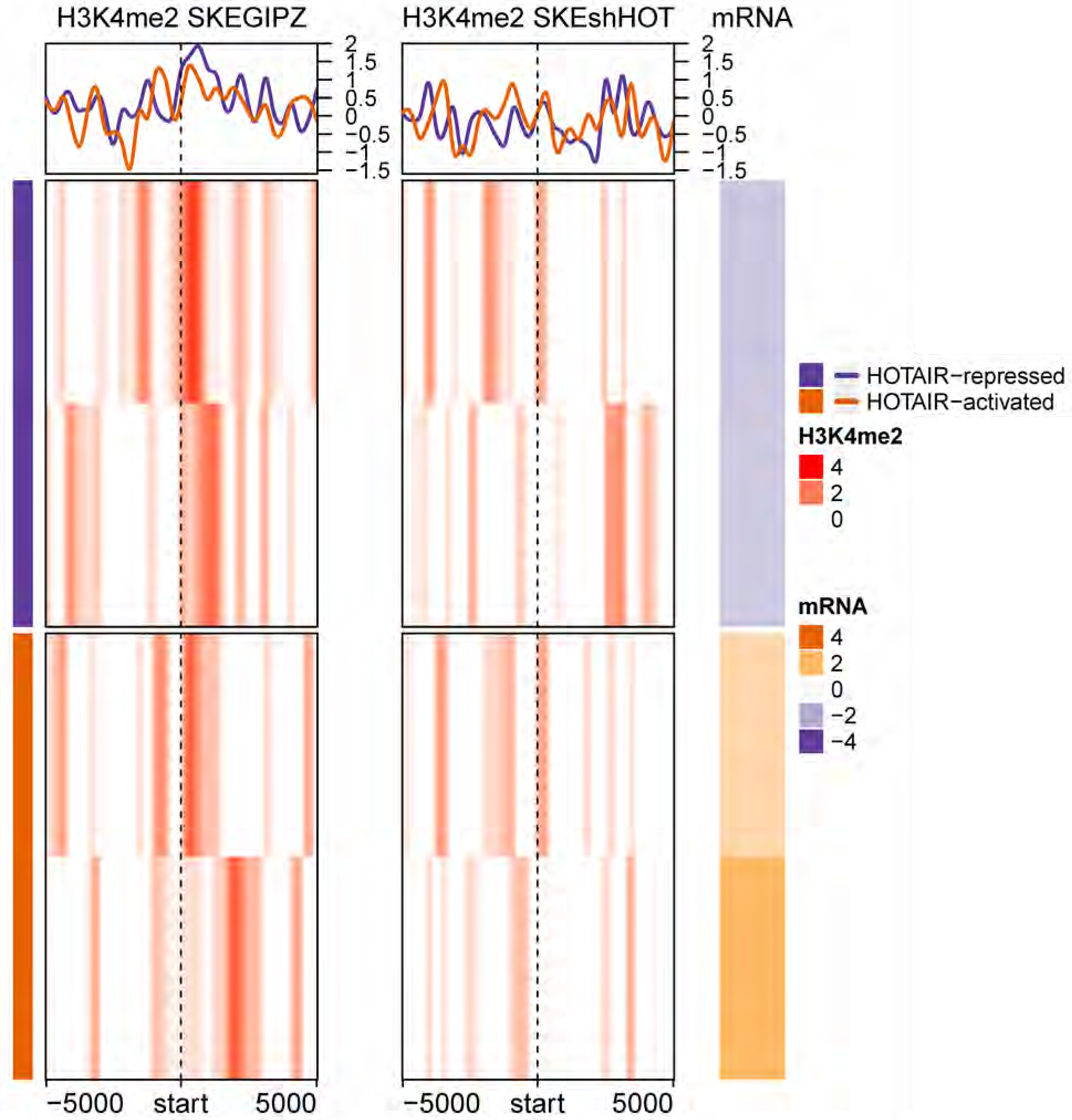
Profiles of HOTAIR-induced H3K4me2 bindings in SKESGIPZ and SKESshHOTAIR around TSS of HOTAIR-regulated genes (4 genes in venn diagram on Figure 13). Each row corresponds to HOTAIR-regulated genes (2 down- and 2 up-regulated genes). Color for H3K4me2 profile represents log_2_ ratio of ChIP to input control. Positive (red) represents enrichment of H3K4me2. Mean H3K4me2 profile in each cell line are plotted on top. The corresponding mRNA expression (log2 ratio) is shown on the right panel. Negative (purple) represents decreased expression in SKESGIPZ compared to SKESshHOTAIR (i.e. genes that are repressed by HOTAIR). “hm_clustSKEGIPZenrich_H3K4M2_vsSKEshHOT.pdf”

**Figure 38.**
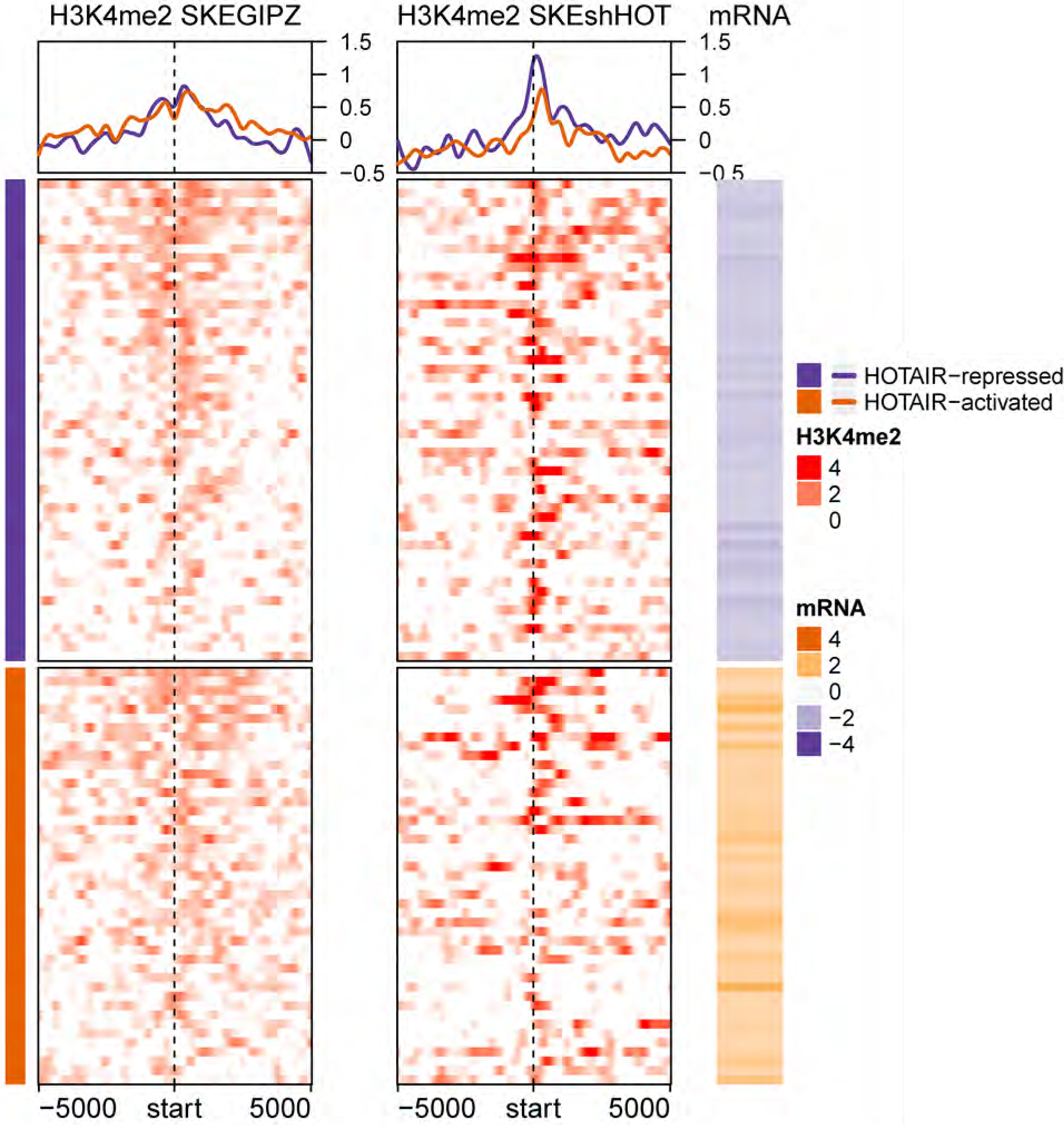
Profiles of HOTAIR-repressed H3K4me2 bindings around TSS of HOTAIR-regulated genes (97 genes in venn diagram on Figure 36). Each row corresponds to HOTAIR-regulated genes (52 down- and 45 up-regulated genes). Color for H3K4me2 profile represents log_2_ ratio of ChIP to input control. Positive (red) represents enrichment of H3K4me2. Mean H3K4me2 profile in each cell line are plotted on top. The corresponding mRNA expression (log2 ratio) is shown on the right panel. Negative (purple) represents decreased expression in SKESGIPZ compared to SKESshHOTAIR (i.e. genes that are repressed by HOTAIR). “hm_clustSKEshHOTenrich_H3K4M2_vsSKEGIPZ.pdf”

## SKES Cell Lines with different threshold for HOTAIR-regulated genes

### H3K4me1 enrichment in SKES in relation to HOTAIR expression

67,906 (adjacent to 16,872 genes) and 359 (adjacent to 243 genes) H3K4me1 broad peaks were identified at FDR ≤ 0.05 using MACS2 in SKESGIPZ and SKESshHOTAIR, respectively. Peaks in each cell line were normalized to their corresponding input. For SKES cell lines, we changed the threshold to define differentially expressed genes. 1,668 genes were found to be differentially regulated by HOTAIR (FDR ≤ 0.01 and absolute fold-change ≥ 1.5). (Note: the number of differential genes is 1,852 if we used FDR ≤ 0.05). Genes regulated by HOTAIR have positive fold-change. 1,039 and 15 HOTAIR-regulated genes are found adjacent to H3K4me1 peaks in SKESGIPZ and SKESshHOTAIR, respectively.

**Figure 30.**
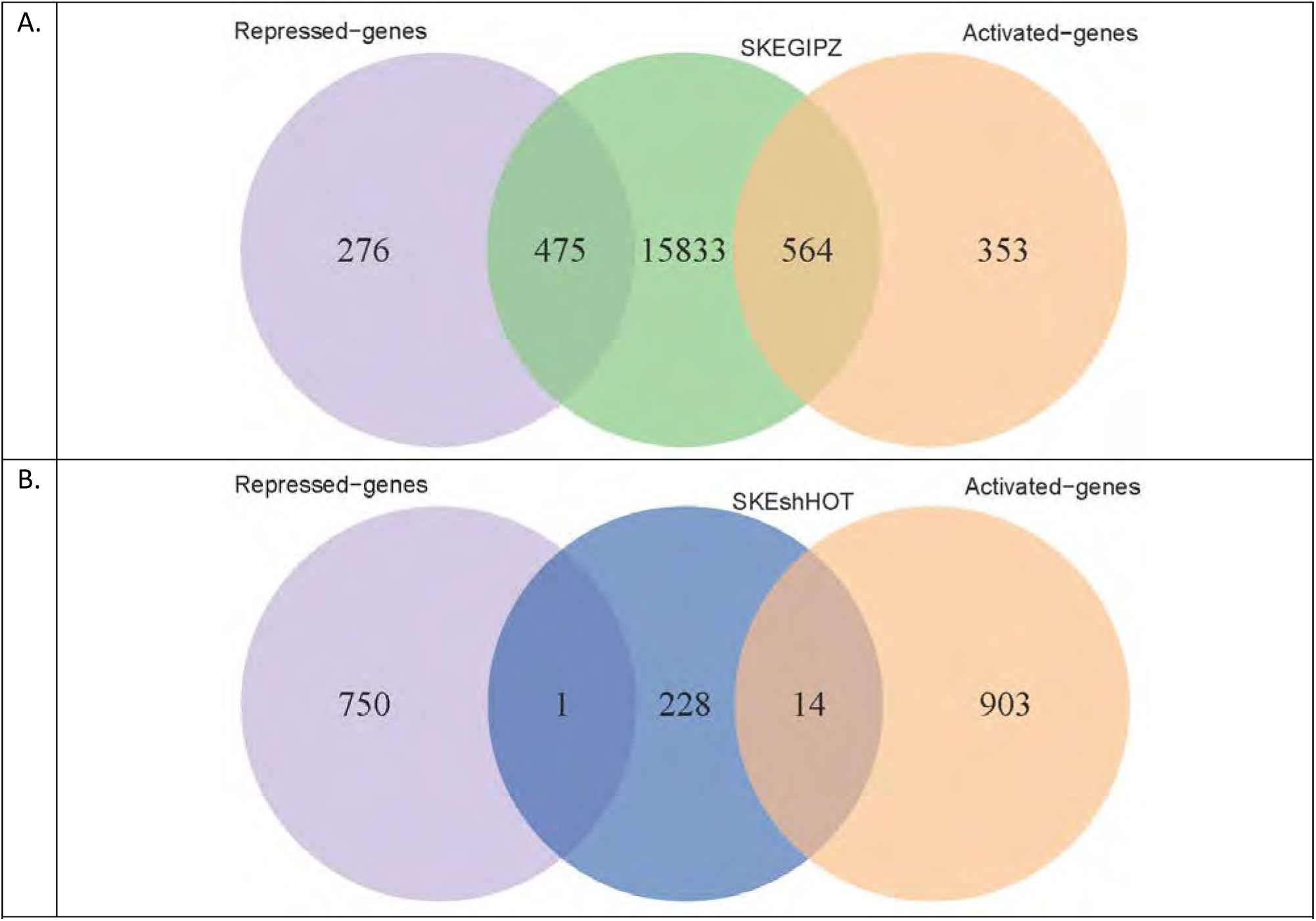
Overlap between genes adjacent to H3K4me1 peaks in (A) SKESGIPZ and (B) SKESshHOT with the HOTAIR-regulated genes. venn_SKEshHOT_H3K4M1_diff_genes_2.pdf and venn_SKEGIPZ_H3K4M1_diff_genes_2.pdf

In SKES cells, HOTAIR expression seems to induce increased H3K4me1 in broad regions surrounding TSS of HOTAIR-regulated genes. Without HOTAIR expression, there is depletion of H3K4me1 binding at their TSSs.

**Figure 31.**
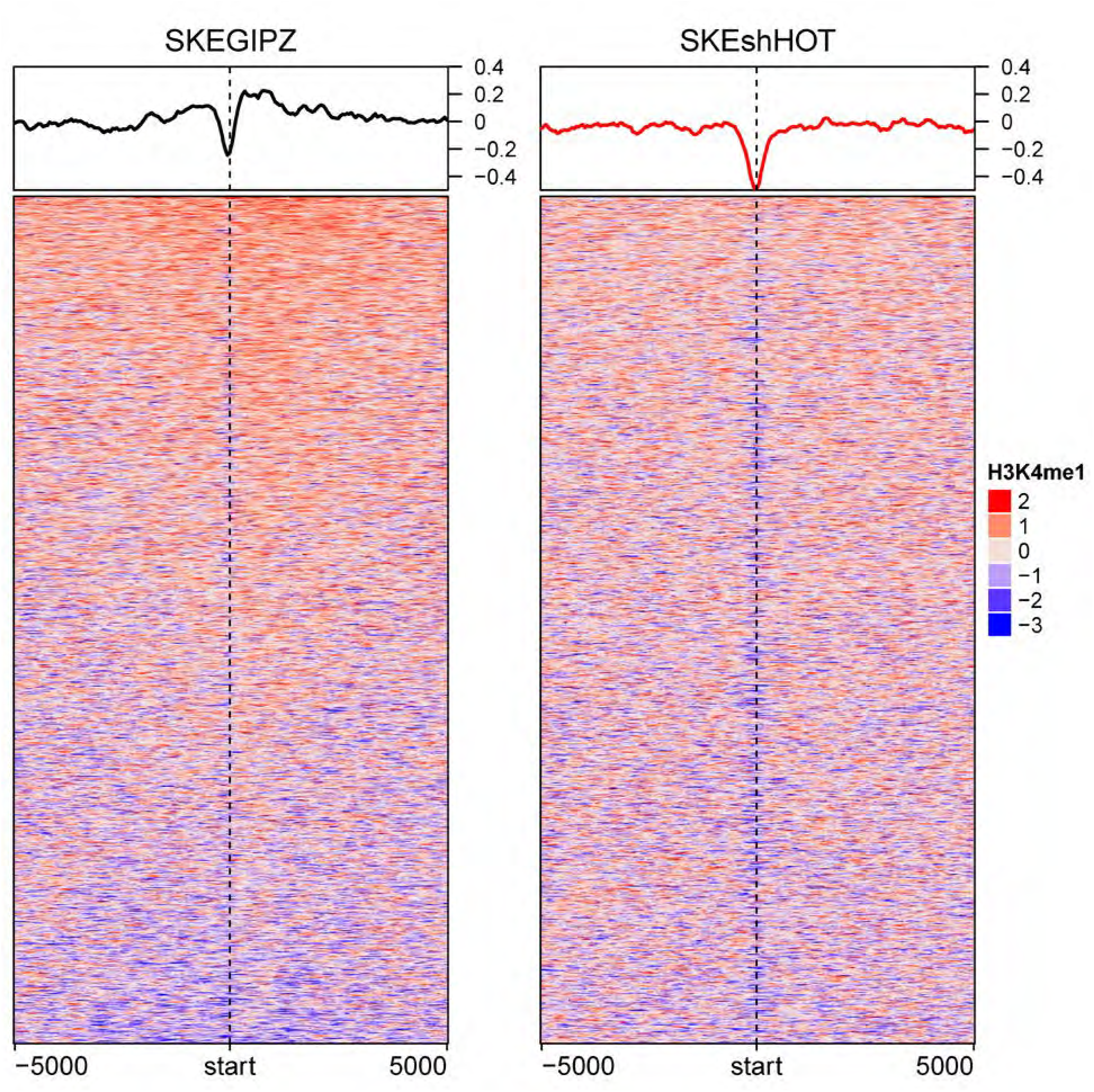
HOTAIR expression induces H3K4me1 binding around TSS of HOTAIR-regulated genes. Visualization of H3K4me1 profiles in SKESGIPZ (left) and SKESshHOTAIR (right) around ± 5000-bp of HOTAIR-regulated genes in ES2. Color represents log_2_ ratio of normalized read counts over input. Mean signals in each cell line are plotted on top. Each row represents HOTAIR-regulated gene. hm_SKEshHOT_SKEGIPZ_H3K4M1_norm_diff.pdf

### H3K4me1- induced and -repressed binding by HOTAIR expression

We found 1,757 H3K4me1 bindings that are induced by HOTAIR and 51 H3K4me1 bindings that are repressed by HOTAIR. HOTAIR-induced H3K4me1 binding are adjacent to 94 HOTAIR-regulated genes, while there are only 5 HOTAIR-regulated genes near HOTAIR-repressed H3K4me1 binding.

#### Methods

**Figure 32.**
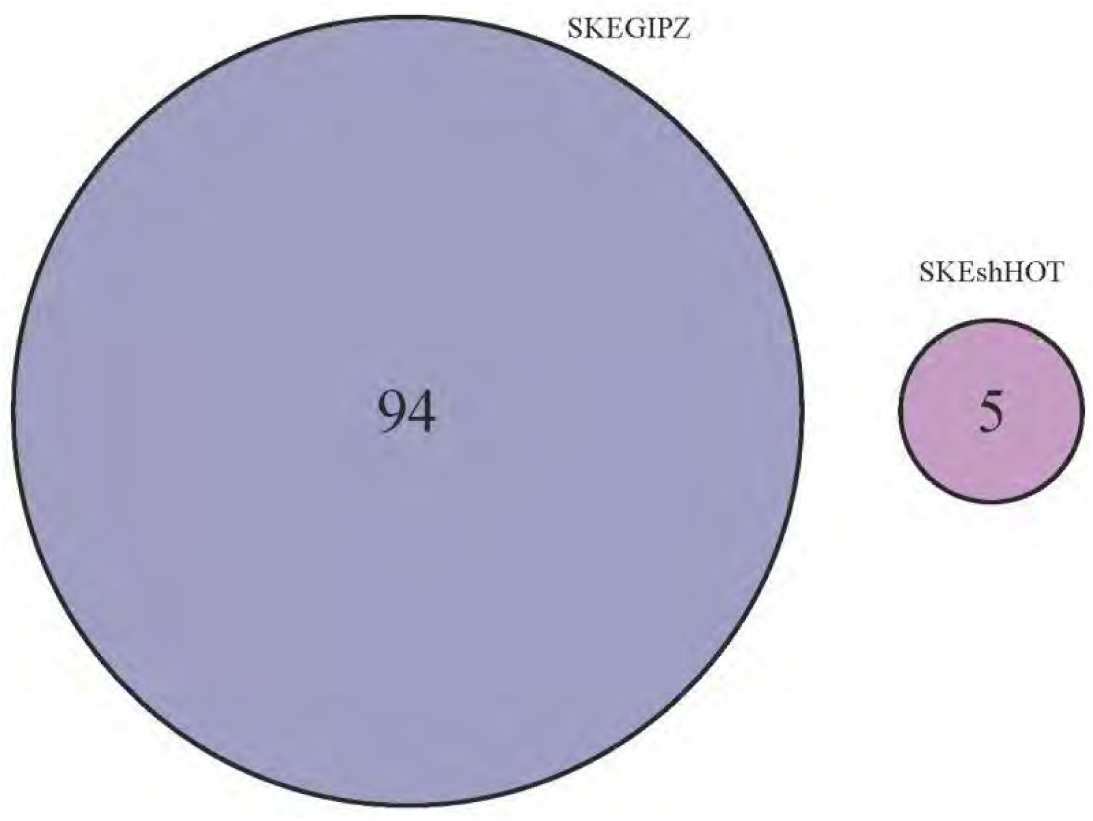
Venn diagram shows the number of HOTAIR-regulated genes adjacent to increased H3K4me1 binding in SKESGIPZ (dark blue) and increased H3K4me1 binding in SKESshHOTAIR (magenta). Note that multiple differential binding may be adjacent to the same gene. venn_SKEGIPZvsSKEshHOT_H3K4M1_2.pdf.

**Figure 33.**
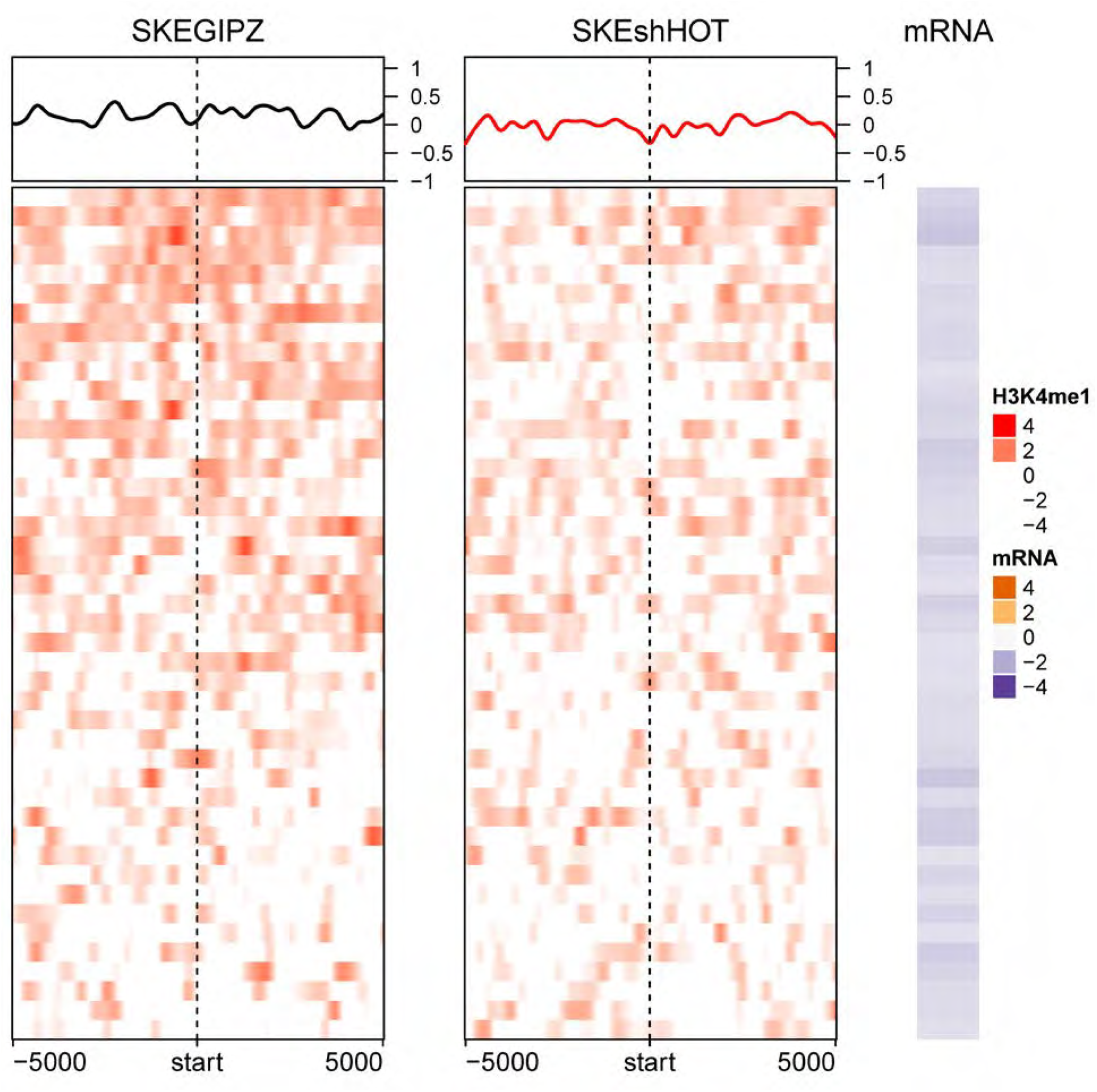
Profiles of HOTAIR-induced H3K4me1 modification around TSS of HOTAIR-repressed genes. Each row corresponds to HOTAIR-regulated genes (44 downregulated genes). Color for H3K4me1 profile represents log_2_ ratio of ChIP to input control. Positive (red) represents enrichment of H3K4me1. Mean H3K4me1 profile in each cell line are plotted on top. The corresponding mRNA expression (log2 ratio) is shown on the right panel. Negative (purple) represents decreased expression in SKESGIPZ compared to SKESshHOTAIR (i.e. genes that are repressed by HOTAIR). “hm_clustSKEGIPZenrich_H3K4M1_vsSKEshHOT_rep_genes_2.pdf”

### H3K4me2 modulation in SKES cell line

2,646 (adjacent to 2,090 genes) and 19,586 (adjacent to 10,265 genes) H3K4me2 peaks were identified at FDR cut-off of 0.05 using MACS2 in ES2GIPZ and ES2shHOTAIR, respectively. Peaks in each cell line were normalized to their corresponding input. 205 and 657 HOTAIR-regulated genes are found adjacent to H3K4me2 peaks in ES2GIPZ and ES2shHOTAIR, respectively.

**Figure 34.**
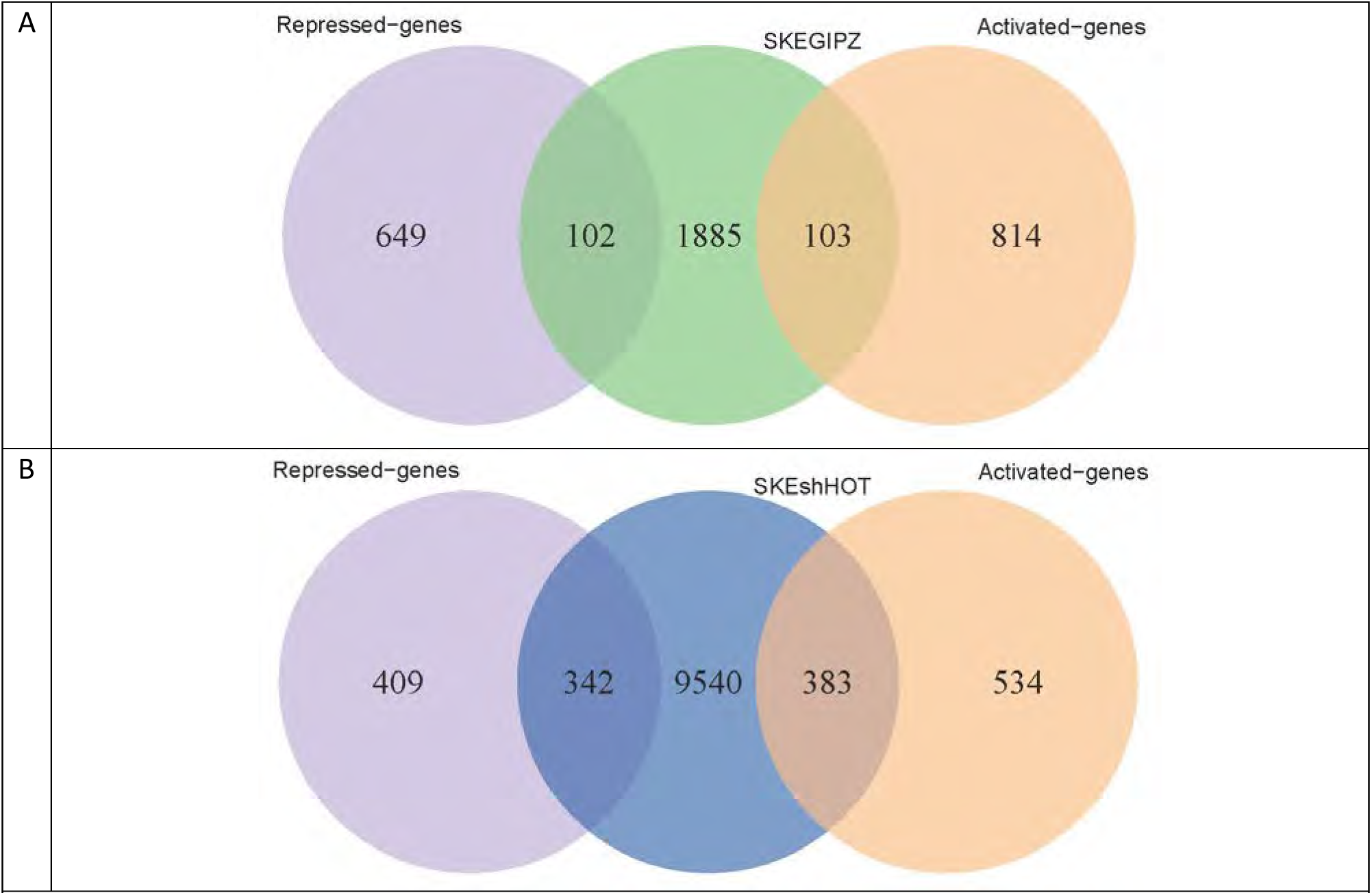
Overlap between genes adjacent to H3K4me2 peaks in (A) SKESGIPZ and (B) SKESshHOT with the HOTAIR-regulated genes.

Globally, HOTAIR expression induces H3K4me2 modification mainly in the broad region surrounding the TSS in SKES cell line, however not at the TSSs.

**Figure 35.**
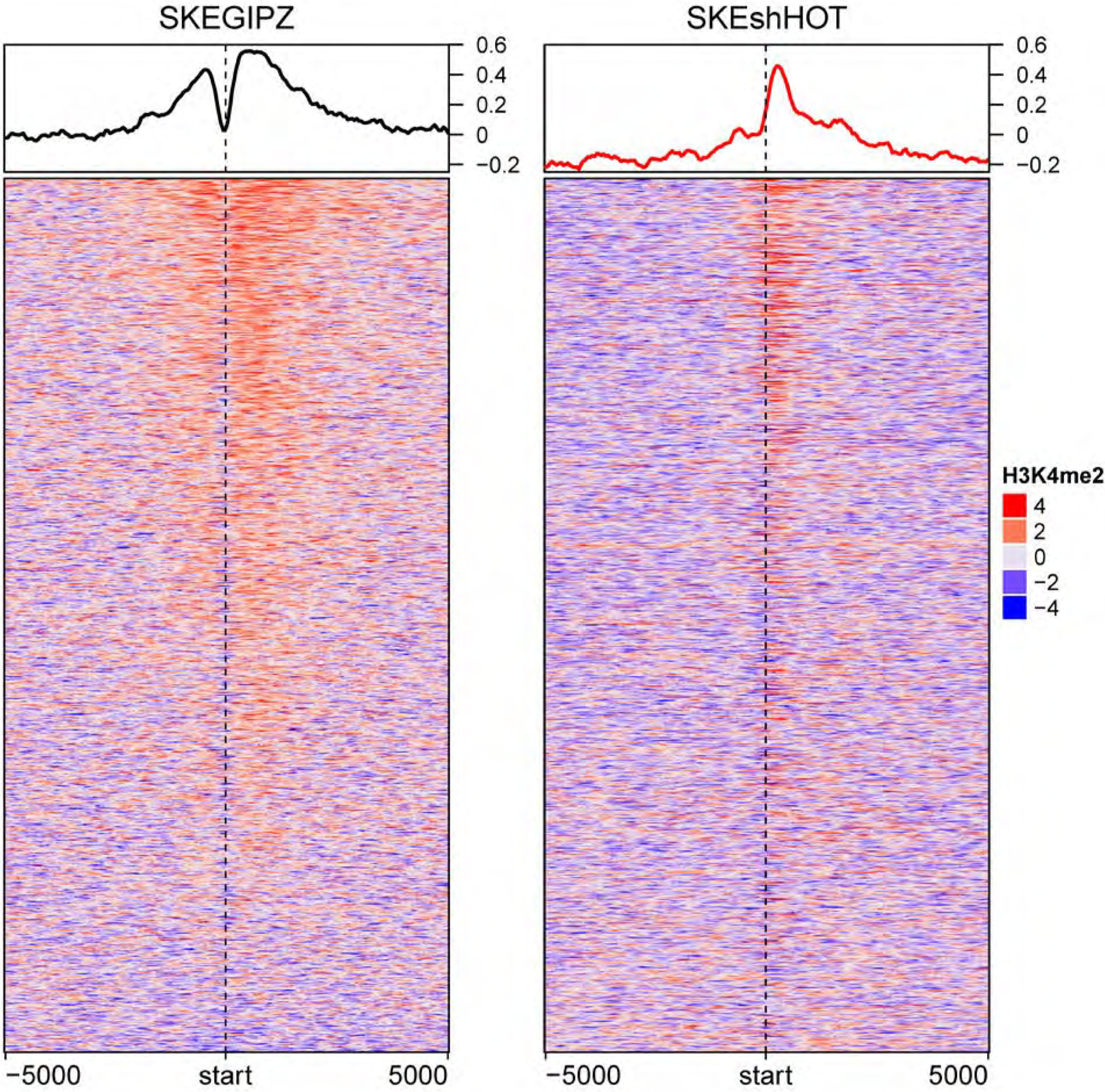
HOTAIR expression induces decreased H3K4me2 binding around TSS of HOTAIR-regulated genes. Visualization of H3K4me2 profiles in SKESGIPZ (left) and SKESshHOTAIR (right) around ± 5000-bp of HOTAIR-regulated genes. Color represents log_2_ ratio of normalized read counts over input. Mean signals in each cell line are plotted on top. Each row represents HOTAIR-regulated gene. hm_SKEshHOT_SKEGIPZ_H3K4M2_norm_diff.pdf

Profiles of H3K4me2 binding around non-differentially expressed genes are shown in “hm_SKEshHOT_SKEGIPZ_H3K4M2_norm_nodiff_2.pdf”

### H3K4me2- induced and -repressed binding by HOTAIR expression

We found 1,197 H3K4me2 bindings that are induced by HOTAIR and 17,969 H3K4me2 bindings that are repressed by HOTAIR. HOTAIR-induced H3K4me2 binding are adjacent to 91 HOTAIR-regulated genes, while HOTAIR-repressed H3K4me2 binding are adjacent to 679 HOTAIR-regulated genes. H3K4me2 bindings adjacent to 29 HOTAIR-regulated gene are exclusively induced by HOTAIR. On the other hand, 617 genes have H3K4me2 bindings that are exclusively repressed by HOTAIR.

#### Methods

**Figure 36.**
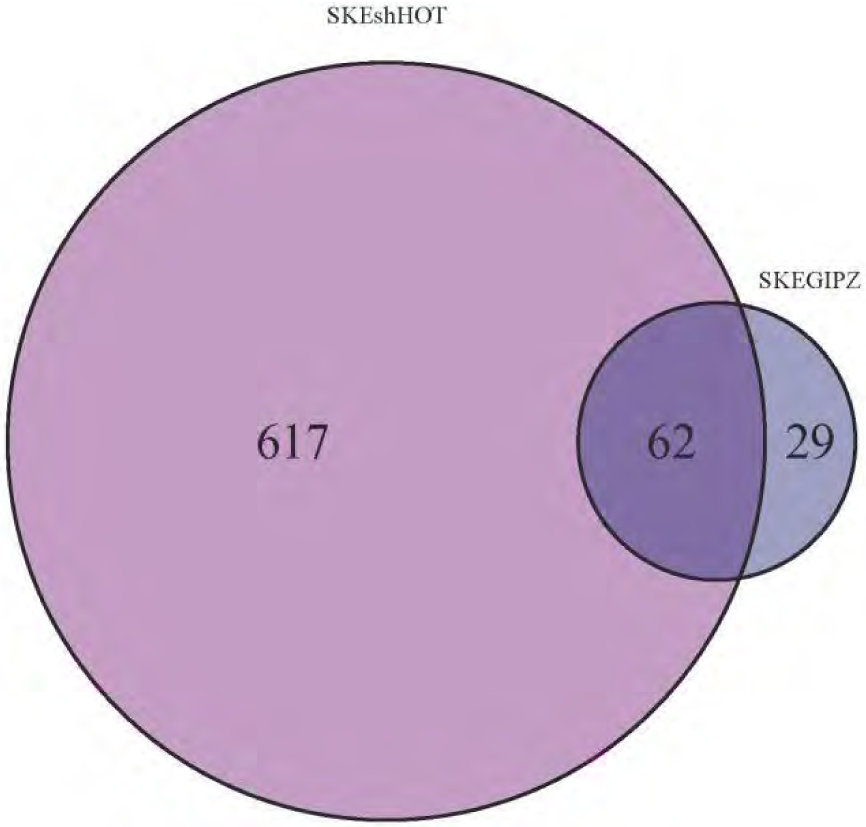
Overlap between HOTAIR-induced and -repressed H3K4me2 bindings. Venn diagram shows overlap between numbers of HOTAIR-regulated genes adjacent to increased H3K4me2 binding in SKESGIPZ (dark blue) and increased H3K4me2 binding in SKESshHOTAIR (magenta). Note that multiple differential binding may be adjacent to the same gene. venn_SKEGIPZvsSKEshHOT_H3K4M2_2.pdf.

**Figure 37.**
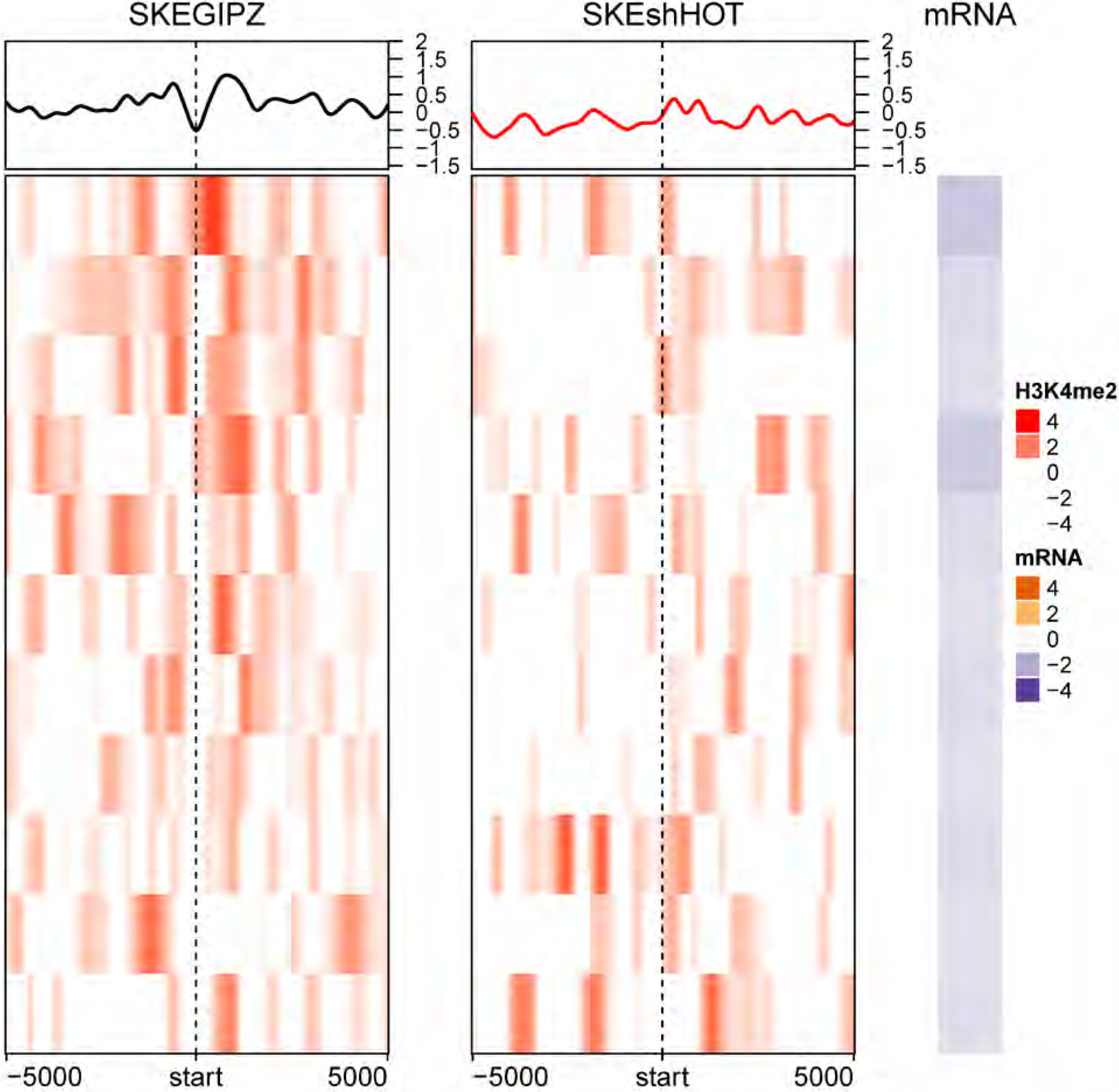
Profiles of HOTAIR-induced H3K4me2 bindings in SKESGIPZ and SKESshHOTAIR around TSS of HOTAIR-repressed genes. Each row corresponds to HOTAIR-repressed genes (11 down-regulated genes). Color for H3K4me2 profile represents log_2_ ratio of ChIP to input control. Positive (red) represents enrichment of H3K4me2. Mean H3K4me2 profile in each cell line are plotted on top. The corresponding mRNA expression (log2 ratio) is shown on the right panel. Negative (purple) represents decreased expression in SKESGIPZ compared to SKESshHOTAIR (i.e. genes that are repressed by HOTAIR). “hm_SKEGIPZenrich_H3K4M2_vsSKEshHOT_rep_genes_2.pdf”

**Figure 38.**
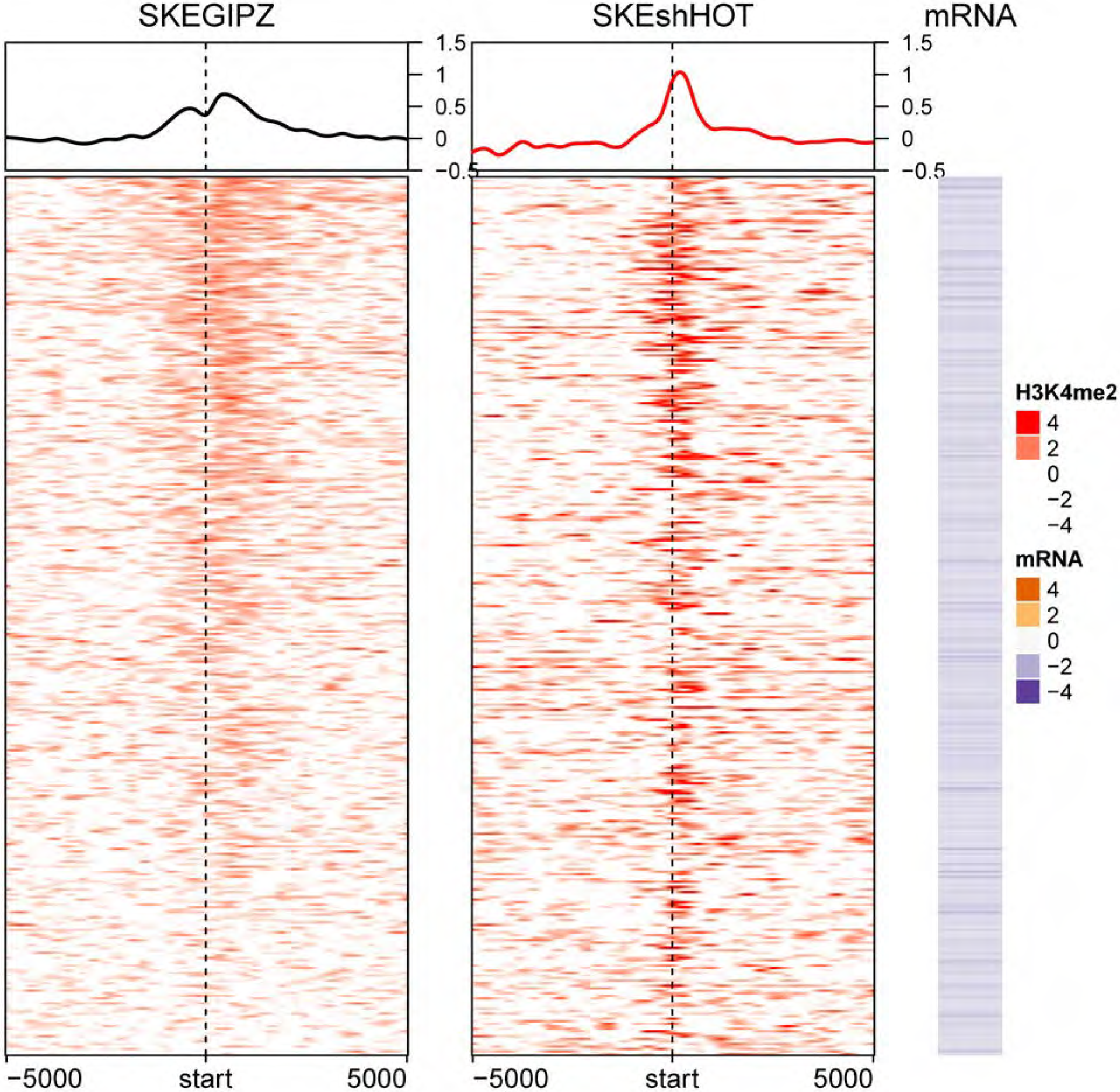
Profiles of HOTAIR-repressed H3K4me2 bindings around TSS of HOTAIR-repressed genes. Each row corresponds to HOTAIR-regulated genes (347 down-regulated genes). Color for H3K4me2 profile represents log_2_ ratio of ChIP to input control. Positive (red) represents enrichment of H3K4me2. Mean H3K4me2 profile in each cell line are plotted on top. The corresponding mRNA expression (log2 ratio) is shown on the right panel. Negative (purple) represents decreased expression in SKESGIPZ compared to SKESshHOTAIR (i.e. genes that are repressed by HOTAIR). “hm_SKEshHOTenrich_H3K4M2_vsSKEshHOT_rep_genes_2.pdf”

#### File lists

_peaks.xls contains ChIP-seq peaks location output files from MACS2

diff_<cell1><chip>_vs_<cell2><chip>_c3.0_common.bed contains ChIP-seq peaks locations (output from MACS2 diff) that are increased in cell1 compared to cell2

diff_<cell1><chip>_vs_<cell2><chip>_c3.0_cond1.bed contains ChIP-seq peaks locations (output from MACS2 diff) that are significantly increased in cell1 compared to cell2

diff_<cell1><chip>_vs_<cell2><chip>_c3.0_cond2.bed contains ChIP-seq peaks locations (output from MACS2 diff) that are significantly increased in cell2 compared to cell1

_peaks_annot.txt contains ChIP-seq peaks annotated with nearest genes

<cell1>enrich_<chip>_vs<cell2>.txt contains ChIP-seq peaks with chip pull down that are significantly higher in cell1 when compared to cell2 annotated with nearest genes

SI10 (Excel workbook): Gene expression datasets, including are differentially-expressed gene lists and quality-control files for RNA-Seq analyses described in the manuscript, and Figures 5A and B with gene names

